# Evaluation of 6 MALDI-Matrices for 10 µm lipid imaging and on-tissue MSn with AP-MALDI-Orbitrap

**DOI:** 10.1101/2021.10.27.466111

**Authors:** Tina B. Angerer, Jerome Bour, Jean-Luc Biagi, Eugene Moskovets, Gilles Frache

## Abstract

Mass spectrometry imaging (MSI) is a technique uniquely suited to localize and identify lipids in a tissue sample. Using an AP-MALDI UHR source coupled to an Orbitrap Elite, numerous lipid locations and structures can be determined in high mass resolution spectra and at cellular spatial resolution, but careful sample preparation is necessary. We tested 11 protocols on serial brain sections for the commonly used MALDI matrices, CHCA, Norharmane, DHB, DHAP, THAP, and DAN, in combination with tissue washing and matrix additives, to determine the lipid coverage, signal intensity, and spatial resolution achievable with AP-MALDI. In positive ion mode, the most lipids could be detected with CHCA and THAP, while THAP and DAN without additional treatment offered the best signal intensities. In negative ion mode, DAN showed the best lipid coverage and DHAP performed superior for Gangliosides. DHB produced intense cholesterol signals in the white matter. 155 lipids were assigned in positive (THAP), 137 in negative ion mode (DAN) and 76 lipids were identified using on tissue tandem-MS. The spatial resolution achievable with DAN was 10 μm, confirmed with on tissue line-scans. This enabled the association of lipid species to single neurons in AP-MALDI images. The results show that the performance of AP-MALDI is comparable to vacuum MALDI techniques for lipid imaging.

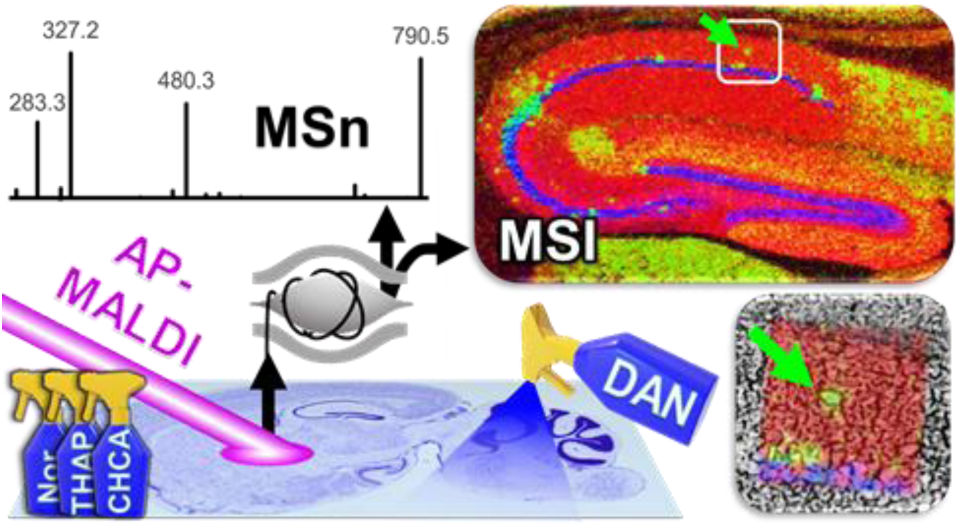

## Introduction

Mass spectrometry imaging (MSI) is a technique capable of locating and identifying atoms and molecules in a sample. By scanning across a surface and recording individual mass spectra at each location, MSI generates “chemical maps” which display the distribution of all detected species on a sample surface. MSI was first performed in 1949^1^ but it has only gained traction in the last 20 years. With an ever-growing number of techniques emerging, MSI is now regularly applied in a variety of fields, ranging from inorganic materials science to bio-medical research. Especially for biologists MSI is of great interest, due to the fact that it is now possible to analyze intact macromolecules (e.g. proteins, lipids and neurotransmitter)^2^ at cellular resolutions^3^ while small molecules and atoms can be detected in cell organelles.^4^ Multimodal experiments connect proteomic and lipidomic data and deepen our understanding of biological processes.^5-7^ The capabilities, applications and drawbacks of MSI techniques are described in several reviews.^8-12^

Lipidomics is of ever growing importance in the medical field.^13^ The changes in lipid compositions due to the onset of disease have been recognized, but traditional techniques, such as liquid chromatography mass spectrometry (LC-MS) fail to capture the complexity of heterogamous samples (e.g. tumors). In contrast to proteins, single lipids cannot be labelled easily. MSI is uniquely suited to localize individual lipid species in a sample, as it distinguishes lipids based on their accurate mass. This improves upon unspecific dyes and techniques requiring sample homogenization (e.g. LC) but is insufficient to determine the exact species, since a number of different lipids can be present within a narrow mass range and lipids can have isomers, even from different classes. For this reason, an increasing number of MSI devices offer high resolution mass analyzers and tandem-MS capability for structural identification to enhance specificity.^14^ With these capabilities, MSI has significantly contributed to the field of lipidomics by revealing lipid alterations in various diseases.^15-18^

Matrix assisted laser desorption ionization (MALDI)-MSI is a technique where a matrix is applied to a sample surface, molecules of interest are extracted, embedded in matrix crystals, desorbed with a laser beam and ionized, and finally detected with a mass analyzer. The detectable species largely depend on the applied matrix, of which there are numerous options. In general, a MALDI matrix must be able to absorb the laser light and transfer charges to/from the target molecules, usually be vacuum stable, and, in the case of MSI, form (sub)micron sized crystals to enable highly localized detection. The achievable spatial resolution depends on the target molecules, the matrix properties, and the laser beam focus. For lipids 5-10 μm has been demonstrated, and proteins are usually imaged at 50-100 μm (better spatial resolutions for lipids and proteins have been demonstrated with e.g. t-MALDI-2).^15, 19-22^

MALDI performed under ambient conditions (atmospheric pressure (AP)-MALDI) removes the requirement of a vacuum stable matrix and enables the use of more volatile substances, such as 2′,6′-Dihydroxyacetophenone (DHAP) and 2,4,6-Trihydroxyacetophenone (THAP). Woods et al state that the addition of heptafluorobutyric acid (HFBA) stabilizes DHAP in vacuum but recent reports show its sublimation, even with added HFBA.^23-24^ An added benefit of AP-MALDI is that the tissue sample is not subjected to the harsh vacuum conditions leading to drying and cracking, which makes subsequent procedures such as histological tissue staining more likely to succeed.^25-29^ The downside is the shortened mean free path for the generated ions, possibly leading to their neutralization and diminishing sensitivity. This can reduce the useful spatial resolution and the ability to perform tandem-MS. Previous reports investigated AP-MALDI capabilities^30^ and have compared the performance of several matrices for AP-MALDI imaging^24, 31-35^ but, to our knowledge, there is currently no comprehensive report, comparing the numerous matrices available and their lipid coverage on the same sample type and device, in positive and negative ion mode.

Therefore, in this study we tested 11 sample preparation protocols with 6 different matrices, washing steps and additives, that have previously been reported to yield good results for high spatial resolution, lipid imaging with vacuum and AP-MALDI techniques. Protocols were adapted for the matrices: α-cyano-4-hydroxycinnamic acid (CHCA),^36-37^ Norharmane (Nor),^36, 38^ 1,5-Diaminonapthalene (DAN),^24, 39^ DHAP,^23-24^ THAP,^40^ and 2’,5’-dihydroxybenzoic acid (DHB),^38, 41^ and their performance was evaluated, in terms of signal intensity/ability to perform tandem-MS, lipid coverage and achievable, useful spatial resolution, in positive and negative ion mode. Additionally, 76 peaks were identified using on tissue tandem-MS.

## Materials and Methods

### Chemicals

Chemicals and solvents (analytical grade) were purchased from the following sources: α-Cyano-4-hydroxycinnamic acid 98% (CHCA) (Sigma Aldrich), 2,5-Dihydroxybenzoic acid 98% (DHB) (Sigma Aldrich), Norharmane 98% (Nor) (Acros Organics), 1,5-Diaminonaphthalene 97% (DAN) (Sigma Aldrich), 2’,4’,6’-Trihydroxyacetophenone 99.5% (THAP) (Sigma Aldrich), 2,6-Dihydroxyacetophenone 99.5% (DHAP) (Sigma Aldrich), acetonitrile (ACN) (Honeywell), chloroform (Acros Organics), methanol (Carl Roth), ammonium acetate (AmAc) (VWR). ammonium sulphate (AmS) (Sigma Aldrich), heptafluorobutyric acid (HFBA) (Sigma Aldrich), and trifluoroacetic acid (TFA) (Sigma Aldrich). All chemicals used in this study were stored, handled, and disposed of according to good laboratory practices (GLP).

### Sample preparation

10 μm sagittal mouse brain sections on indium tin oxide (ITO)-coated glass slides (Diamond Coatings, UK) were prepared at Swansea University, as stated in a recent publication.^42^ Sections were kept at -80ºC until analysis and dried in a vacuum desiccator for 30 minutes prior to matrix application. Optical images of the tissue sections were taken using an Olympus BX51 Microscope (Olympus, Belgium). On some sections, tissue washing was performed with AmAc at 50 mM concentration, chilled to 4°C, for 3×5 seconds, as described previously.^43^ Serial sections on separate ITO glass slides were coated with the various matrices using an HTX TM sprayer (HTX Technologies LLC, USA), flow rate 0.12 ml/min, velocity 1200 mm/min, drying time 2 s, line spacing 2.5 mm. Matrix composition and additives, sprayer temperature, number of matrix layers/passes, and laser settings are listed in table 1. The analyses showing the best performance (most lipids detected, best signal) were repeated on different days, in positive ion mode for CHCA, THAP and DAN70, and in negative ion mode for DAN70.

**Table 1.** Matrix compositions and additives, HTX-TM sprayer settings and AP-MALDI source laser settings. Listed are matrices, matrix concentration (mg/ml), matrix solvents, matrix additives or sample preparation, HTX-TM sprayer temperature, matrix layers/passes (Z), laser frequency (Hz) and intensity (%) for 40/20 and 10 μm imaging, experiments and ion-mode (+/-/±) included for each protocol, and references.

| Matrix (mg/ml) | Solvents | Add./ Sample prep. | C° | Z | Laser (40/20 μm) | Laser (10 μm) | Hip. | Cer. | Chol. | Adapted from: |
| --- | --- | --- | --- | --- | --- | --- | --- | --- | --- | --- |
| CHCA (5) | CHCl<sub>3</sub>:MeOH 1:1 | 0.2% TFA | 40 | 16 | 3000 Hz 10% | 3000 Hz 2.5% | × (+) | × (±) | × (+) | Barré et. al.<sup>36</sup><br>Hochart et. al.<sup>37</sup> |
| Nor (7) | CHCl<sub>3</sub>:MeOH 2:1 |  | 30 | 12 | 500 Hz 10% |  |  | × (±) | × (+) | Barré et. al.<sup>36</sup> |
| DHB (10) | MeOH:H<sub>2</sub>O 7:3 | 0.1% TFA | 50 | 8 | 3000 Hz 10% |  |  | × (+) | × (+) | McMillen et. al.<sup>38</sup><br>Leopold et. al.<sup>41</sup> |
| DHAP (10) | ACN:AmS 7:3 | 125mM (NH<sub>4</sub>)<sub>2</sub>SO<sub>4</sub>, 0.05% HFBA | 50 | 8 | 3000Hz 7.5% |  | × (-)* | × (-) | × (+) | Jackson et. al.<sup>24</sup><br>Colsch et. al.<sup>23</sup> |
| THAP (10) | ACN:AmS 7:3 | 125mM (NH<sub>4</sub>)<sub>2</sub>SO<sub>4</sub>, 0.05% HFBA | 50 | 8 | 3000 Hz 15% | 3000 Hz 10% | × (+)<br>× (-)* | × (±) | × (+) | Pieles et. al.<sup>40</sup> |
| THAPnS (10) | ACN:H<sub>2</sub>O 7:3 | 0.05% HFBA | 50 | 8 | 3000 Hz 15% |  |  | × (+) |  | Pieles et. al.<sup>40</sup> |
| DAN70 (10) | ACN:H<sub>2</sub>O 7:3 |  | 30 | 8 | 3000 Hz 5% | 2000 Hz 1.5% | × (±) | × (±) | × (+) | Sun et. al.<sup>39</sup> |
| DAN90 (10) | ACN:H<sub>2</sub>O 9:1 |  | 30 | 8 | 3000 Hz 5% | 2000 Hz 1.5% |  | × (+) |  | Jackson et. al.<sup>24</sup> |
| DANtfa (10) | ACN:H<sub>2</sub>O 7:3 | 0.1% TFA | 30 | 8 | 3000 Hz 5% |  |  | × (+) |  | Sun et. al.<sup>39</sup> |
| DANhfba (10) | ACN:H<sub>2</sub>O 7:3 | 0.05% HFBA | 30 | 8 | 3000 Hz 5% |  |  | × (+) |  | Sun et. al.<sup>39</sup><br>Colsch et. al.<sup>23</sup> |
| DANw (10) | ACN:H<sub>2</sub>O 9:1 | AmAc wash | 30 | 8 | 3000 Hz 5% | 2000 Hz 1.5% | × (+) | × (±) |  | Sun et. al.<sup>39</sup><br>Angel et. al.<sup>43</sup> |
| * Hippocampus imaging with THAP/DHAP in negative ion mode performed at 20 μm spatial resolution | | | | | | | | | | |

### AP-MALDI-MSI

MALDI analysis on brain sections was performed using an AP-MALDI UHR ion source (Masstech Inc., USA), which has been described in detail elsewhere,^25-26^ coupled to an LTQ/Orbitrap Elite high-resolution mass spectrometer (Thermo-Fisher Scientific, USA) in positive and negative ion mode. For imaging, the AP-MALDI source was operated in “Constant Speed Raster” motion mode with a stage stepping size of 10 μm for hippocampus images and 40 μm for cerebellum and striatum (cholesterol) images. The laser spot size was < 10/40 μm (20 μm max.), settings are listed in Table 1. A laser focus of 8.38 μm was determined with scanning electron microscopy (SEM, Figure S 3) and, using the camera in the source and comparing line to line signal intensities, settings were adjusted for each measurement to ablate as much matrix as possible without oversampling. Spectrum acquisition: 800 ms maximum injection time; mass range: 500 – 2000 Da (250-1000 Da for cholesterol imaging); mass resolution: 120k at m/z 400. Tandem-MS was performed on 76 peaks: 1 Da isolation window, and collision-induced dissociation/ higher-energy collision dissociation (CID/HCD) was performed with collision energies of 20-55%, adjusted for each lipid species individually. Tandem-MS scans were summed up over 30-120 seconds. Details on matrix used, collision mode and energy can be found in the scan header of each analysis in the supplementary information 2. Data analysis and visualization was performed with Thermo Xcalibur 2.2 and Thermo ImageQuest (Thermo-Fisher Scientific, USA), METASPACE,^44^ MSiReader 1.2 (NC State University, USA),^45^ LipostarMSI (Molecular Horizons Srl, Italy) and OriginPro 2019b (OriginLab Corp., USA). All images are normalized to total ion count (TIC).

### ToF-SIMS-MSI

ToF-SIMS analysis was performed using an TOF.SIMS 5 (IONTOF GmbH, Germany) with a 25kV Bi_3_^+^ primary analysis beam. Dried brain sections with DAN70 matrix were analyzed in burst alignment, delayed extraction, positive ion mode with a total primary ion dose of 7 × 10^11^ ions/cm^2^, cycle time 105 us, random raster mode, 1 frame / patch, 1 shot / frame / pixel, 25 scans, mass range: 1 – 1000 Da, mass resolution: 5000 at m/z 300, image size: 256 × 256 μm and 512 × 512 pixels. Data analysis and visualization was performed using SurfaceLab 7 (IONTOF GmbH, Germany).

### SEM

SEM analysis was performed using a Quanta 200 Field Emission Gun Scanning Electron Microscope (FEG SEM, Philips-FEI, USA). DAN70 matrix analysis on brain was performed in a low-vacuum environment (60 Pa) with a “Large Field Detector” (LFD) for a topographical image. CHCA matrix analysis was performed with a Genesis XM 4i Energy Dispersive Spectrometer (from EDAX) system for elemental mapping and line-scans, with a back scattered, composition mode detector called BSED (for high vacuum), generating a greyscale, chemical composition SEM image, with “heavier” elements areas corresponding to brighter areas.

## Results

The aim of this study was to evaluate the performance of various matrices/sample preparation protocols for AP-MALDI in terms of lipid coverage, signal intensity and high-resolution imaging capability. For this purpose, several images were taken on sagittal mouse brain sections. Brain sections are often used in comparison studies, as they are rich in a great variety of lipids and the results can be compared to previously published material and entries in databases like e.g. METASPACE. Figure 1 shows an overview of the datasets included in this study (summarized in Table 1): Full brain sections (Fig. 1a, spatial resolution: 40 μm) were imaged to get a detailed overview of the lipid distributions in the brain and to determine the ideal location to perform tandem-MS on specific lipids. The location of 6 lipid species and representative spectra in positive and negative ion mode are shown in Figure 1b. Evidently, for species like SM(d16:1/24:1) is it important to know the location prior to tandem-MS analysis as it is only present in the ventricles (Fig 1b). The full brain datasets are available in METASPACE: AP_MALDI_Full_Brain. The ability to detect cholesterol was tested on a small area around the fiber tracts (Fig. 1c) with a resolution of 40 μm, in positive ion mode, and a mass range of 250-1000 Da. To test instrument and matrix performance for higher resolution imaging, the hippocampus with its intricate structures was imaged at 10 μm (Fig. 1d, Fig. S2) in positive and negative ion mode (DHAP and THAP at 20 μm). Hippocampus datasets in METASPACE: AP_MALDI_Hippocampus. To determine lipid coverage, the cerebellum was imaged in positive ion mode (Fig. 1e, Fig. S1) and negative ion mode (Fig 1f, Fig S1). DAN matrix performed well in positive and negative ion mode and attempts were made to improve its performance further, including tissue washing (DANw) and acidic additives (DANtfa, DANhfba). Initially HFBA was reported to stabilize DHAP in vacuum which would make it unnecessary in atmospheric conditions.^23^ Since HFBA is a strong acid, that similar to TFA is used as ion-pairing agent in liquid chromatography, we investigated its protonating and potentially ion enhancing effects.^46^ Cerebellum datasets in METASPACE : positive ion mode AP_MALDI_MATRIX_pos; negative ion mode AP_MALDI_MATRIX_neg. Brain tissue not imaged was used for on-tissue tandem-MS. For simplicity, phospholipid fragments resulting from a headgroup loss (e.g. PS -serine) are referred to as PA(x/x) and fatty acid fragments as FA(x/x). Images for all cerebellum and hippocampus analyses are displayed in the supplementary information 1 (Fig S1 and S2 respectively).

**Figure 1.**
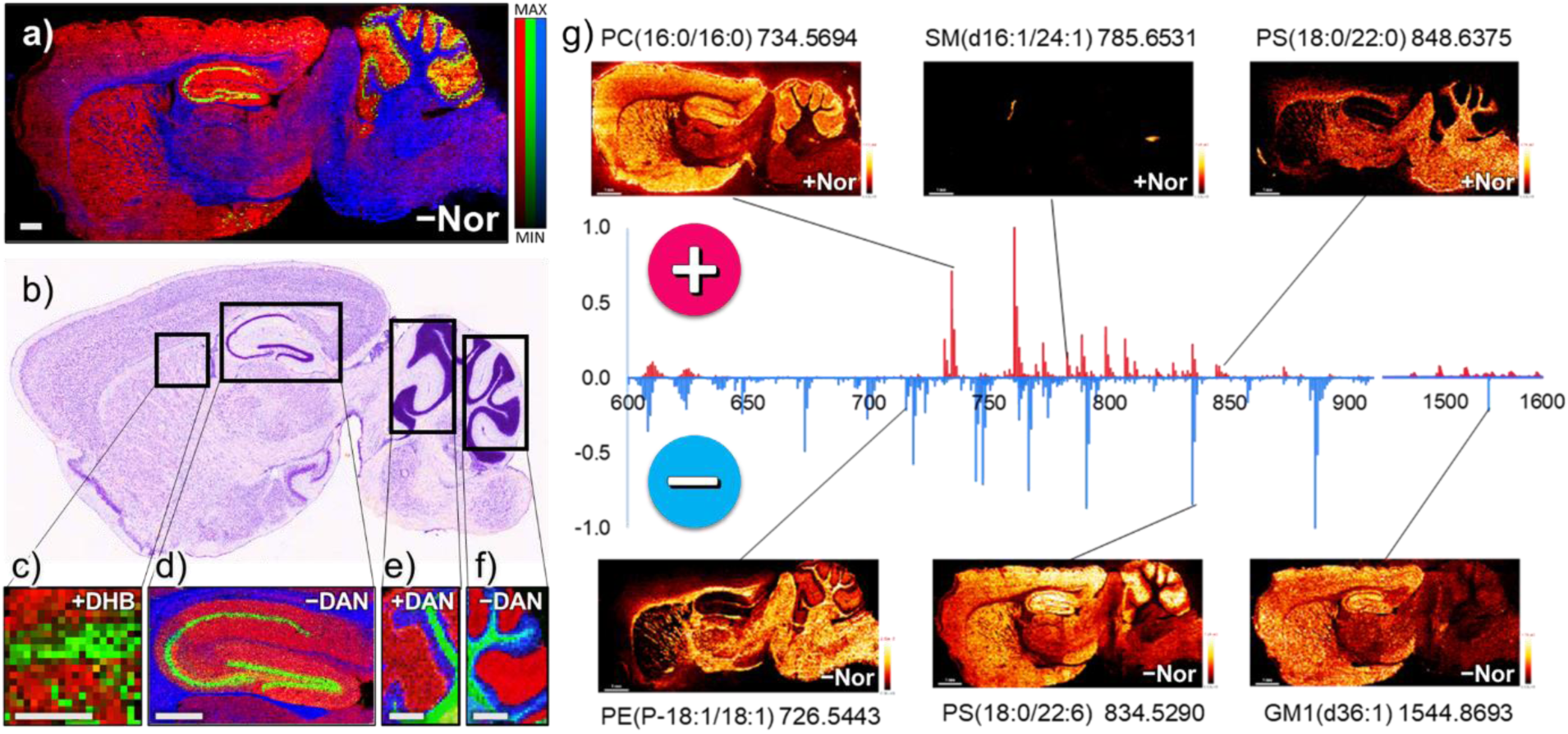
AP-MALDI imaging on sagittal brain section. A) Full brain AP-MALDI-image, 40 μm spatial resolution, (red: PS(40:6) green: PI(38:5), blue: C24:1-Sulf), b) H&E stained sagittal mouse brain section (Allen Developing Mouse Brain Atlas, dataset P56, sagittal), c-f) small area AP-MALDI-images of c) fiber tracts/striatum images for cholesterol analysis (40 μm, red: PC(32:0) green: Cholesterol), d) hippocampus imaged at 10 μm, (red: PS(40:6) green: PI(38:5), blue: C24:1-Sulf), e) cerebellum in positive ion mode (red: SM(d36:1), green: HexCer(d40:2), blue: PC(34:1)) and f) negative ion mode (40 μm, red: SM(d36:1), green: C24:1-Sulf, blue: PI(38:4)). Matrix and ion mode stated in each image, scalebar: 500 μm, g) 6 representative lipid images and spectra in positive and (mirroring) in negative ion mode.

### Instrumentation capabilities: Spatial resolution, mass resolution and tandem-MS

The MassTech AP-MALDI source in combination with an Orbitrap mass analyzer and an HTX TM sprayer for sample preparation, allows to attain MS-imaging with 10 μm spatial resolution while collecting high resolution mass spectra (up to 240k FWHM at m/z=400), and on-tissue tandem-MS spectra. Figure 2 shows the analysis of the hippocampus from a sagittal mouse brain section (also shown in Fig. 1d) and lipid species PS(18:1/18:0)-H at m/z 788.5447 and PC(16:1/22:6)-CH_3_ at m/z 788.5236. To distinguish those lipids, a resolving power of about 40k is necessary, which is well within the capabilities of the Orbitrap (Fig 2a). PC(16:1/22:6) is exclusively located in the hippocampus while PS(18:1/18:0) is mainly in the surrounding fiber tracts (Fig. 2b). Linescans across the hippocampus/fiber tract interface demonstrate that we can monitor chemical changes with a spatial resolution of 10 μm (Fig. 2c, position of the linescans is shown in 2b, indicated with white arrows).

**Figure 2.**
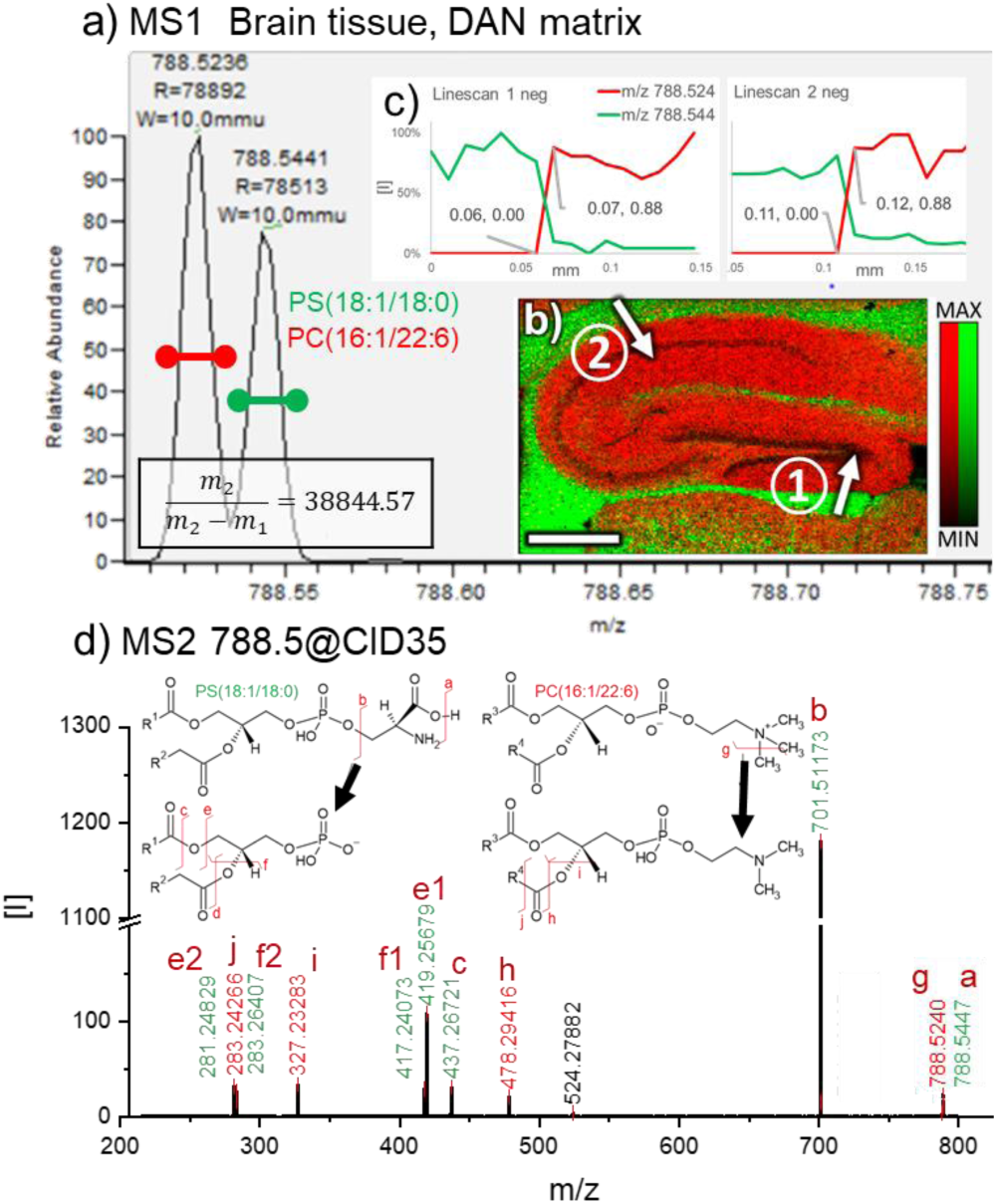
Capabilities of the AP-MALDI-Orbitrap system demonstrated on the hippocampus, a) MS1 scan on brain tissue (matrix: DAN70, negative ion mode, showing m/z 788.5447 and m/z 788.5236 b) Distribution of PS(18:1/18:0) (green) and PC(16:1/22:6)-CH_3_ (red) in a 10 μm AP-MALDI image, white arrows show the positions of linescans in c) scalebar: 500 μm.. c) 2 linescans, normalized to their individual maximum intensity (100%). d) On-tissue tandem-MS analysis of m/z 788.5 ± 0.5 Da, containing PS (green) and PC (red) fragments. Fragmentation mechanism shown for both lipids, referring to the letters above each mass peak.

SEM images show that DAN70 matrix applied with an HTX-TM sprayer produces ∼1 μm crystals and that the AP-MALDI laser can be focused below 10 μm (Fig. S3). At 10 μm spatial resolution sufficient signal is produced to assign 52 species with DAN70 in the hippocampus (72 with HCA, 134 with THAP) in positive ion mode, and 121 species in negative ion mode (METASPACE, database: LIPIDMAPS, FDR:20%). Significantly more species are assigned in the THAP dataset due to the slightly different analysis area which included the lateral ventricle with unique lipids. (The cerebellum datasets used to compare lipid coverage do not have this issue). In terms of spatial resolution, CHCA and DAN70 produce sharp images and perform better than THAP although all datasets were acquired with the same analysis settings (line spacing and scanning speed) and laser focus was < 10 μm. The reason could be that laser settings were adjusted for maximum signal and were higher for THAP than for other matrices. Less laser energy would have provided less signal but could have resulted in a sharper image.

Figure 2d shows a tandem-MS spectrum of m/z 788.5, and it demonstrates that AP-MALDI produces sufficient lipid signal on tissue to perform tandem-MS analysis. PS(18:1/18:0) and PC(16:1/22:6) are fairly well spatially separated in the hippocampus but can overlap in other brain areas and are therefore both present in the tandem-MS spectrum (Fig 1d). Due to the complexity of biological samples, it is to be expected that a tandem-MS spectrum will contain more than one lipid species, albeit with different intensities. Due to the high mass resolution and mass accuracy provided by the Orbitrap this is not an issue, as one still can identify both species and assign their fragments accordingly.

### Imaging single cells in tissues with AP-MALDI

Figure 3 shows that AP-MALDI imaging with DAN70 matrix has the potential of imaging individual neuron cells in tissues. PC(18:0/22:6) (Fig. 3a/c) shows a unique distribution, localized to small areas in the hippocampus. The associated microscopic image (Fig. 3b), taken before matrix application, shows small features corresponding to this distribution, that appear to be single cells. Their location, mainly in the *stratum oriens*, surrounding pyramidal cells (PC(18:0/22:6) blue, Fig. 3a), suggests that those are inhibitory neurons/basket cells. This is supported by the resemblance of the PC(18:0/22:6) hippocampus images (Fig 3a/c) to immunohistochemistry images of basket cells^47^ and the PC(18:0/22:6) distribution in the cerebellum, most intense in the region of the Purkinje cells (Fig 3d/e) that are surrounded by basket cells.^48^ This association will have to be confirmed in future experiments.

**Figure 3.**
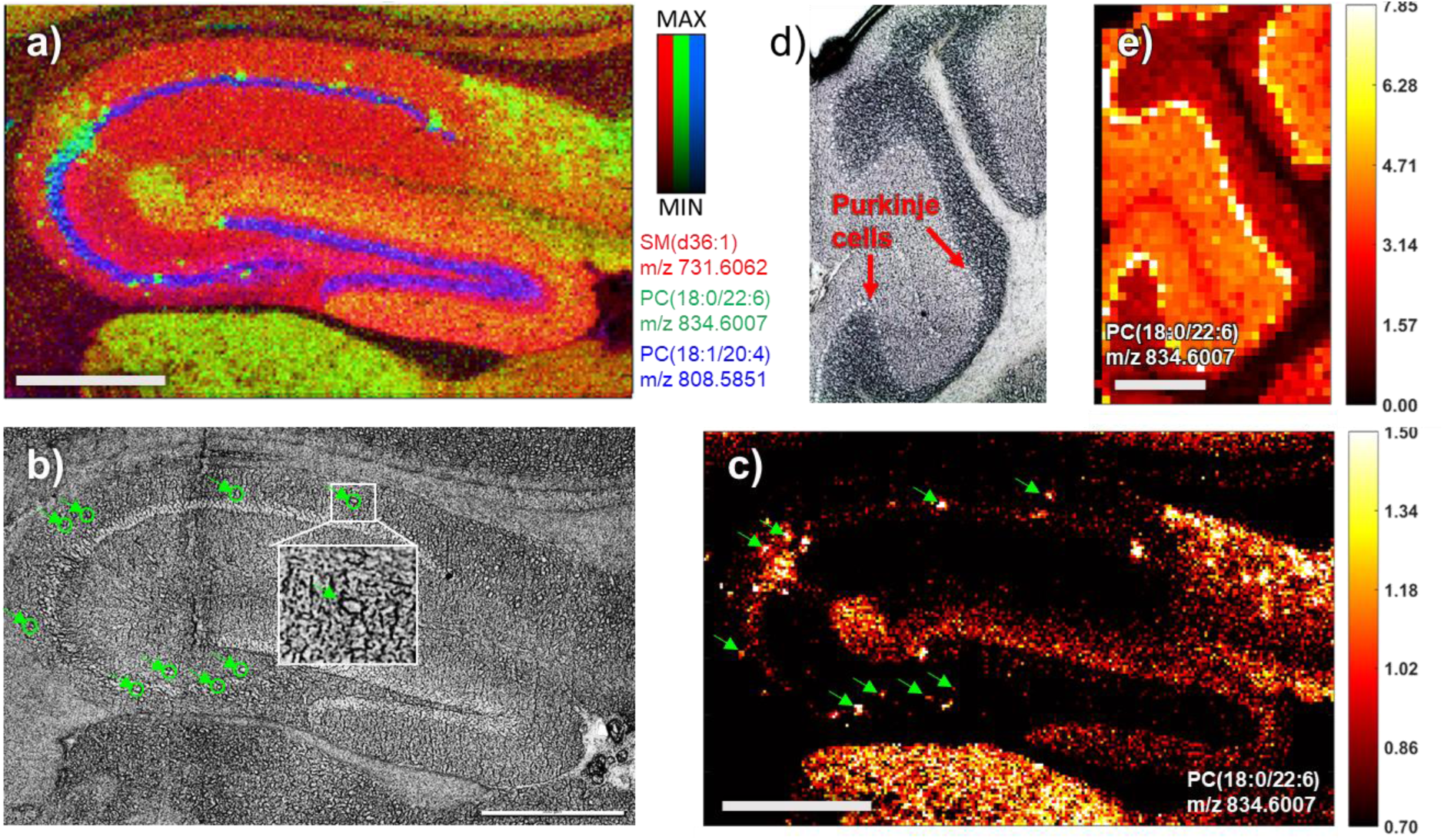
Single cell imaging in the hippocampus with AP-MALDI. A) 10 μm Hippocampus analysis (matrix: DAN70, positive ion mode), RGB overlay of: SM(d36:1) red, grey matter; PC(18:0/22:6) green, single cells; PC(18:1/20:4) blue, pyramidal layer. B) Microscope image of the hippocampus pre matrix application, features corresponding to the distribution of PC(18:0/22:6) highlighted in green. C) Single ion image of PC(18:0/22:6) with the same features highlighted as in B), color scale: hot. D) Microscope image of the cerebellum pre matrix application. E) Single ion image of PC(18:0/22:6) in the cerebellum, spatial resolution: 40 μm, color scale: hot. All scale-bars: 500 μm

### Lipid coverage with various matrices

Figure 4 shows the number of lipid species detected in the cerebellum (Fig. 1d/e) using all 11 sample preparation protocols for positive (Fig. 4a) and negative (Fig. 4b) ion mode, grouped into lipid classes (data was excluded if the most intense signals were below 1×10^3^ counts). Spectra for all included datasets in positive (Fig. S4) and negative ion mode (Fig S5) and signal to noise ratios for several signals (Fig. S6) can be found in the supplementary information 1. Using LipostarMSI and the LIPIDMAPS database, lipids were putatively assigned (Fig. 4a/e), with strict selection criteria (mass accuracy: 2 ppm; mass and isotopic pattern score: 80%+). The most species detected using one matrix were 155 in positive (THAP), 136 in negative ion mode (DAN70), and 224 for positive and negative ion mode combined (DAN70). Repeat measurements on different days for THAP (155 lipids 2020-12-01, 144 lipids 2020-10-19) and CHCA (136 lipids 2021-01-19, 124 lipids 2020-10-15) in positive ion mode, and DAN70 in positive (88 lipids 2020-10-07, 83 lipids 2021-01-25) and negative ion mode (136 lipids 2020-09-24, 120 lipids 2021-01-26) showed similar results (dataset included in the cerebellum METASPACE projects). Other parameters and/or databases (e.g. HMBD, SwissLipids) could lead to different results in terms of peak identity and the number of species, but this consistent approach was suitable to determine performance of each matrix/protocol.

**Figure 4.**
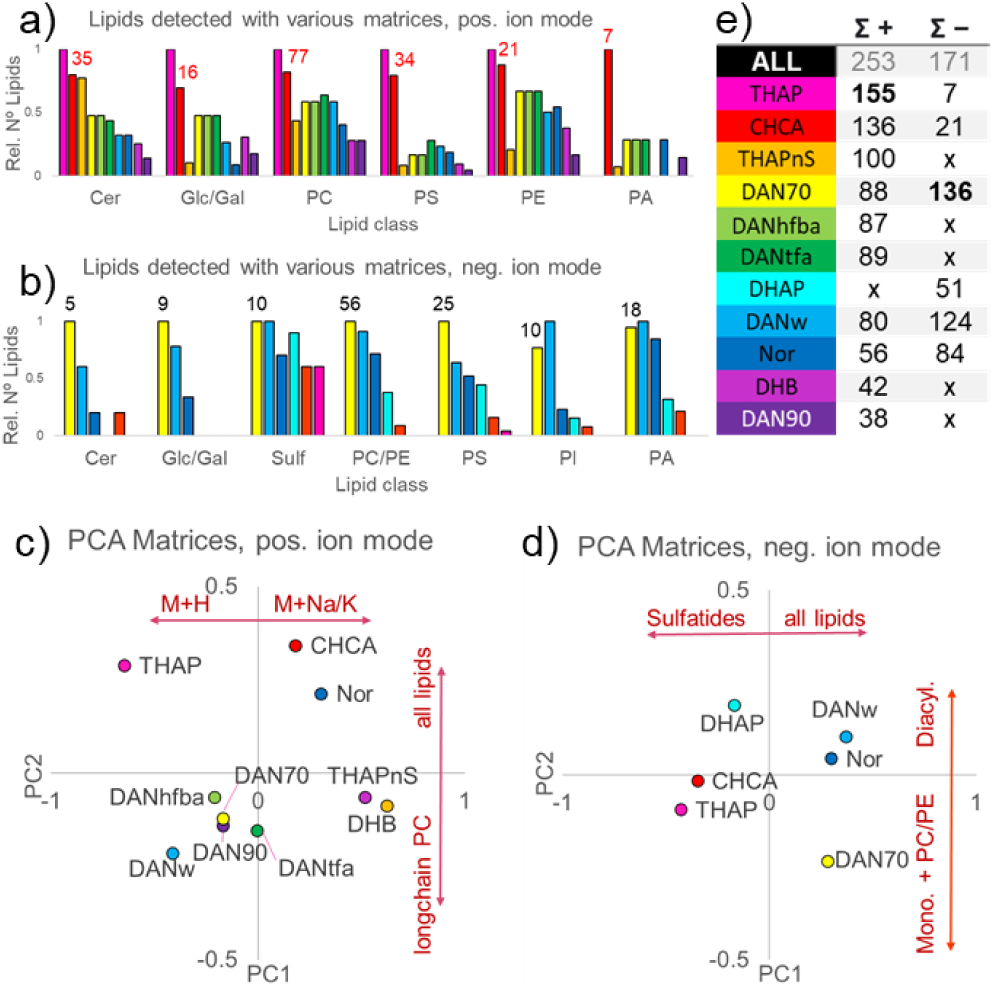
Lipid coverage for various matrices with AP-MALDI. A) Number of identified lipids by class, detected in the cerebellum using 5 matrices (10 sample preparation protocols) in positive ion mode and b) 5 matrices (6 sample preparation protocols) in negative ion mode listed in e). Displayed in the graph is the relative number of lipids normalized to the highest number of detected lipids for each class, the actual number is stated above CHCA (pos) and DAN70 (neg). C) Scores plots (PC1 vs PC2) of PCA analysis for cerebellum mass spectra in positive and d) negative ion mode (loadings plots: Fig. S7). E) Sum of assigned lipids for each matrix in positive and negative ion mode. Σ+ (x) DHAP was not analyzed in positive ion mode, Σ-(x) all matrices were analyzed in negative ion mode but excluded if insufficient signal was detected (below 1×10^3^ counts). Lipids were assigned using LipostarMSI.

In negative ion mode, DAN70 was the only matrix that enabled the detection of a broad range of lipid species while DHAP and THAP mainly produced sulfatides and ganglioside signals (Fig. 4b/d). Tissue washing (DANw) did increase signal intensities about 1.5-fold and double the S/N ratio for certain peaks (Fig. S6) compared to DAN70. Jackson *et al*. noted that DHAP is superior to DAN for detecting gangliosides, especially for intact GD1(d36:1) at m/z 1835.965. Similarly, in this study DHAP was the only matrix producing sufficient GD1 signal to perform on tissue tandem-MS (supplementary information 2).

PCA analysis of the spectra (average sum spectra for the whole cerebellum image, as shown in Fig. S1) shows that, in positive ion mode (Fig. 4c, loadings plots in Fig. S7), matrices CHCA, Nor and DHB produce higher intensity, sodiated/potassiated [M+Na/K]^+^ species while THAP and DAN (especially with AmAc wash, DANw) favor protonated [M+H]^+^ species. Also, DAN70 shows higher intensities for long chain fatty acid PC species. It has been reported that the addition of ammonium salts to a matrix, aids in suppressing sodium and potassium, especially for oligonucleotide analysis with THAP.^40, 49^ Similarly, we observed that THAPnS (no salt added) produces almost exclusively [M+Na/K]^+^ species. We did not observe a drastic signal increase after tissue washing (DANw), as was previously reported,^43, 50^ and no improvement in the number of detected species (Fig. 4e). The slight decrease in the number of detected species can be explained by the lack of salt adducts which often cause a single lipid species to be detected 3 times in positive ion mode [M+H/Na/K]^+^. The addition of acids, used in the CHCA and THAP protocols, to DAN (DANtfa, DANhfba) did not improve performance either.

The identity of 76 peaks (Table S1) was confirmed with on tissue tandem-MS (MS2 and MS3 for GD1(d36:1)) of which 60 were unique lipid species and 16 were repeat species detected as [M+H/Na/K]^+^, were detected in both ion modes (e.g. PE(18:0/22:6) as [M±H]^±^ at m/z 790.539 and 792.554), or peaks consisting of lipid dimers (e.g. m/z 1548.194 = PC(18:1/18:0)+PC(16:0/18:1), ≠ CL(78:1)). The mass accuracy (ppm) in full scan (MS1) and tandem-MS data are within ±1ppm (with lock mass) and ±3ppm (without lock mass). All tandem-MS spectra, with analysis conditions in the scan header plus: identified fragments, mass accuracy and molecular formulas of parent and fragment ions, are listed in the supplementary information 2. Lipid species and fragments were identified using a combination of the LipostarMSI lipid catalogue with rule based fragmentation entries, LIPIDMAPS and published literature.^51-59^

### Cholesterol imaging with AP-MALDI

Figure 5 shows images (Fig. 5a) and spectra (Fig. 5b) of the striatum/fiber tracts with cholesterol as [M-OH+H]^+^ species (m/z 369.3516). Cholesterol species [M − H]^+^ at m/z 385.3464, was detected at ∼10% of the intensity of m/z 369.3516. Cholesterol is not detectable with DAN70 and Nor, which produces low intensity signals in general. DHAP and THAP produced relatively low (below 1×10^3^ counts) and CHCA and DHB high cholesterol signals (1×10^4^ counts and above). While cholesterol detection was vastly different, other lipids were detected at similar levels in the range of 1-4×10^4^ (Fig. S8, lower for Nor). For DHB, cholesterol-related peaks were the most intense signals in the white matter.

**Figure 5.**
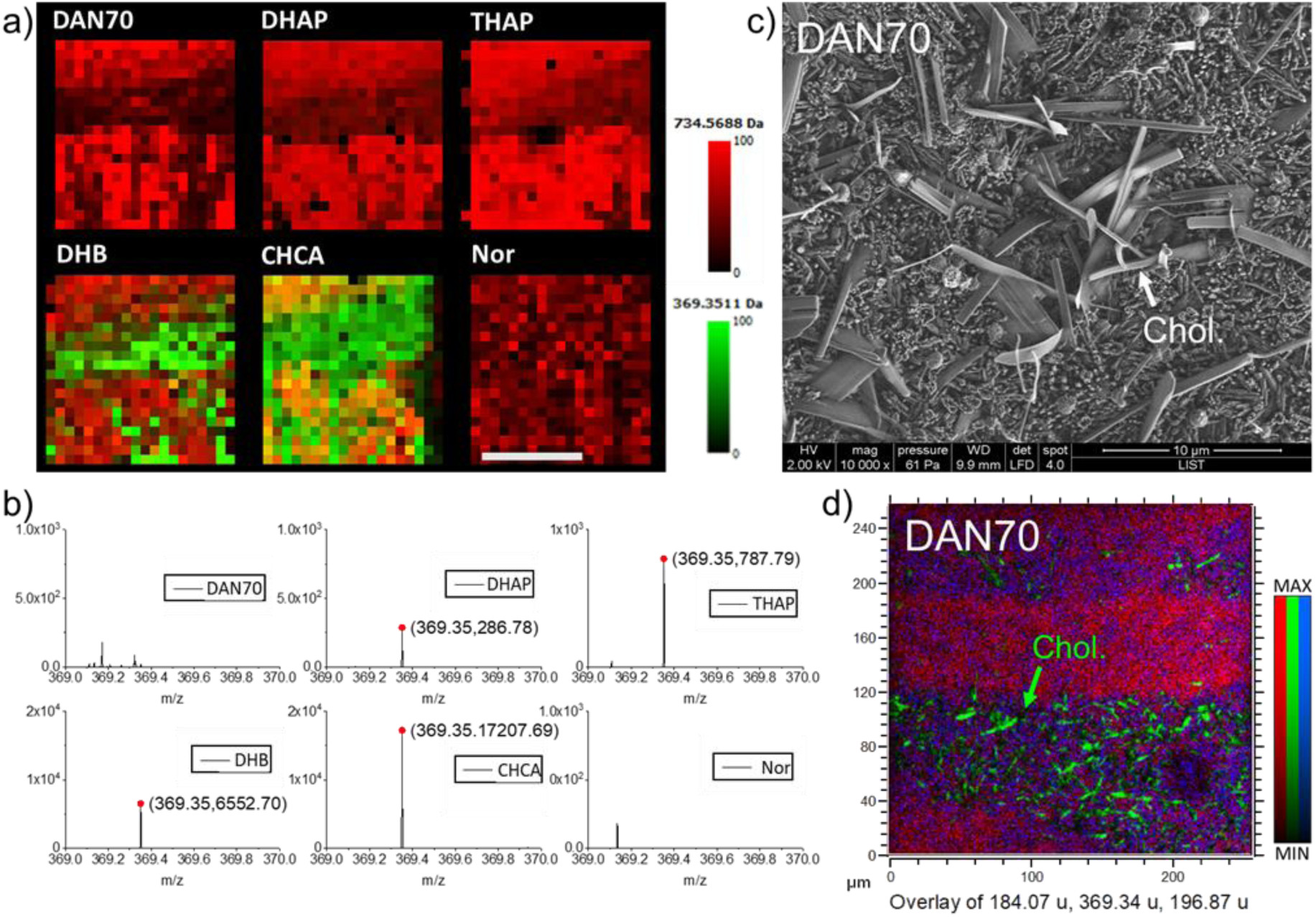
Cholesterol imaging with AP-MALDI, SEM and ToF-SIMS. A) AP-MALDI fiber tract images with various matrices showing cholesterol at m/z 369.3516 (green) and PC(16:0/16:0) at m/z 734.5688 (red). Scalebar: 500 μm b) Associated mass spectra showing the cholesterol peak and detection levels for each matrix. C) SEM and d) ToF-SIMS image of a brain section covered in DAN matrix, in the fiber tract region, cholesterol, m/z 369.35 (green); PC-headgroup, m/z 184.07 (red); m/z 196.87 (blue).

The tendency of cholesterol to migrate to the surface and crystalize during tissue drying has been reported previously.^60-61^ During matrix application, DAN70 does not seem to alter the cholesterol crystals. SEM images of DAN70 on brain sections show regular DAN crystals in the grey matter (Fig. S3). In the fiber tract region, additional larger spike-like crystals are visible (Fig. 5c). Those crystals resemble cholesterol crystals on dried tissue sections reported in the literature.^61^ Indeed, ToF-SIMS imaging on brain tissue covered with DAN70 matrix confirmed them to be cholesterol crystals (Fig. 5d). The addition of 0.1% TFA to DAN70 matrix did not alleviate the issue. No such crystals were found on the surfaces of tissue slices covered with CHCA or DHB, suggesting that these matrices can dissolve and incorporate cholesterol, therefore enabling its detection with MALDI.

## Discussion

We analyzed 11 sample preparation protocols with 6 different, vacuum stable and unstable matrices, to evaluate their performance for lipid imaging on brain tissue with AP-MALDI-Orbitrap-MSI and on-tissue tandem-MS.

Using strict selection criteria, we were able to assign 155/136 lipid species in AP-MALDI images in positive/negative ion mode. These results are comparable to vacuum MALDI-MSI approaches.^62^ AP-MALDI detected lipids with sufficient S/N to perform on-tissue tandem-MS, and with the added benefit of less pronounced tissue drying and cracking due to the harsh vacuum environment.^29, 32^ Of the studied matrices, DAN was the most versatile matrix (high spatial resolution, high signal intensity and lipids detected in positive+negative ion mode, similar results found for vacuum MALDI)^63^, but other matrices were superior for studying specific molecules (e.g. DHAP for gangliosides, DHB for cholesterol, more species detected in positive ion with THAP/CHCA). DAN protocol alterations such as tissue washing, and the addition of acids did increase performance but not drastically. Norharmane could be used in both ion modes as well but due to low signal intensity (even taking the low noise levels in the spectrum into account), many isotopical peaks fell below the signal-to-noise threshold and therefore fewer peaks were assigned with our selection criteria. Nevertheless, many lipids were present and the full brain analysis with Norharmane could be used to guide tandem-MS analysis.

The majority of MALDI-MSI publications state their imaging resolution/pixel size in terms of laser crater size or stage stepping size.^32^ This can be misleading, as it does not accurately reflect the spatial resolution for detecting chemical changes in the sample. In this study we demonstrated our spatial resolution of 10 μm with on-tissue linescans of lipid species, that combine laser focus, stage stepping size (or in our case, raster speed), and crystal size into one metric. A better laser focus is possible, but this would significantly reduce the signal. 10 μm corresponds to the size of one cell and was sufficient to correlate lipid species with single cells/neurons in the hippocampus (Fig. 3).

DHB used to be the gold-standard for MALDI analysis and it is still used in many matrix comparison studies.^38, 43^ Here, DHB was outperformed in all aspects by other matrices, other than for cholesterol imaging. Cholesterol is of interest to many scientists due to its abundance and its involvement in metabolism and disease.^64-65^ It can be easily detected with ToF-SIMS,^61^ but MALDI usually requires additional steps to enhance cholesterol ionization.^42, 66^ The main issue seems to be that cholesterol forms large, solid crystals on the sample surface. Even though they are destroyed by the laser, cholesterol is not ionized sufficiently without being integrated into (and co-crystallized with) the matrix. Therefore, only matrix protocols that can dissolve cholesterol crystals enable its detection with AP-MALDI-MSI. The DAN-matrix protocols used here left the cholesterol crystals intact, however, other lipids were still detected in the fiber tracts.

For tandem-MS analysis, THAP and DAN70 worked comparatively well due to their high signal intensities and lipid coverage. For most lipids assigned with LipostarMSI and METASPACE, tandem-MS confirmed their identity in accordance with their possible assignments. Only the assigned cardiolipins detected in positive ion mode were discovered to be lipid dimers instead. This highlights the importance of tandem-MS analysis, not only for the structural elucidation of the detected species but also for confident assignments.

All datasets included in this study contained hundreds of assigned species with different distributions. This data would have been too vast to include in this manuscript. Therefore, for transparency all datasets were uploaded to METASPACE, where their lipid distributions can be viewed. Additionally, the METASPCE projects, created for this publication, can be expanded upon, as further analyses with novel matrix compounds are performed. Links to all dataset: full brain positive/negative: AP_MALDI_Full_Brain; hippocampus positive/negative: AP_MALDI_Hippocampus; cerebellum positive: AP_MALDI_MATRIX_pos; cerebellum negative: AP_MALDI_MATRIX_neg.

## Conclusion

For a long time, lipids have taken a backseat to proteins concerning their importance in disease mechanisms. This was partially due to the inability to track the changes in lipid distributions in heterogeneous samples, changes which can be very subtle in homogenized sample extracts. MSI techniques like AP-MALDI-imaging can capture those changes, but data quality can depend strongly on sample preparation. We tested 11 sample preparation protocols for 6 matrices and the instrument capabilities of the AP-MALDI-Orbitrap system for lipidomics studies. We defined their characteristics in terms of lipid coverage, signal intensities, high spatial resolution imaging capability and usability for positive and negative ion mode. Every matrix had its advantages and disadvantages and knowing their characteristics is crucial for deciding which one is best suited for the scientific needs of a study. For now, we recommend THAP for on tissue tandem-MS, CHCA or DAN70 for high spatial resolution imaging in positive ion mode, and DAN70 for high spatial resolution imaging and tandem-MS in negative ion mode. The matrix recipes were adapted from previous MALDI/AP-MALDI publications and had all been optimized by their respective users, but they ultimately represented only a small number of options. Therefore, we intend to keep testing matrices and expanding the publicly available datasets in METASPACE. We will publish an update, should a novel matrix prove to be superior. In conclusion, AP-MALDI has shown to be comparable to vacuum MALDI for lipid detection, with the added benefit of less pronounced tissue drying and no requirement for vacuum stable matrices. In addition, the AP-MALDI source in combination with Orbitrap-MS allows for a direct transposition of conventional LC-MS fragmentation parameters for lipids. Therefore, AP-MALDI can be considered cost-effective addition to widely available LC/HRMS instruments and a valuable asset in applied, biomedical research.

## Supporting Information

The supplementary information 1 includes additional AP-MALDI images for all datasets included in this study, SEM images of DAN70 and CHCA matrices, a table containing all identified lipids, mass spectra for cerebellum analysis with all matrices in pos/neg ion mode and signal to noise levels for selected species, PCA loadings plots, and images and spectra of the small area - striatum analyses. The supplementary information 2 includes tandem-MS data for 76 peaks with assigned fragments.

## Author Contributions

TBA and GF designed the experiments. JB performed the ToF-SIMS analysis. JLB performed the SEM analysis. TBA performed the AP-MALDI analyses and wrote the manuscript. All authors commented on the manuscript and discussed the results.

## Acknowledgements

The Authors want to thank Roberto Angelini, Eylan Yutuc and William J. Griffiths from the Medical School, Swansea University, UK for providing the brain tissue sections, MassTech Inc. for their financial and material support of the AP-MALDI demo lab at LIST, and Venkat Panchagnula for proofreading the manuscript. The project was supported by the Luxembourg National Research Fund (FNR) (SKIMAS, Project number: 14292830)

