## supplementary information 1 for "Evaluation of 6 MALDI-Matrices for 10 µm lipid imaging and on-tissue MSn with AP-MALDI-Orbitrap"

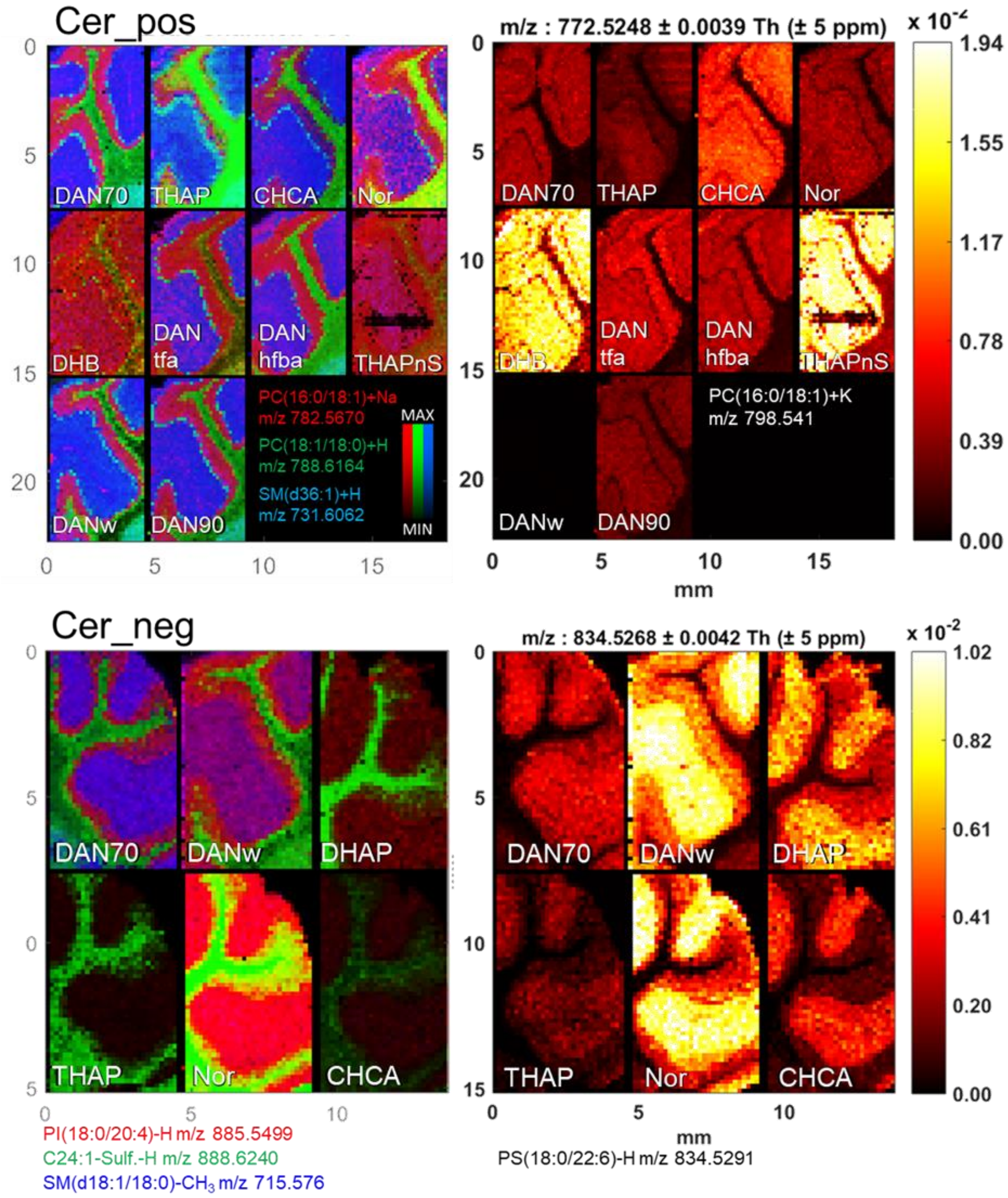

*Figure S 1 All AP-MALDI cerebellum datasets in positive (top) and negative (bottom) ion mode. Matrix stated in left, bottom corner of each image. Displayed lipid species stated underneath each image group for RGB and single ion images.*

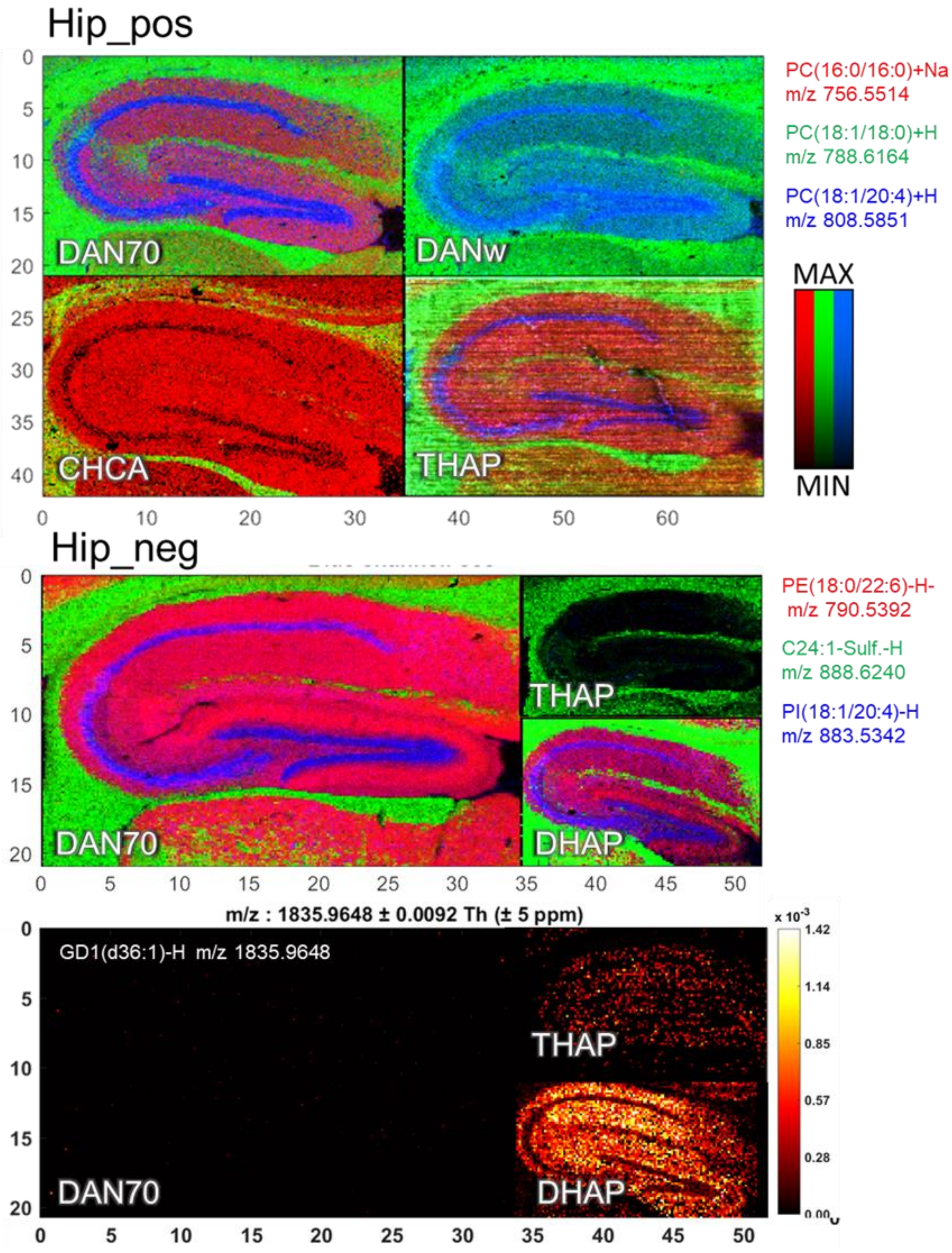

*Figure S 2 All AP-MALDI hippocampus datasets in positive (top) and negative (bottom) ion mode. Matrix stated in left, bottom corner of each image. Displayed lipid species stated next to each image group for RGB and single ion images.*

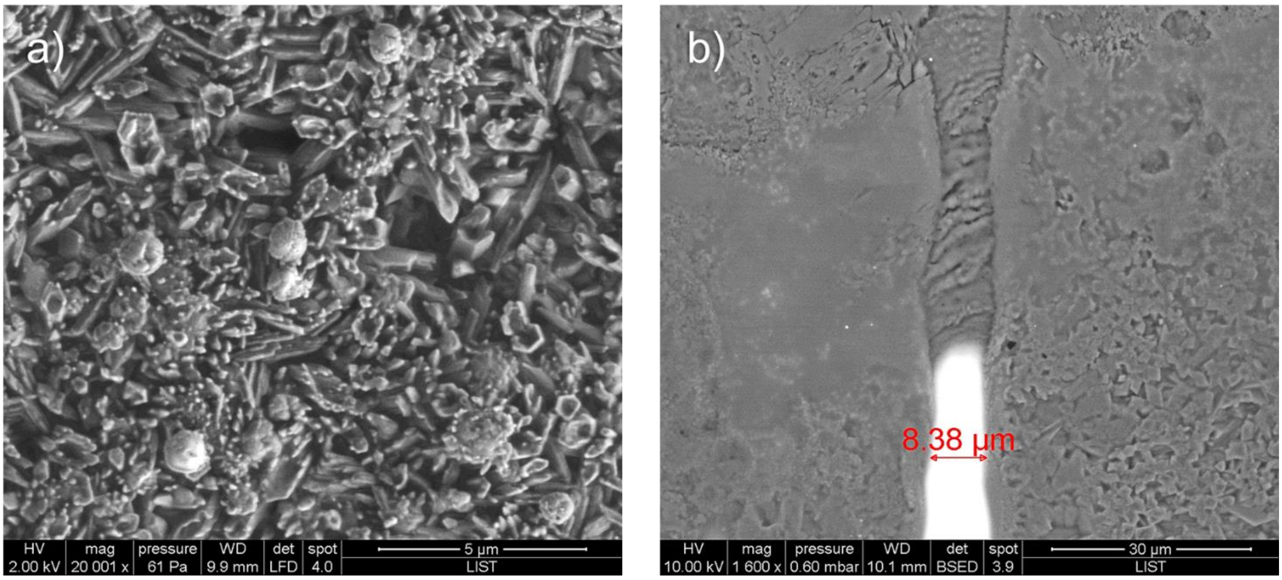

*Figure S 3 SEM images of a) DAN matrix crystals on tissue and b) a laser track through CHCA matrix on ITO glass.*

*Table S 1. Lipid species identified with tandem MS.*

**Positive ion mode**

| **Assignment** | Species | m/z (measured) | Formula | m/z (exact) | Δppm |
|---|---|---|---|---|---|
| **PC(16:0/9:0(OH))** | [M+H]$^+$ | 650.4372 | $C_{33}H_{65}O_9NP$ | 650.439 | 3.00 |
| **PC(16:0/9:0(COOH))** | [M+H]$^+$ | 666.4321 | $C_{33}H_{65}O_{10}NP$ | 666.434 | 2.94 |
| **PC(18:0/9:0(OH))** | [M+H]$^+$ | 678.4698 | $C_{35}H_{69}O_9NP$ | 678.471 | 0.96 |
| **PC(16:0/16:0)-TMA** | [M+K]$^+$ | 713.4497 | $C_{37}H_{71}O_8PK$ | 713.452 | 2.95 |
| **PDME(16:0/16:0)** | [M+H]$^+$ | 720.5543 | $C_{39}H_{79}O_8NP$ | 720.554 | -0.72 |
| **SM(d36:1)** | [M+H]$^+$ | 731.606 | $C_{41}H_{84}O_6N_2P$ | 731.606 | 0.21 |
| **PC(16:0/16:0)** | [M+H]$^+$ | 734.5689 | $C_{40}H_{81}O_8NP$ | 734.569 | 0.72 |
| **PC(16:0/16:0)** | [M+Na]$^+$ | 756.553 | $C_{40}H_{80}O_8NPNa$ | 756.551 | -2.14 |
| **PC(16:0/18:1)** | [M+H]$^+$ | 760.5826 | $C_{42}H_{83}O_8NP$ | 760.585 | 3.26 |
| **PC(16:0/16:0)** | [M+K]$^+$ | 772.5254 | $C_{40}H_{80}O_8NPK$ | 772.525 | -0.12 |
| **PC(16:0/18:1)** | [M+Na]$^+$ | 782.5685 | $C_{42}H_{82}O_8NPNa$ | 782.567 | -1.88 |
| **HexCer(d18:1/22:1)** | [M+H]$^+$ | 782.6489 | $C_{46}H_{88}O_8N$ | 782.651 | 1.98 |
| **SM(d40:2)** | [M+H]$^+$ | 785.6534 | $C_{45}H_{90}O_6N_2P$ | 785.653 | -0.38 |
| **PC(18:1/18:1)** | [M+H]$^+$ | 786.6004 | $C_{44}H_{85}O_8NP$ | 786.601 | 0.42 |
| **PC(18:1/18:0)** | [M+H]$^+$ | 788.6161 | $C_{44}H_{87}O_8NP$ | 788.616 | 0.36 |
| **PE(40:7)*** | [M+H]$^+$ | 790.5357 | $C_{45}H_{77}O_8NP$ | 790.538 | 3.07 |
| **PE(18:0/22:6)** | [M+H]$^+$ | 792.5555 | $C_{45}H_{79}O_8NP$ | 792.554 | -2.17 |
| **PC(16:0/18:1)** | [M+K]$^+$ | 798.5403 | $C_{42}H_{82}O_8NPK$ | 798.541 | 0.83 |
| **PC(16:0/22:6)** | [M+H]$^+$ | 806.5691 | $C_{46}H_{81}O_8NP$ | 806.569 | 0.41 |
| **PE(38:4)** | [M+K]$^+$ | 806.5067 | $C_{43}H_{78}NO_8PK$ | 806.510 | 3.67 |
| **PC(18:1/20:4)** | [M+H]$^+$ | 808.5839 | $C_{46}H_{83}O_8NP$ | 808.585 | 1.45 |
| **HexCer(d18:1/24:2)** | [M+H]$^+$ | 808.6642 | $C_{48}H_{90}O_8N$ | 808.666 | 2.34 |
| **SM(d42:2)** | [M+H]$^+$ | 813.6846 | $C_{47}H_{94}O_6N_2P$ | 813.684 | -0.25 |
| **PC(18:1/18:0)*** | [M+K]$^+$ | 826.5739 | $C_{44}H_{86}O_8NPK$ | 826.572 | -1.98 |
| **PC(18:0/22:6)** | [M+H]$^+$ | 834.6012 | $C_{48}H_{85}O_8NP$ | 834.601 | -0.56 |
| **PC(38:6)** | [M+K]$^+$ | 844.5228 | $C_{46}H_{80}NO_8PK$ | 844.525 | 2.97 |
| **PC(38:4)** | [M+K]$^+$ | 848.5556 | $C_{46}H_{84}O_8NPK$ | 848.557 | 1.19 |
| **HexCer(d18:1/24:1(2OH))** | [M+Na]$^+$ | 848.6564 | $C_{48}H_{91}O_9NNa$ | 848.659 | 2.59 |
| **PC(40:6)** | [M+K]$^+$ | 872.5565 | $C_{48}H_{85}O_8NPK$ | 872.557 | 0.13 |
| **PC(16:0/18:1)+PC(16:0/16:0)** | [M+M+H]$^+$ | 1494.1488 | $C_{82}H_{163}O_{16}N_2P_2$ | 1494.147 | -1.04 |
| **PC(16:0/18:1)** | [2M+H]$^+$ | 1520.1604 | $C_{84}H_{165}O_{16}N_2P_2$ | 1520.163 | 1.64 |
| **PC(18:1/18:0)+PC(16:0/18:1)** | [M+M+H]$^+$ | 1548.1868 | $C_{86}H_{169}O_{16}N_2P_2$ | 1548.194 | 4.64 |

### Negative ion mode

| Assignment | Species | m/z (measured) | Formula | Da | ppm |
|---|---|---|---|---|---|
| **Cer(d18:0/18:1)** | $[M-H]^-$ | 564.535 | $C_{36}H_{70}O_3N$ | 564.535 | 0.04 |
| **CerP(d18:1/18:0)** | $[M-H]^-$ | 644.5019 | $C_{36}H_{71}NO_6P$ | 644.503 | 0.85 |
| **PA(16:0/16:1)** | $[M-H]^-$ | 645.4503 | $C_{35}H_{66}O_8P$ | 645.450 | -0.34 |
| **PA(16:0/16:0)** | $[M-H]^-$ | 647.4645 | $C_{35}H_{68}O_8P$ | 647.466 | 1.9 |
| **CerP(d38:2)** | $[M-H]^-$ | 670.5173 | $C_{38}H_{73}O_6NP$ | 670.518 | 1.19 |
| **PA(16:0/18:1)** | $[M-H]^-$ | 673.4795 | $C_{37}H_{70}O_8P$ | 673.481 | 2.79 |
| **SM(d34:1)** | $[M-CH_3]^-$ | 687.5426 | $C_{38}H_{76}O_6N_2P$ | 687.545 | 2.98 |
| **SM(d18:1/18:0)** | $[M-CH_3]^-$ | 715.575 | $C_{40}H_{80}O_6N_2P$ | 715.576 | 1.33 |
| **PE(16:0/18:0)** | $[M-H]^-$ | 718.5377 | $C_{39}H_{77}O_8NP$ | 718.539 | 2.13 |
| **PE(P-18:1/18:1)** | $[M-H]^-$ | 726.5427 | $C_{41}H_{77}O_7NP$ | 726.544 | 2.22 |
| **PE(P-18:0/18:1)** | $[M-H]^-$ | 728.5584 | $C_{41}H_{79}O_7NP$ | 728.560 | 2.14 |
| **PE(18:1/18:1)** | $[M-H]^-$ | 742.5391 | $C_{41}H_{77}O_8NP$ | 742.539 | 0.17 |
| **PC(16:0/18:1)** | $[M-CH_3]^-$ | 744.5535 | $C_{41}H_{79}O_8NP$ | 744.555 | 1.85 |
| **PA(18:0/22:6)** | $[M-H]^-$ | 747.4959 | $C_{43}H_{72}O_8P$ | 747.497 | 1.51 |
| **PE(P-18:1/20:1)** | $[M-H]^-$ | 754.5749 | $C_{43}H_{81}O_7NP$ | 754.576 | 0.94 |
| **PE(18:0/20:4)** | $[M-H]^-$ | 766.5373 | $C_{43}H_{77}O_8NP$ | 766.539 | 2.52 |
| **SM(d18:1/22:0)** | $[M-CH_3]^-$ | 771.637 | $C_{44}H_{88}O_6N_2P$ | 771.639 | 2.01 |
| **PE(P-18:0/22:6)** | $[M-H]^-$ | 774.5427 | $C_{45}H_{77}O_7NP$ | 774.544 | 2.08 |
| **PE(18:0/20:4(OH)** | $[M-H]^-$ | 782.5347 | $C_{43}H_{77}O_9NP$ | 782.534 | -0.72 |
| **PC(16:1/22:6)** | $[M-CH_3]^-$ | 788.5244 | C45H75O8NP | 788.524 | -1.04 |
| **PS(18:1/18:0)** | $[M-H]^-$ | 788.5447 | $C_{42}H_{79}O_{10}NP$ | 788.545 | 0.01 |
| **PE(18:0/22:6)** | $[M-H]^-$ | 790.5374 | $C_{45}H_{77}O_8NP$ | 790.539 | 2.31 |
| **HexCer(d18:1/22:0(2OH))** | $[M-H]^-$ | 798.6443 | $C_{46}H_{88}O_9N$- | 798.647 | 2.70 |
| **C18:1-Sulf** | $[M-H]^-$ | 806.5436 | $C_{42}H_{80}O_{11}NS$ | 806.546 | 2.68 |
| **C18(OH)-Sulf** | $[M-H]^-$ | 822.5411 | $C_{42}H_{80}O_{12}NS$ | 822.541 | -0.52 |
| **HexCer(d18:1/24:0(2OH))** | $[M-H]^-$ | 826.6757 | $C_{48}H_{92}O_9N$ | 826.678 | 2.49 |
| **PS(18:0/22:6)** | $[M-H]^-$ | 834.5274 | $C_{46}H_{77}O_{10}NP$ | 834.529 | 1.99 |
| **PS(18:1/22:0)** | $[M-H]^-$ | 844.6081 | $C_{46}H_{87}O_{10}NP$ | 844.607 | -0.93 |
| **PI(16:0/20:4)** | $[M-H]^-$ | 857.518 | $C_{45}H_{78}O_{13}P$ | 857.519 | 0.64 |
| **C22-Sulf.** | $[M-H]^-$ | 862.6062 | $C_{46}H_{88}O_{11}NS$ | 862.608 | 2.50 |
| **PI(18:0/18:1)** | $[M-H]^-$ | 863.5636 | $C_{45}H_{84}O_{13}P$ | 863.566 | 2.20 |
| **PS(18:1/24:0)** | $[M-H]^-$ | 872.6341 | $C_{48}H_{91}O_{10}NP$ | 872.639 | 5.12 |
| **C22(OH)-Sulf** | $[M-H]^-$ | 878.6008 | $C_{46}H_{88}O_{12}NS$ | 878.603 | 2.81 |
| **PI(18:1/20:4)** | $[M-H]^-$ | 883.5321 | $C_{47}H_{80}O_{13}P$ | 883.534 | 2.38 |
| **PI(18:0/20:4)** | $[M-H]^-$ | 885.5493 | $C_{47}H_{82}O_{13}P$ | 885.550 | 0.62 |
| **C24:1-Sulf.** | $[M-H]^-$ | 888.6228 | $C_{48}H_{90}O_{11}NS$ | 888.624 | 1.36 |
| **C24-Sulf.** | $[M-H]^-$ | 890.6376 | $C_{48}H_{92}O_{11}NS$ | 890.640 | 2.31 |
| **PI(18:0/20:4(OH))** | $[M-H]^-$ | 901.5425 | $C_{47}H_{82}O_{14}P$ | 901.545 | 2.52 |
| **C24(OH)-Sulf.** | $[M-H]^-$ | 906.6333 | $C_{48}H_{92}O_{12}NS$ | 906.635 | 1.40 |
| **Gal-GalNAc-Gal-Glc-(d36:1)** | $[M-H]^-$ | 1254.777 | $C_{61}{}^{13}CH_{113}O_{23}N_2$ | 1254.777 | 0.26 |
| **GM1(d36:1)** | $[M-H]^-$ | 1544.8632 | $C_{73}H_{130}O_{31}N_3$ | 1544.869 | 4.00 |
| **GM1(d38:1)** | $[M-H]^-$ | 1573.899 | $C_{74}{}^{13}C\ H_{134}O_{31}N_3$ | 1573.904 | 3.19 |
| **GD1(d36:1)** | $[M-H]^-$ | 1835.957 | $C_{84}H_{147}O_{39}N_4$ | 1835.965 | 4.24 |
| **GD1(d36:1)** | $[M-2H+K]^-$ | 1873.916 | $C_{84}H_{146}O_{39}N_4K$ | 1873.921 | 2.71 |

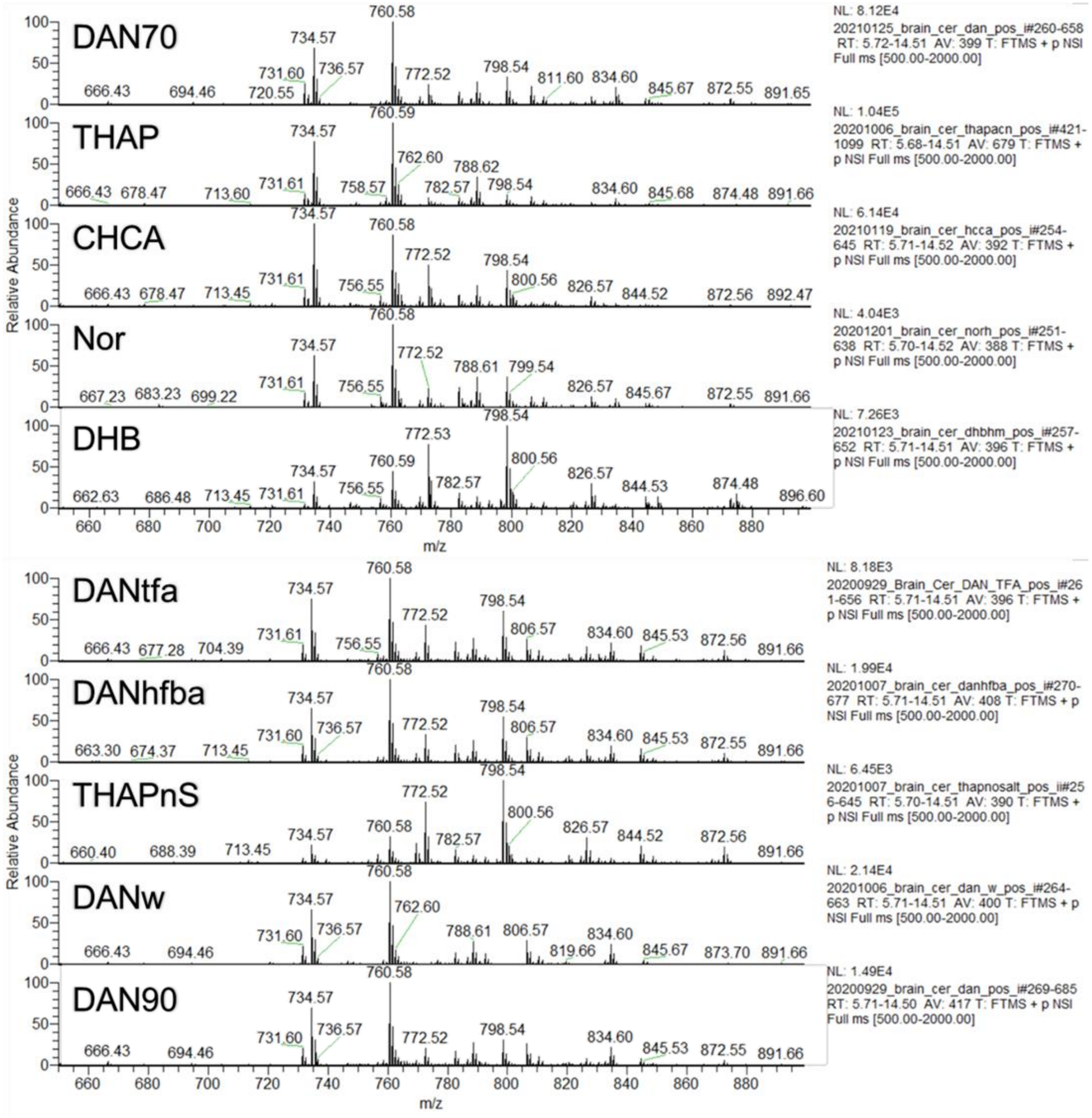

*Figure S 4 Cerebellum spectra of the lipid region for all matrices studied in pos. ion mode.*

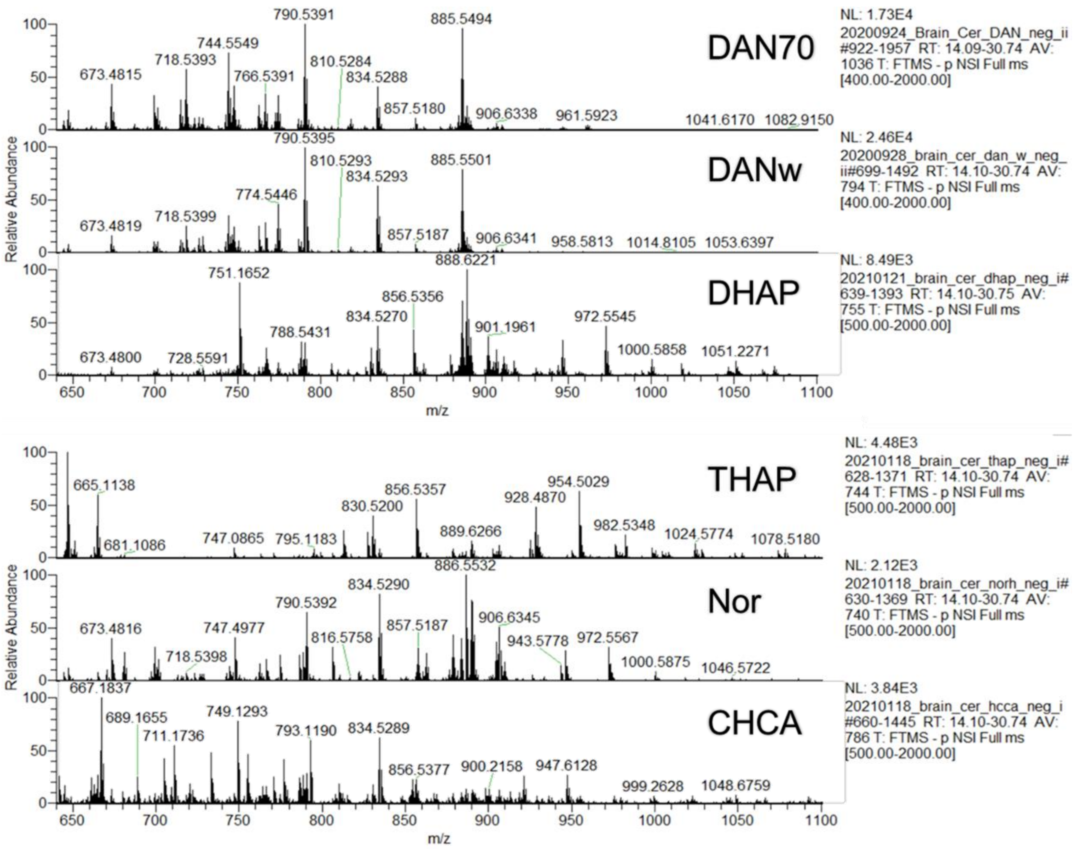

*Figure S 5 Cerebellum spectra of the lipid region for all matrices studied in neg. ion mode.*

| POS | SM(d36:1)+H m/z 731.60 | | |
|---|---|---|---|
| Matrix | S | N | S/N |
| DAN70 | 2.05E+04 | 11.33 | 1809 |
| THAP | 1.43E+04 | 12.56 | 1139 |
| CHCA | 1.32E+04 | 14.89 | 887 |
| Nor | 7.20E+02 | 2.05 | 351 |
| DHB | 3.91E+02 | 8.39 | 47 |
| DANtfa | 1.61E+03 | 2.02 | 797 |
| DANhfba | 3.91E+03 | 2.43 | 1609 |
| THAPnS | 4.32E+02 | 1.56 | 277 |
| DANw | 4.67E+03 | 2.21 | 2113 |
| DAN90 | 3.18E+03 | 2.46 | 1293 |

| POS | PC(16:0/18:1)+H m/z 760.58 | | |
|---|---|---|---|
| Matrix | S | N | S/N |
| DAN70 | 8.12E+04 | 11.75 | 6911 |
| THAP | 1.04E+05 | 13.01 | 7994 |
| CHCA | 5.35E+04 | 16.34 | 3274 |
| Nor | 4.04E+03 | 2.1 | 1924 |
| DHB | 3.28E+03 | 8.98 | 365 |
| DANtfa | 8.18E+03 | 2.1 | 3895 |
| DANhfba | 1.99E+04 | 2.53 | 7866 |
| THAPnS | 2.06E+03 | 1.6 | 1288 |
| DANw | 2.14E+04 | 2.27 | 9427 |
| DAN90 | 1.49E+04 | 2.56 | 5820 |

| POS | PC(16:0/18:1)+K m/z 798.54 | | |
|---|---|---|---|
| Matrix | S | N | S/N |
| DAN70 | 2.75E+04 | 12.29 | 2238 |
| THAP | 1.31E+04 | 13.61 | 963 |
| CHCA | 2.66E+04 | 17.77 | 1497 |
| Nor | 1.49E+03 | 2.16 | 690 |
| DHB | 7.28E+03 | 9.66 | 754 |
| DANtfa | 4.99E+03 | 2.19 | 2279 |
| DANhfba | 1.10E+04 | 2.66 | 4135 |
| THAPnS | 6.45E+03 | 1.65 | 3909 |
| DANw | 8.17E+01 | 2.34 | 35 |
| DAN90 | 4.67E+02 | 2.68 | 174 |

| NEG | PE(18:0/22:6)-H m/z 790.54 | | |
|---|---|---|---|
| Matrix | S | N | S/N |
| DAN70 | 1.81E+04 | 8.96 | 2020 |
| DANw | 1.33E+04 | 2.83 | 4700 |
| DHAP | 2.49E+03 | 7.81 | 319 |
| THAP | 7.45E+01 | 5.97 | 12 |
| Nor | 1.33E+03 | 5.76 | 231 |
| CHCA | 1.01E+03 | 6.94 | 146 |

| NEG | PI(18:0/20:4)-H m/z 885.55 | | |
|---|---|---|---|
| Matrix | S | N | S/N |
| DAN70 | 2.21E+04 | 14.18 | 1559 |
| DANw | 1.25E+04 | 4.39 | 2847 |
| DHAP | 5.94E+03 | 11.41 | 521 |
| THAP | n.d. | n.d. | |
| Nor | 2.28E+01 | 9.19 | 2 |
| CHCA | 6.26E+01 | 11.34 | 6 |

| NEG | C24:1-Sulf-H m/z 888.62 | | |
|---|---|---|---|
| Matrix | S | N | S/N |
| DAN70 | 4.26E+04 | 9.95 | 4281 |
| DANw | 3.21E+03 | 3.47 | 925 |
| DHAP | 4.94E+03 | 10.72 | 461 |
| THAP | 1.92E+01 | 8.44 | 2 |
| Nor | 1.07E+01 | 8.22 | 1 |
| CHCA | 1.03E+01 | 10.59 | 1 |

*Figure S 6 Signal to noise ratios (S/N) for various signals in positive and negative ion mode.*

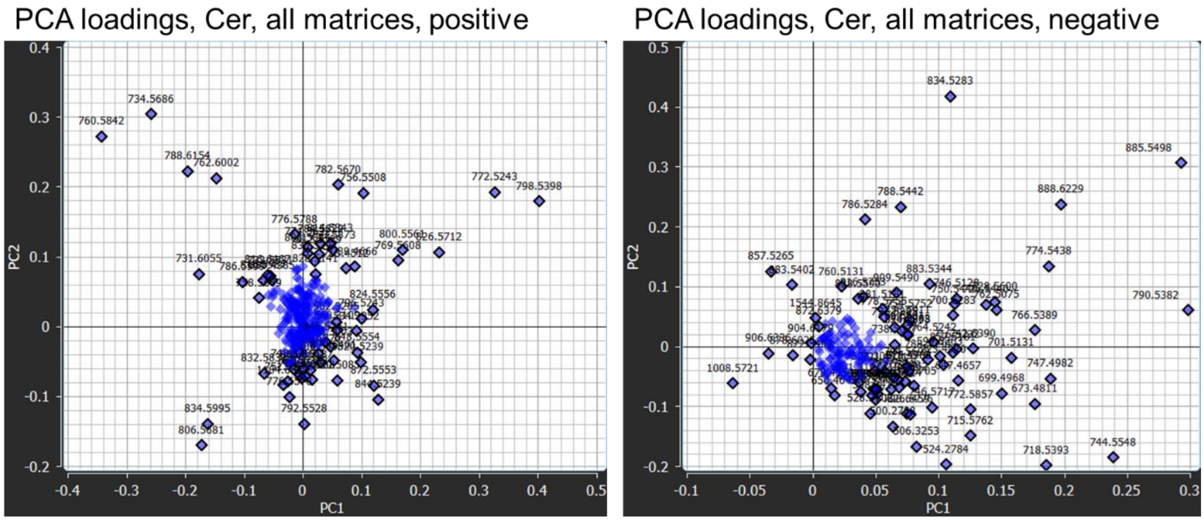

*Figure S 7 PCA loadings plots for cerebellum spectra analysis in positive and negative ion mode.*

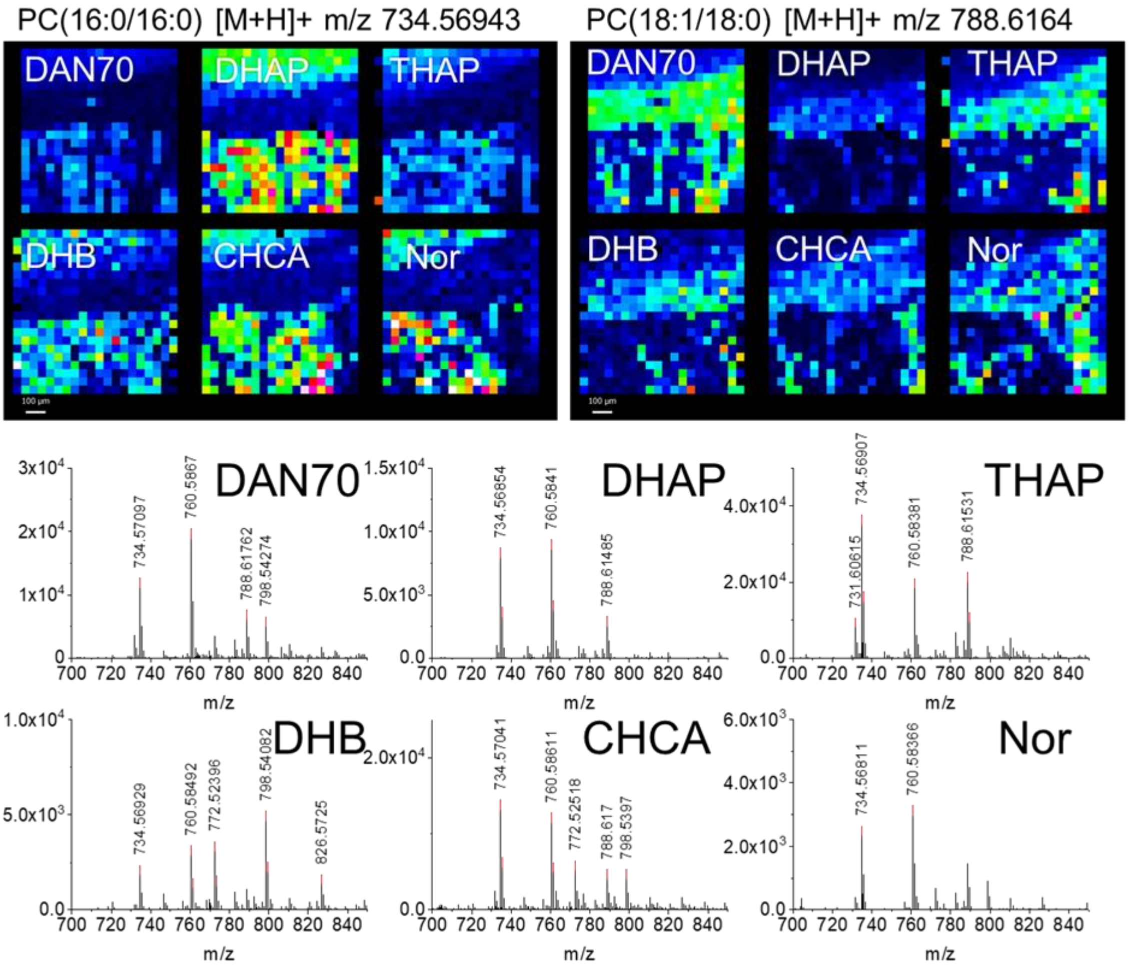

*Figure S 8 Small area fibre tract images with various matrices. Distribution of PC(16:0/16:0) in the grey matter, PC(18:1/18:0) in the white matter and spectra of the lipid region.*
