## supplementary information 2 for "Evaluation of 6 MALDI-Matrices for 10 µm lipid imaging and on-tissue MSn with AP-MALDI-Orbitrap"

### Positive ion mode MS/MS, Identified species

PC(16:0/9:0(OH)) [M+H]+ m/z 650.4392

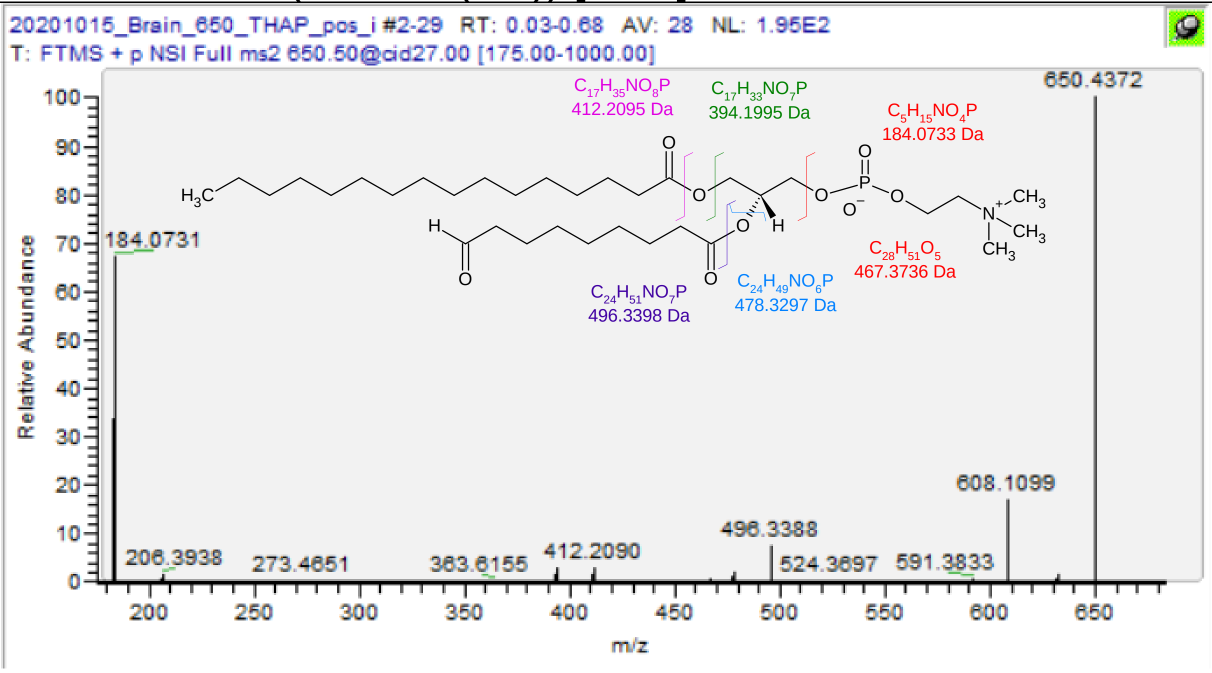

| m/z | [I] | Species | Formula | Da | ppm |
|---|---|---|---|---|---|
| 184.0731 | 69.03 | PC-Hg | C5H15O4NP | 184.0733 | 1.20 |
| 394.1983 | 2.52 | PC(9:0(OH)/-) -H2O | C17H33O7NP | 394.1989 | 1.57 |
| 412.2090 | 3.22 | PC(9:0(OH)/-) | C17H35O8NP | 412.2095 | 1.16 |
| 467.3718 | 0.58 | PC -Hg | C28H51O5 | 467.3731 | 2.78 |
| 478.3282 | 1.82 | PC(16:0/-) -H2O | C24H49O6NP | 478.3292 | 2.09 |
| 496.3388 | 7.52 | PC(16:0/-) | C24H51O7NP | 496.3398 | 1.95 |
| **650.4372** | **100** | **PC(16:0/9:0(OH))** | **C33H65O9NP** | **650.4392** | **3.00** |

| PC(16:0/9:0(COOH)) [M+H]+ m/z 666.4341 |
|---|

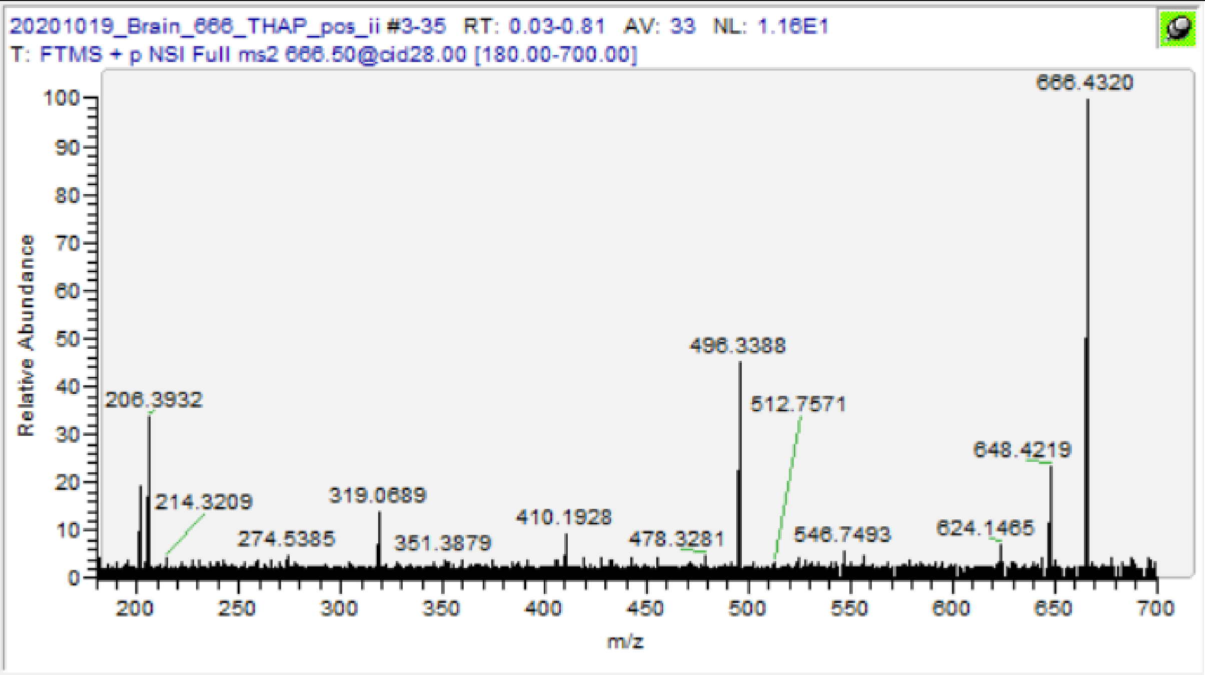

| m/z | [I] | Species | Formula | Da | ppm |
|---|---|---|---|---|---|
| 410.1930 | 8.19 | PC(9:0(COOH)/-) -H2O | C17H33O8NP | 410.1938 | 2.02 |
| 428.2039 | 4.77 | PC(9:0(COOH)/-) | C17H35O9NP | 428.2044 | 1.14 |
| 478.3281 | 5.67 | PC(16:0/-) -H2O | C24H49O6NP | 478.3292 | 2.30 |
| 496.3388 | 52.22 | PC(16:0/-) | C24H51O7NP | 496.3398 | 1.95 |
| 648.4218 | 20.89 | PC -H2O | C33H63O9NP | 648.4235 | 2.62 |
| **666.4321** | **100** | **PC(16:0/9:0(COOH))** | **C33H65O10NP** | **666.4341** | **2.94** |

| PC(18:0/9:0(OH)) [M+H]+ m/z 678.4705 |
|---|

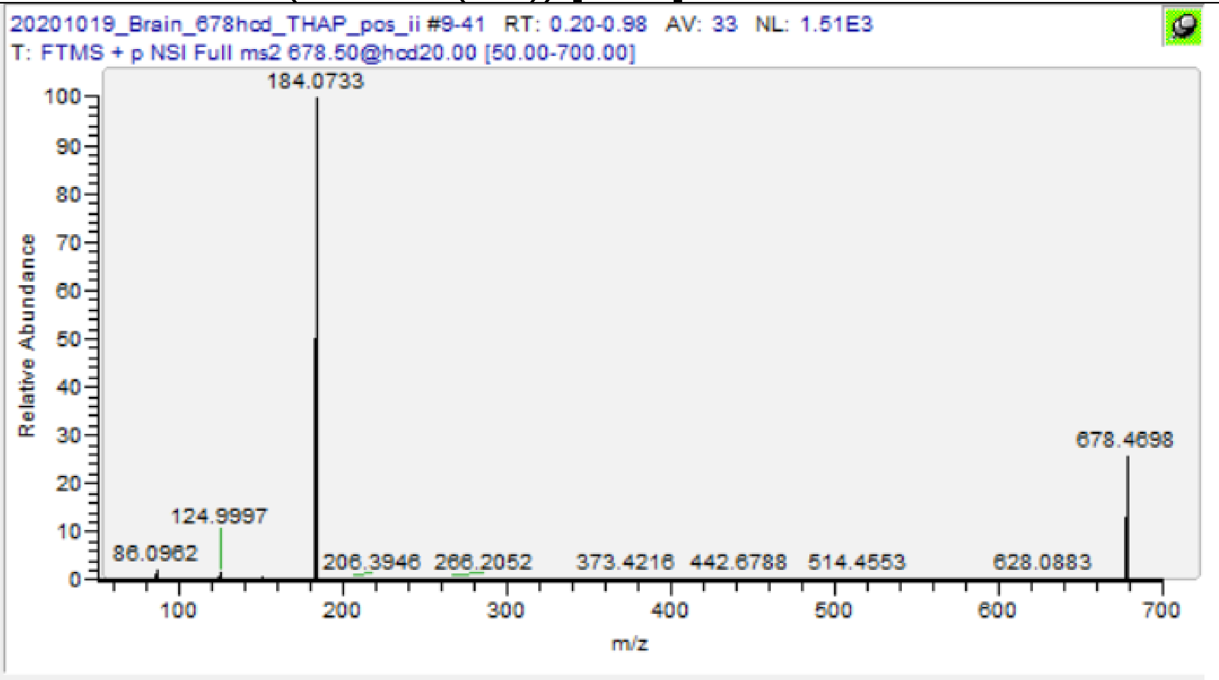

| m/z | [I] | Species | Formula | Da | ppm |
|---|---|---|---|---|---|
| 86.0962 | 2.41 | PC-Frag | C5H12N | 86.09643 | 2.67 |
| 124.9997 | 1.86 | PC-Frag | C2H6O4P | 124.9998 | 0.96 |
| 184.0733 | 100 | PC-Hg | C5H15O4NP | 184.0733 | 0.11 |
| **678.4698** | **23.73** | **PC(18:0/9:0(OH))** | **C35H69O9NP** | **678.4705** | **0.96** |

PC(16:0/16:0)-TMA [M+K]+ m/z 713.4518

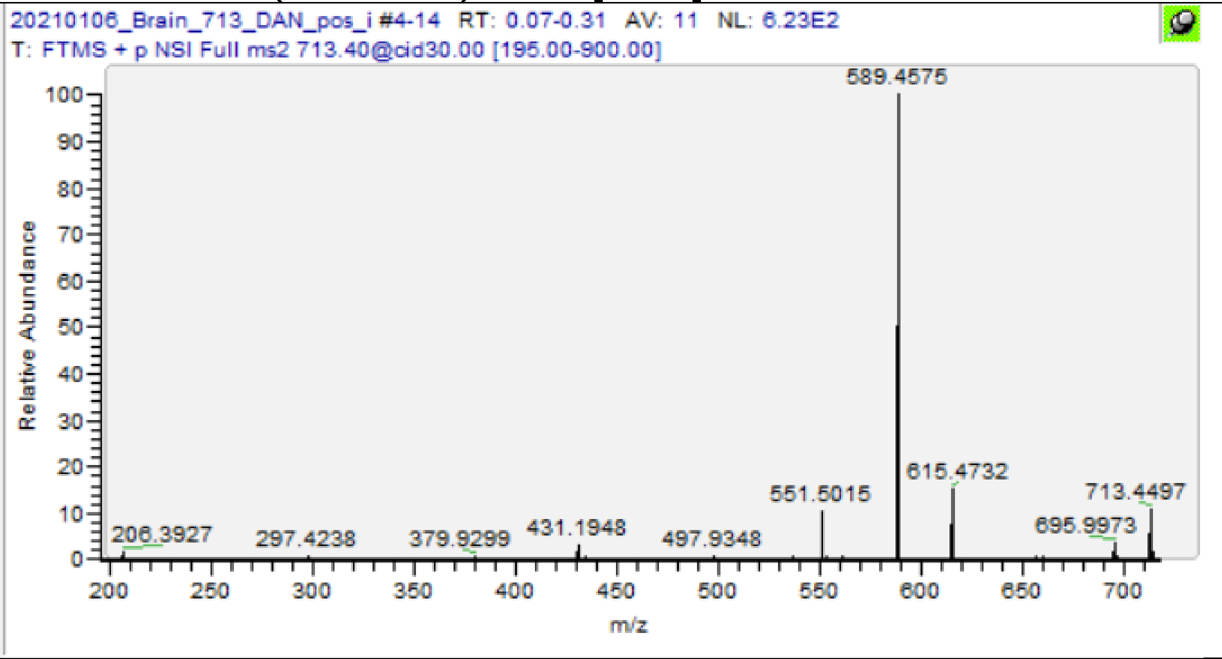

| m/z | [I] | Species | Formula | Da | ppm |
|---|---|---|---|---|---|
| 162.9557 | 100 | PC-Frag +K | C2H5O4PK | 162.9557 | 0 |
| 431.1948 | 3.04 | PA(16:0) +K | C19H37O6PK | 431.1959 | 2.66 |
| 551.5015 | 10.16 | PL(16:0/16:0) -Hg | C35H67O4 | 551.5034 | 3.42 |
| 589.4575 | 100 | PL(16:0/16:0) -Hg +K | C35H66O4K | 589.4593 | 3.00 |
| **713.4497** | **10.79** | **PC(16:0/16:0)-TMA +K** | **C37H71O8PK** | **713.4518** | **2.95** |

PDME(16:0/16:0) [M+H]+ m/z 720.5538

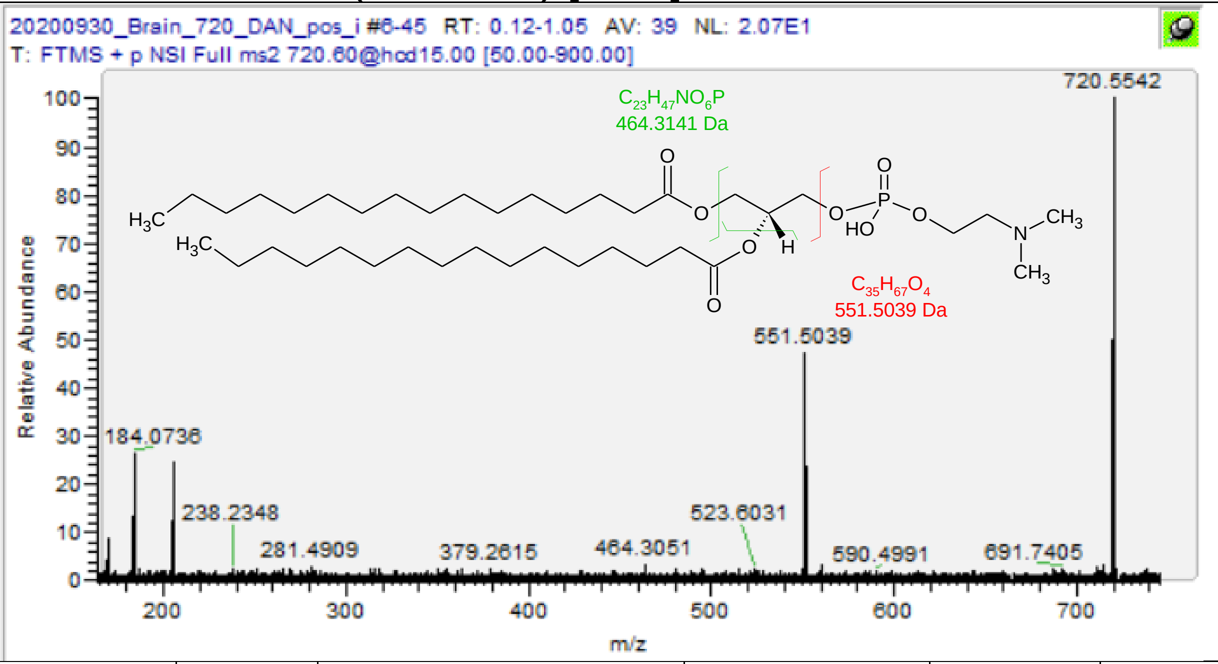

| m/z | [I] | Species | Formula | Da | ppm |
|---|---|---|---|---|---|
| 170.0579 | 11.96 | PDME-Hg | C4H13O4NP | 170.0577 | -1.35 |
| 464.3059 | 3.05 | PDME(16:0/-) | C23H47O6NP | 464.3136 | 16.48 |
| 551.5038 | 55.43 | PDME(16:0/16:0) -Hg | C35H67O4 | 551.5034 | -0.74 |
| **720.5543** | **100** | **PDME(16:0/16:0)** | **C39H79O8NP** | **720.5538** | **-0.72** |
| 184.0736 | 28.46 | PC-Hg | C5H15O4NP | 184.0733 | -1.52 |
| 720.5913 | 49.88 | PC(O-32:0) | C40H83O7NP | | -1.57 |

SM(d36:1) [M+H]+ m/z 731.6062

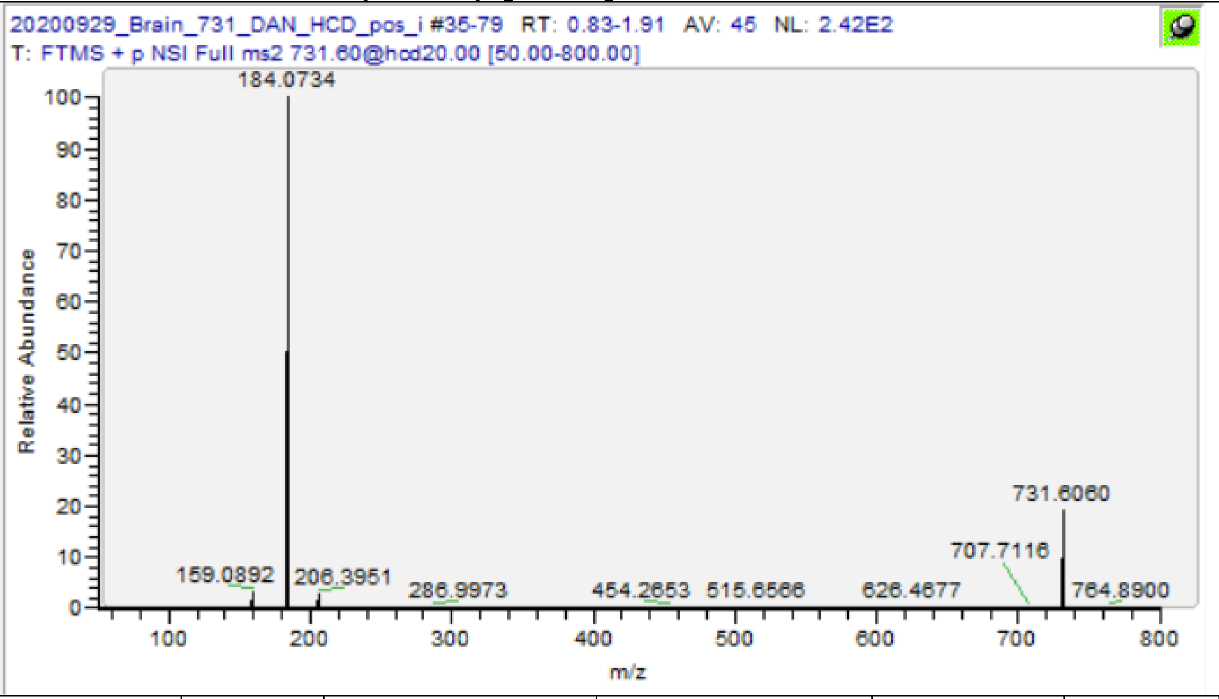

| m/z | [I] | Species | Formula | Da | ppm |
|---|---|---|---|---|---|
| 184.0734 | 100 | SM-Hg | C5H15O4NP | 184.0733 | -0.43 |
| 713.5938 | 24.68 | SM(d36:1) -H2O | C41H82O5N2P | 713.5956 | 2.51 |
| **731.6060** | **27.2** | **SM(d36:1)** | **C41H84O6N2P** | **731.6062** | **0.21** |

PC(16:0/16:0) [M+H]+ m/z 734.56943

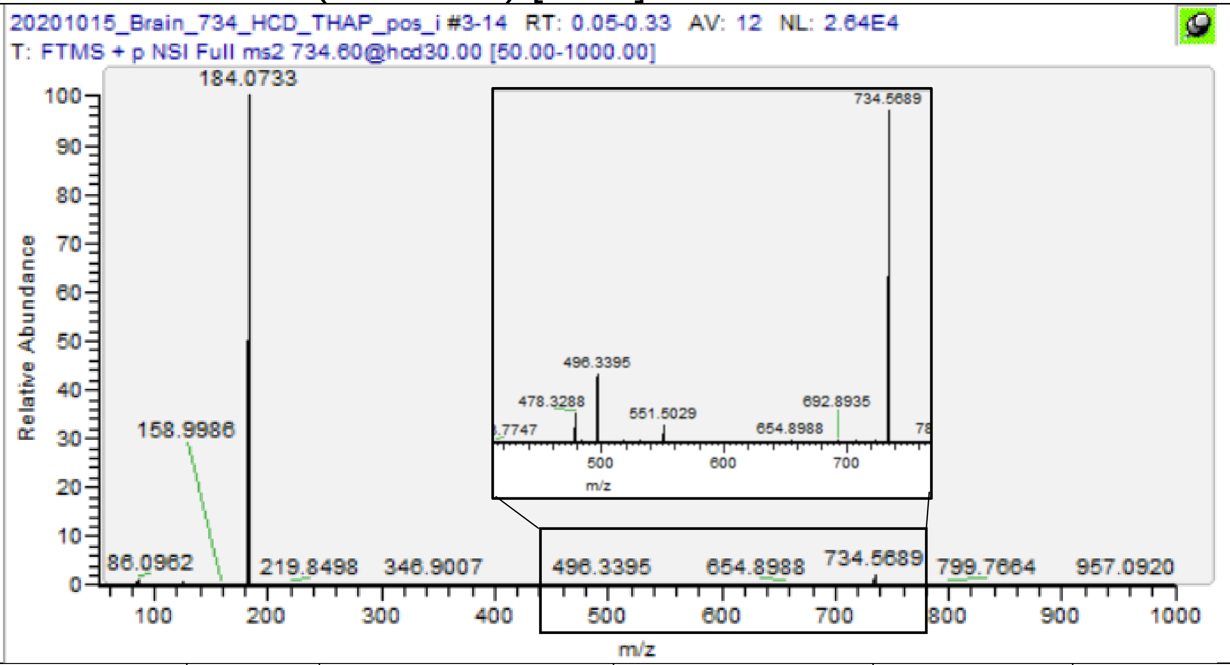

| m/z | [I] | Species | Formula | Da | ppm |
|---|---|---|---|---|---|
| 86.0962 | 0.88 | PC-Frag | C5H12N | 86.09643 | 2.67 |
| 124.9997 | 0.53 | PC-Frag | C2H6O4P | 124.9998 | 0.96 |
| 184.0732 | 100 | PC-Hg | C5H15O4NP | 184.0733 | 0.65 |
| 478.3289 | 0.18 | PC(16:0/-) -H2O | C24H49O6NP | 478.3292 | 0.63 |
| 496.3394 | 0.46 | PC(16:0/-) | C24H51O7NP | 496.3398 | 0.75 |
| 551.5029 | 0.11 | PL(16:0/16:0) -Hg | C35H67O4 | 551.5034 | 0.89 |
| **734.5689** | **2.31** | **PC(16:0/16:0)** | **C40H81O8NP** | **734.56943** | **0.72** |

PC(16:0/16:0) [M+Na]+ m/z 756.5514

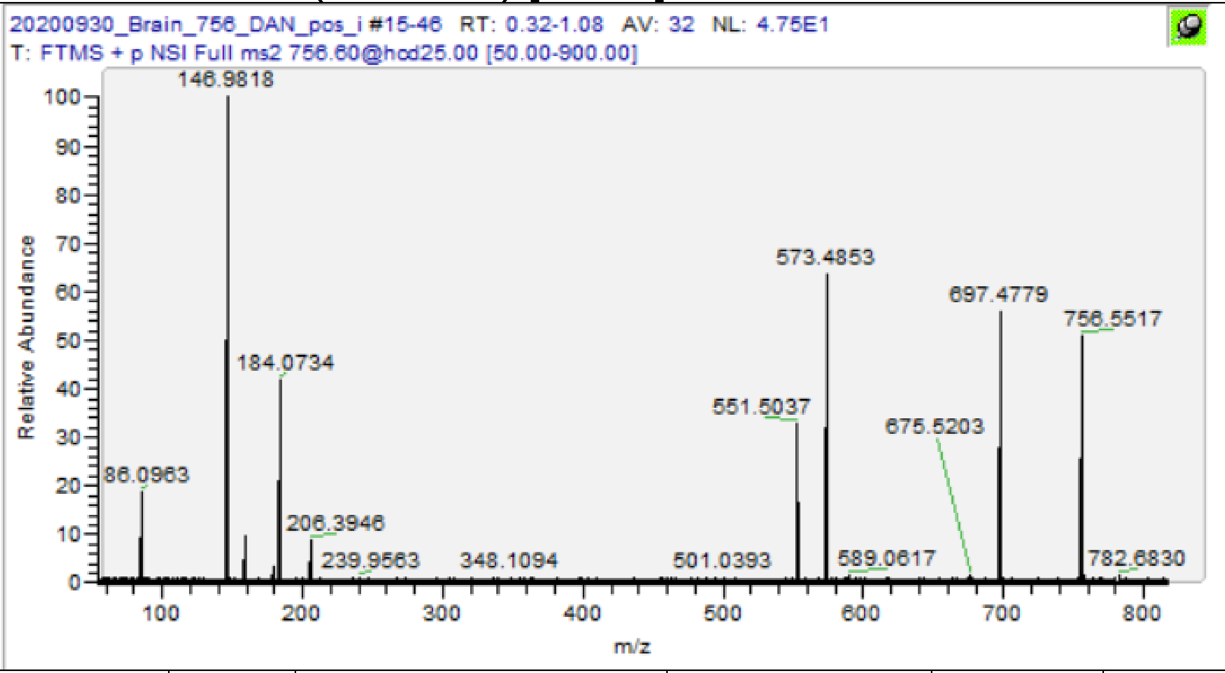

| m/z | [I] | Species | Formula | Da | ppm |
|---|---|---|---|---|---|
| 86.0962 | 16.11 | PC-Frag | C5H12N | 86.09643 | 2.67 |
| 146.9818 | 100 | PC-Frag+Na | C2H5O4PNa | 146.9818 | -0.20 |
| 184.0734 | 60.35 | PC-Hg | C5H15O4NP | 184.0733 | -0.43 |
| 551.5042 | 31.31 | PL(16:0/16:0) -Hg | C35H67O4 | 551.5034 | -1.47 |
| 573.4849 | 84.93 | PL(16:0/16:0) -Hg +Na | C35H66O4Na | 573.4853 | 0.75 |
| 697.4789 | 54.74 | PC(16:0/16:0)-TMA +Na | C37H71O8PNa | 697.4779 | -1.46 |
| **756.5530** | **47.36** | **PC(16:0/16:0)+Na** | **C40H80O8NPNa** | **756.5514** | **-2.14** |

# PC(16:0/18:1) [M+H]+ m/z 760.5851

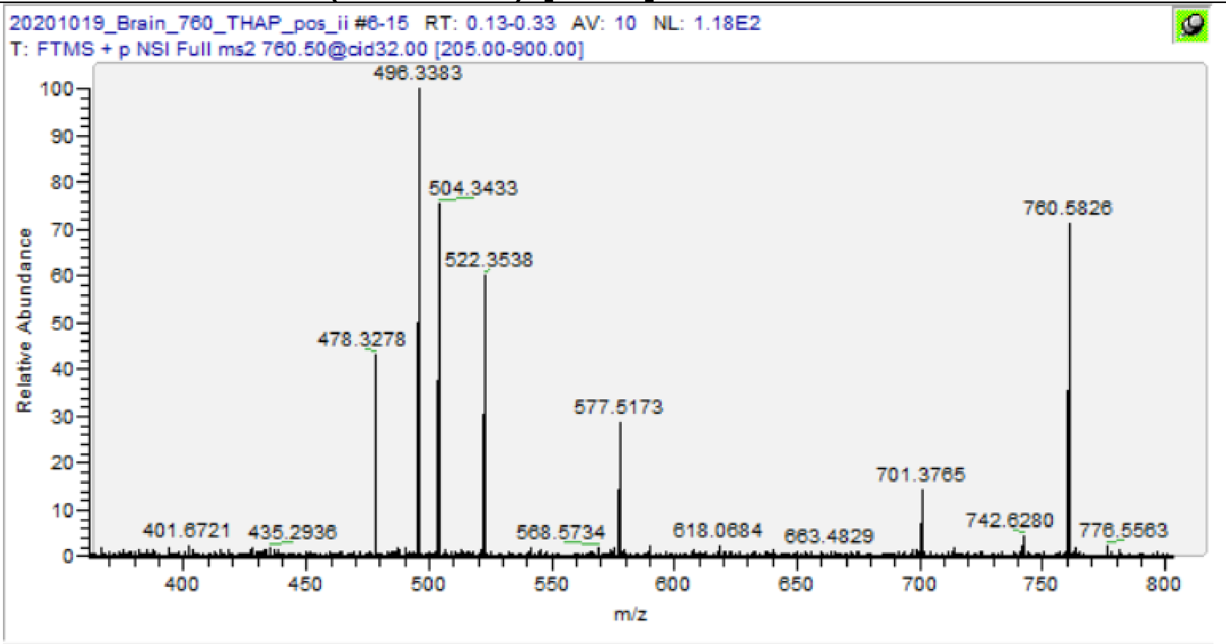

| m/z | [I] | Species | Formula | Da | ppm |
|---|---|---|---|---|---|
| 478.3279 | 45.02 | PC(16:0/-) -H2O | C24H49O6NP | 478.3292 | 2.72 |
| 496.3383 | 100 | PC(16:0/-) | C24H51O7NP | 496.3398 | 2.96 |
| 504.3433 | 74.77 | PC(18:1/-) –H2O | C26H51O6NP | 504.3449 | 3.07 |
| 522.3538 | 58.46 | PC(18:1/-) | C26H53O7NP | 522.3554 | 3.10 |
| 577.5174 | 30.31 | PL(16:0/18:1)-Hg | C37H69O4 | 577.519 | 2.84 |
| 701.3765 | 13.7 | PC(16:0/18:1) -TMA | C39H74O8P | 701.5116 | 192.56 |
| **760.5826** | **68.52** | **PC(16:0/18:1)** | **C42H83O8NP** | **760.5851** | **3.26** |

PC(16:0/16:0)* [M+K]+ m/z 772.5253

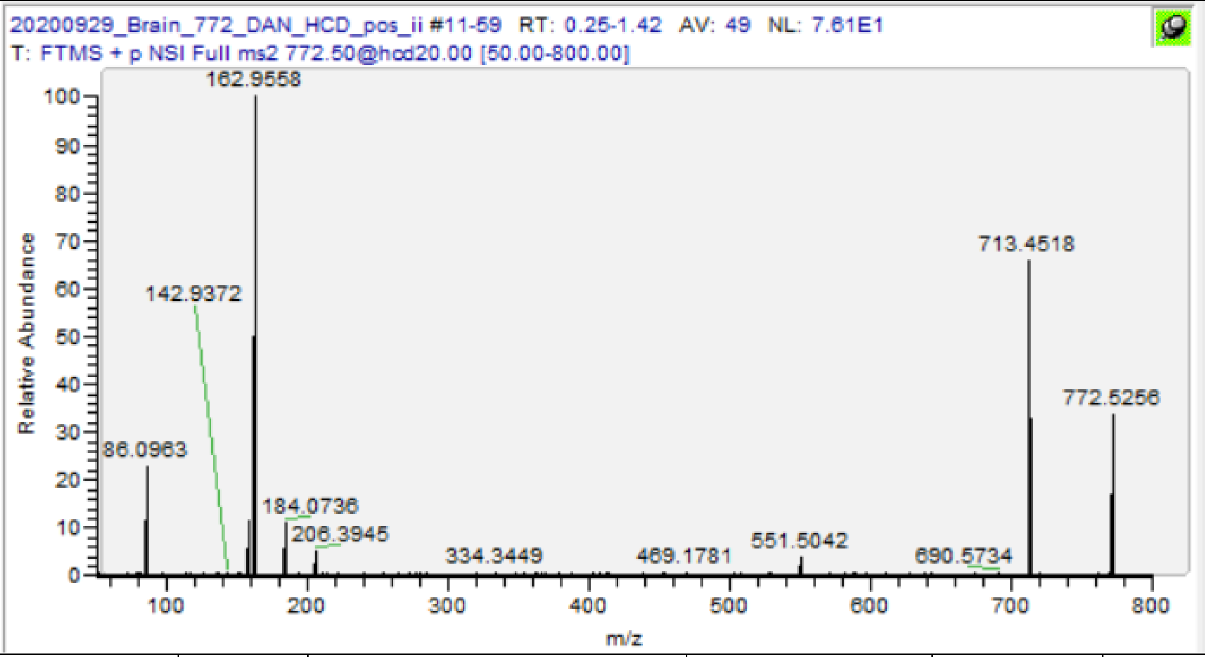

| m/z | [I] | Species | Formula | Da | ppm |
|---|---|---|---|---|---|
| 86.0963 | 24.16 | PC-Frag | C5H12N | 86.09643 | 1.51 |
| 162.9558 | 100 | PC-Frag +K | C2H5O4PK | 162.9557 | -0.61 |
| 184.0735 | 11.33 | PC-Hg | C5H15O4NP | 184.0733 | -0.98 |
| 551.5043 | 2.74 | PL(16:0/16:0) -Hg | C35H67O4 | 551.5034 | -1.65 |
| 589.4594 | 0.71 | PL(16:0/16:0) -Hg +K | C35H66O4K | 589.4593 | -0.22 |
| 713.4518 | 76.03 | PC(16:0/16:0)-TMA +K | C37H71O8PNa | 713.4518 | 0.01 |
| **772.5254** | **41.02** | **PC(16:0/16:0)+K** | **C40H80O8NPK** | **772.5253** | **-0.12** |

PC(16:0/18:1)* [M+Na]+ m/z 782.5670

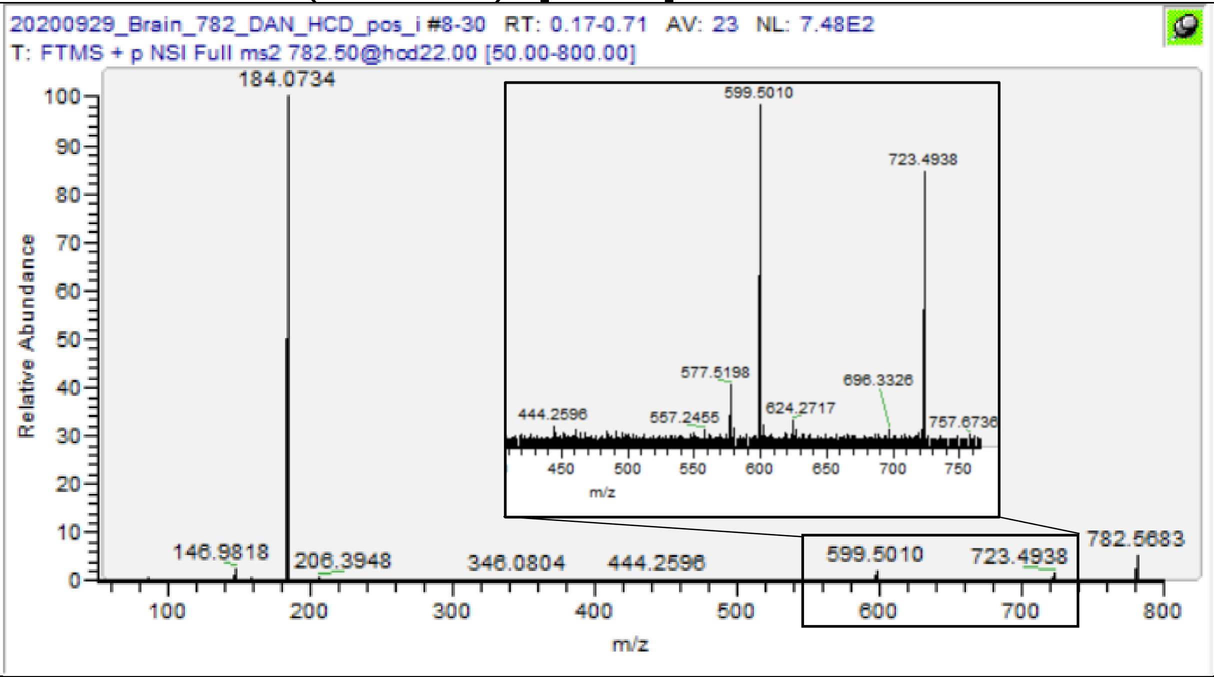

| m/z | [I] | Species | Formula | Da | ppm |
|---|---|---|---|---|---|
| 86.0962 | 0.5 | PC-Frag | C5H12N | 86.09643 | 2.67 |
| 184.0734 | 100 | PC-Hg | C5H15O4NP | 184.0733 | -0.43 |
| 577.5200 | 0.36 | PL(16:0/18:1)-Hg | C37H69O4 | 577.519 | -1.66 |
| 599.5006 | 1.42 | PL(16:0/18:1)-Hg +Na | C37H69O4Na | 599.501 | 0.63 |
| 723.4939 | 1.93 | PC(16:0/18:1)-TMA +Na | C39H73O8PNa | 723.4935 | -0.51 |
| **782.5685** | **8.33** | **PC(16:0/18:1) +Na** | **C42H82O8NP Na** | **782.567** | **-1.88** |

### HexCer(d18:1/22:1) [M+H]+ m/z 782.6505

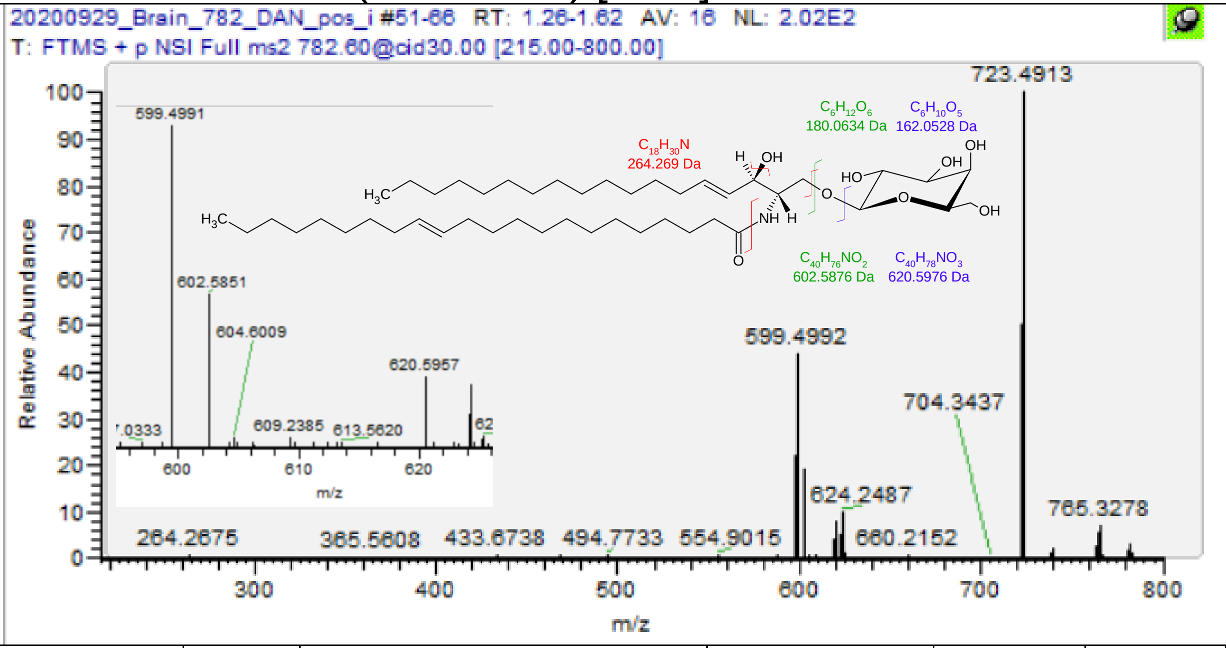

| m/z | [I] | Species | Formula | Da | ppm |
|---|---|---|---|---|---|
| 264.2678 | 0.59 | Cer d18:1 Frag. | C18H34N | 264.2686 | 2.95 |
| 602.5853 | 16.38 | HexCer(d18:1/22:1) -Glc -H2O | C40H76O2N | 602.5871 | 2.92 |
| 620.5959 | 7.77 | HexCer(d18:1/22:1) -Glc | C40H78O3N | 620.5976 | 2.77 |
| **782.6489** | **3.04** | **HexCer(d18:1/22:1)** | **C46H88O8N** | **782.6505** | **1.98** |

SM(d40:2) [M+H]+ m/z 785.6531

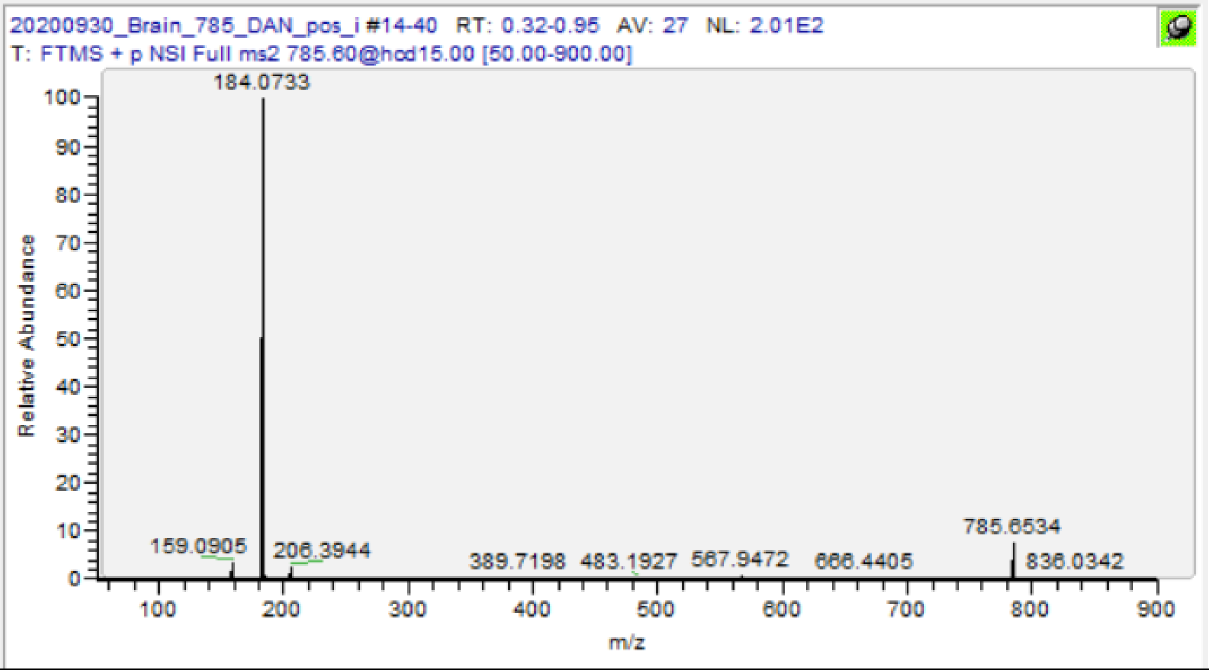

| m/z | [I] | Species | Formula | Da | ppm |
|---|---|---|---|---|---|
| 184.0733 | 100 | PC-Hg | C5H15O4NP | 184.0733 | 0.11 |
| 767.6404 | 9.88 | SM(d40:2) -H2O | C45H88O5N2P | 767.6425 | 2.79 |
| **785.6534** | **7.56** | **SM(d40:2)** | **C45H90O6N2P** | **785.6531** | **-0.38** |

PC(18:1/18:1) [M+H]+ m/z 786.6007

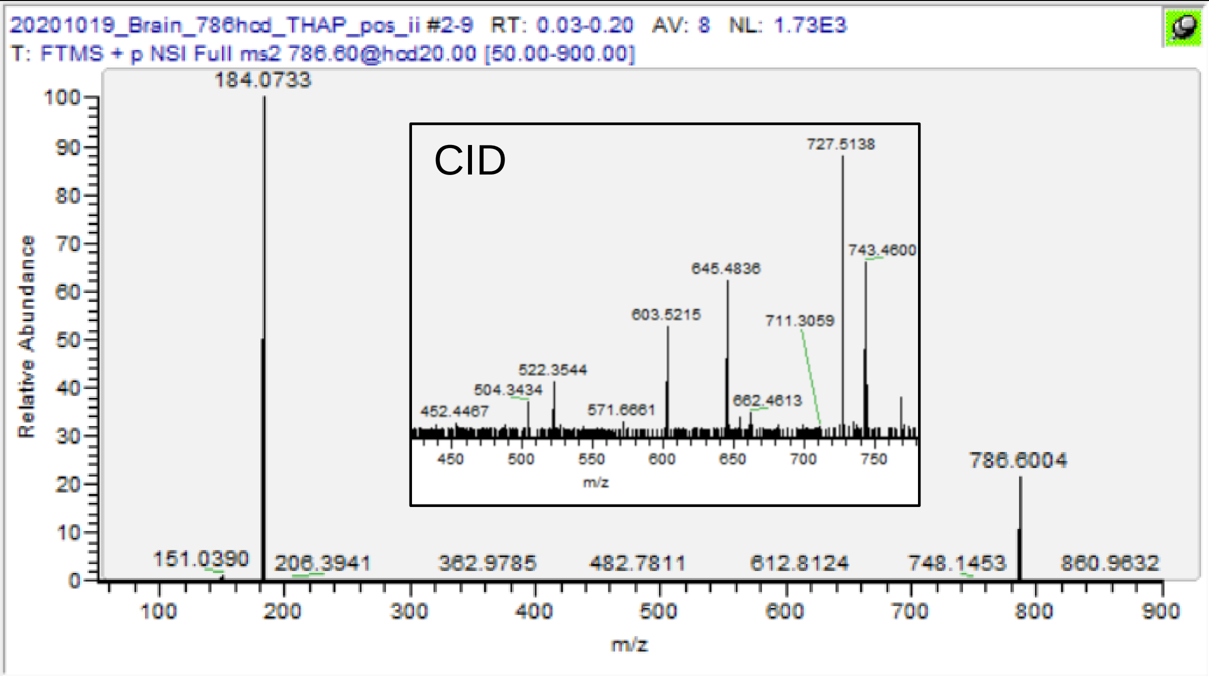

| m/z | [I] | Species | Formula | Da | ppm |
|---|---|---|---|---|---|
| 184.0733 | 100 | PC-Hg | C5H15O4NP | 184.0733 | 0.11 |
| 504.3435 | 9.53 | PC(18:1/-) –H2O | C26H51O6NP | 504.3449 | 2.68 |
| 522.3542 | 17.81 | PC(18:1/-) | C26H53O7NP | 522.3554 | 2.34 |
| 603.5216 | 29.3 | PL(18:1/18:1) -Hg | C39H71O4 | 603.5347 | 21.69 |
| 726.5106 | 4.42 | PC(18:1/18:1) -TMA | C41H75O8P | 726.5194 | 12.13 |
| **786.6004** | **21.45** | **PC(18:1/18:1)** | **C44H85O8NP** | **786.6007** | **0.42** |

PC(18:1/18:0) [M+H]+ m/z 788.6164

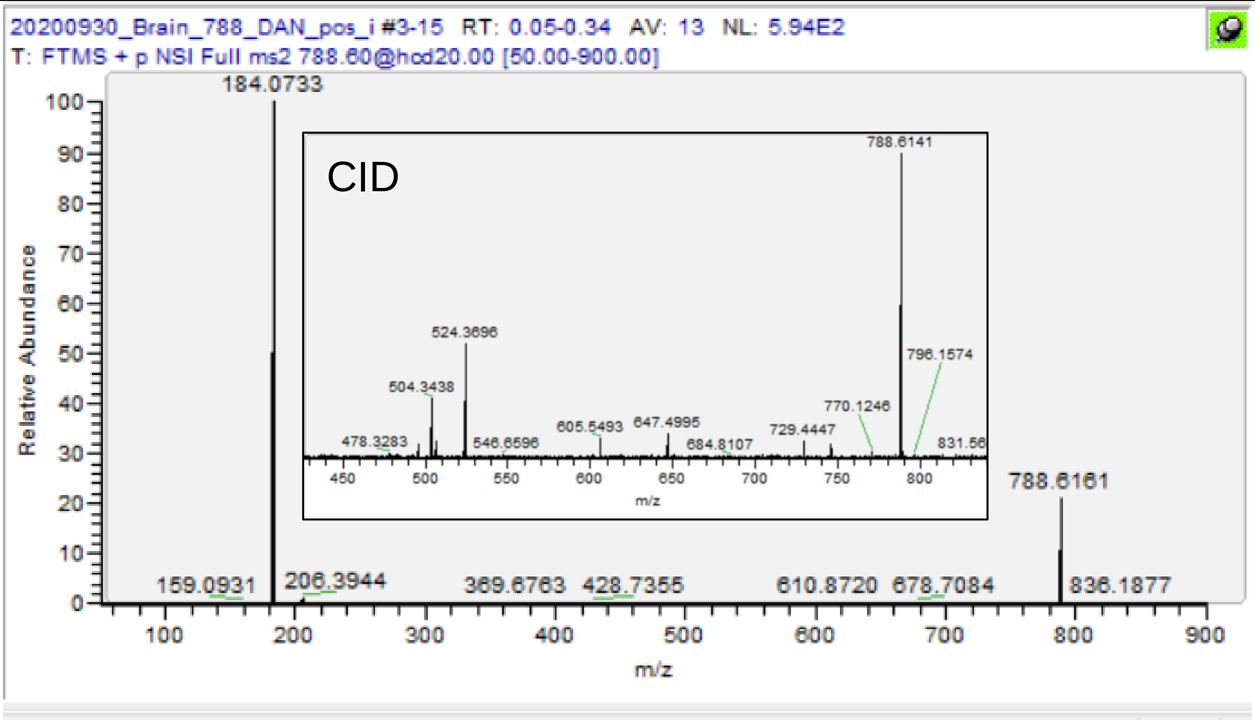

| m/z | [I] | Species | Formula | Da | ppm |
|---|---|---|---|---|---|
| 184.0734 | 100 | PC-Hg | C5H15O4NP | 184.0733 | -0.43 |
| 504.3436 | 18.46 | PC(18:1/-) –H2O | C26H51O6NP | 504.3449 | 2.48 |
| 506.3595 | 5.21 | PC(18:0/-) –H2O | C26H53O6NP | 506.3605 | 1.97 |
| 522.3544 | 2 | PC(18:1/-) | C26H53O7NP | 522.3554 | 1.95 |
| 524.3694 | 28.73 | PC(18:0/-) | C26H55O7NP | 524.3711 | 3.18 |
| 605.5483 | 1.49 | PL(18:1/18:0) -Hg | C39H73O4 | 605.5503 | 3.37 |
| **788.6161** | **21.36** | **PC(18:1/18:0)** | **C44H87O8NP** | **788.6164** | **0.36** |

| PE(40:7)* [M+H]+ m/z 790.5381 | | | | | |
|---|---|---|---|---|---|
| 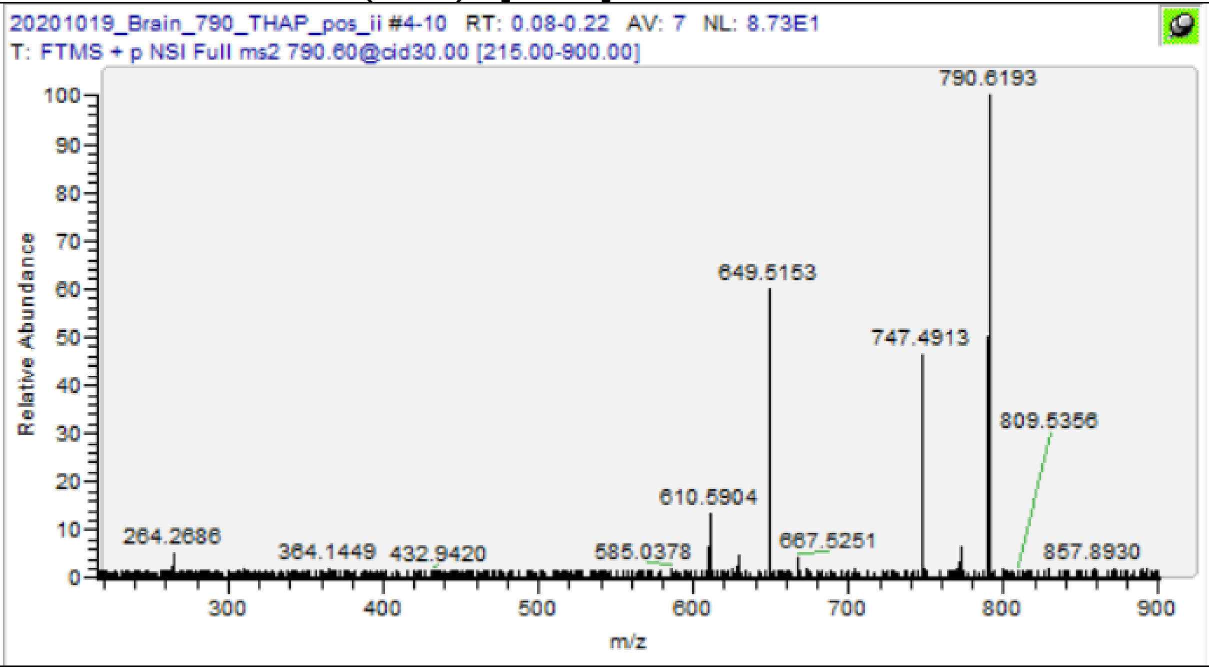 | | | | | |
| m/z | [I] | Species | Formula | Da | ppm |
| 649.5186 | 0.36 | PL(40:7) -Hg | C43H69O4 | 649.519 | 0.68 |
| 747.4914 | 59.81 | PA(40:7) | C43H72O8P | 747.4959 | 6.06 |
| **790.5357** | **1.1** | **PE(40:7)** | **C45H77O8NP** | **790.5381** | **3.07** |

PE(18:0/22:6) [M+H]+ m/z 792.5538

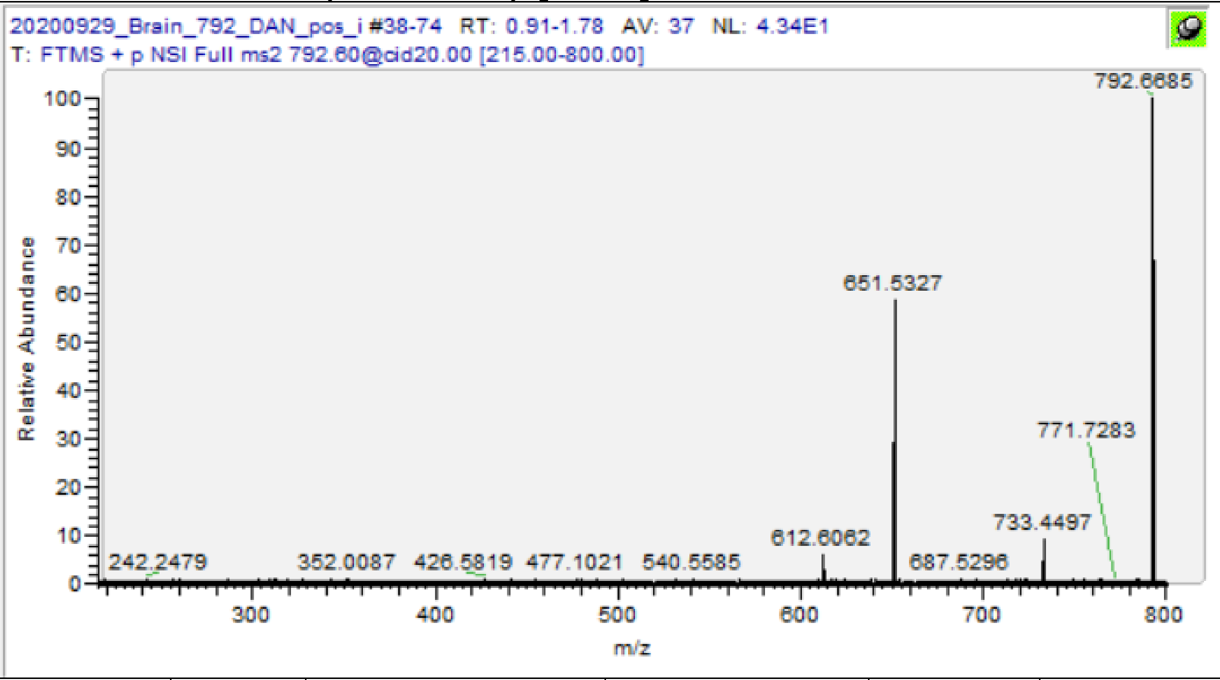

| m/z | [I] | Species | Formula | Da | ppm |
|---|---|---|---|---|---|
| 341.3045 | 1.29 | PL(18:0)-frag | C21H41O3 | 341.305 | 1.52 |
| 651.5344 | 59.55 | PL(18:0/22:6) -Hg | C43H71O4 | 651.5347 | 0.45 |
| **792.5555** | **16.86** | **PE(18:0/22:6)** | **C45H79O8NP** | **792.55378** | **-2.17** |

PC(16:0/18:1)* [M+K]+ m/z 798.541

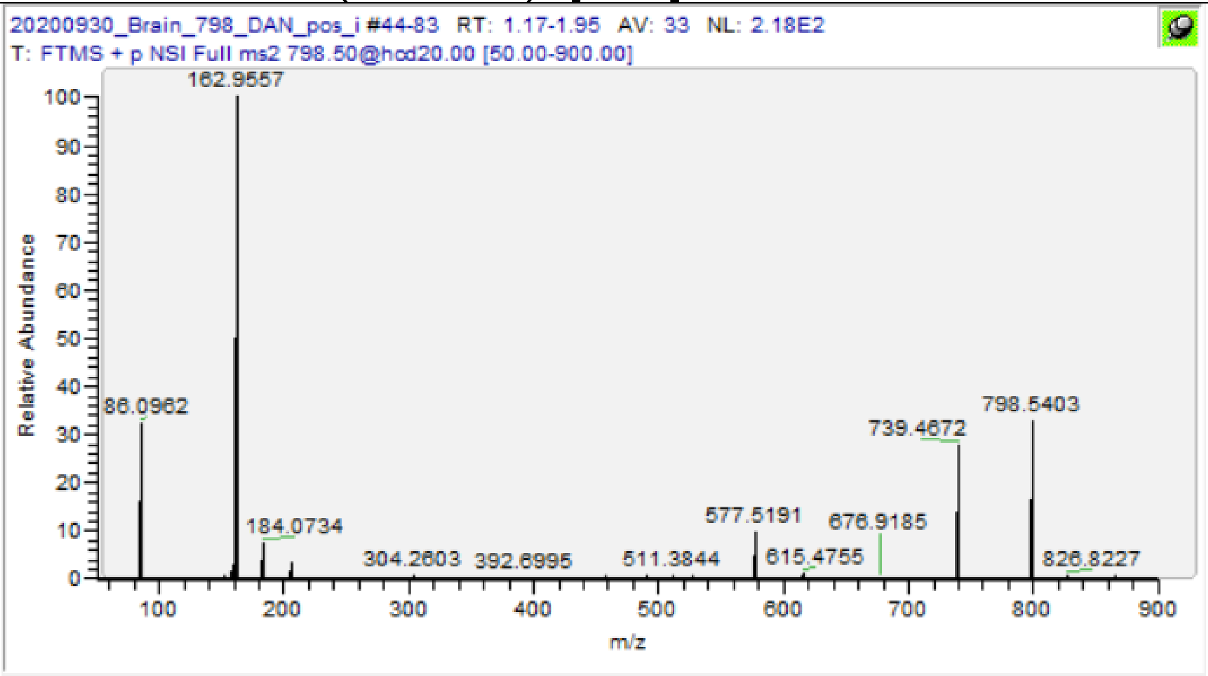

| m/z | [I] | Species | Formula | Da | ppm |
|---|---|---|---|---|---|
| 86.0962 | 31.94 | PC-Frag | C5H12N | 86.09643 | 2.67 |
| 162.9557 | 100 | PC-Frag +K | C2H5O4PK | 162.9557 | 0.00 |
| 184.0734 | 7.49 | PC-Hg | C5H15O4NP | 184.0733 | -0.43 |
| 577.5191 | 9.62 | PL(16:0/18:1) -Hg | C37H69O4 | 577.519 | -0.10 |
| 615.4755 | 1.25 | PL(16:0/18:1) -Hg +K | C37H68O4K | 615.4749 | -0.94 |
| 739.4672 | 29.96 | PC(16:0/18:1) -TMA +K | C39H73O8PK | 739.4675 | 0.35 |
| **798.5403** | **36.79** | **PC(16:0/18:1) +K** | **C42H82O8NPK** | **798.541** | **0.83** |

# PC(16:0/22:6) [M+H]+ m/z 806.5694

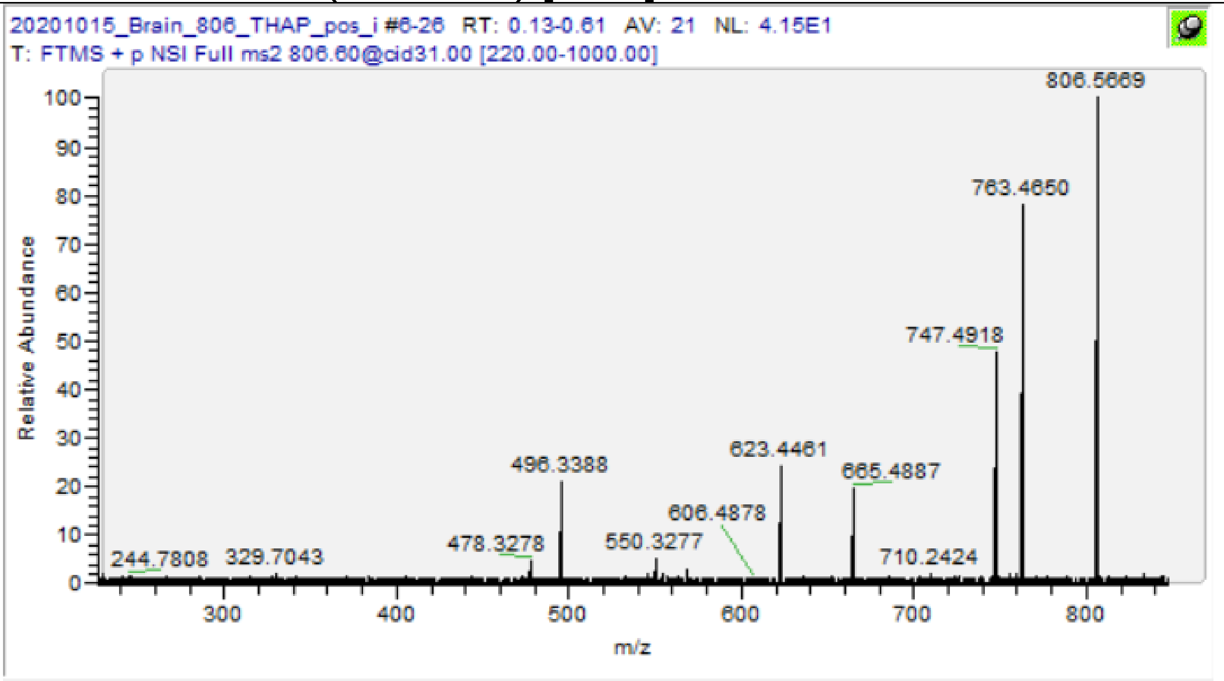

| m/z | [I] | Species | Formula | Da | ppm |
|---|---|---|---|---|---|
| 86.0962 | 2.72 | PC-Frag | C5H12N | 86.09643 | 2.67 |
| 124.9997 | 1.95 | PC-Frag | C2H6O4P | 124.9998 | 0.96 |
| 184.0733 | 100 | PC-Hg | C5H15O4NP | 184.0733 | 0.11 |
| 478.3278 | 2.96 | PC(16:0/-) -H2O | C24H49O6NP | 478.3292 | 2.93 |
| 496.3388 | 19 | PC(16:0/-) | C24H51O7NP | 496.3398 | 1.95 |
| 550.3277 | 5.52 | PC(22:6/-) -H2O | C30H49O6NP | 550.3292 | 2.73 |
| 568.3387 | 3.85 | PC(22:6/-) | C30H51O7NP | 568.3398 | 1.88 |
| 623.5001 | 19.24 | PL(16:0/22:6) -Hg | C41H67O4 | 623.5034 | 5.28 |
| 747.4920 | 45.28 | PC(16:0/22:6) -TMA | C43H72O8P | 747.4959 | 5.26 |
| **806.5691** | **5.47** | **PC(16:0/22:6)** | **C46H81O8NP** | **806.5694** | **0.41** |

| PE(38:4) [M+K]+ m/z 806.5097 |
|---|

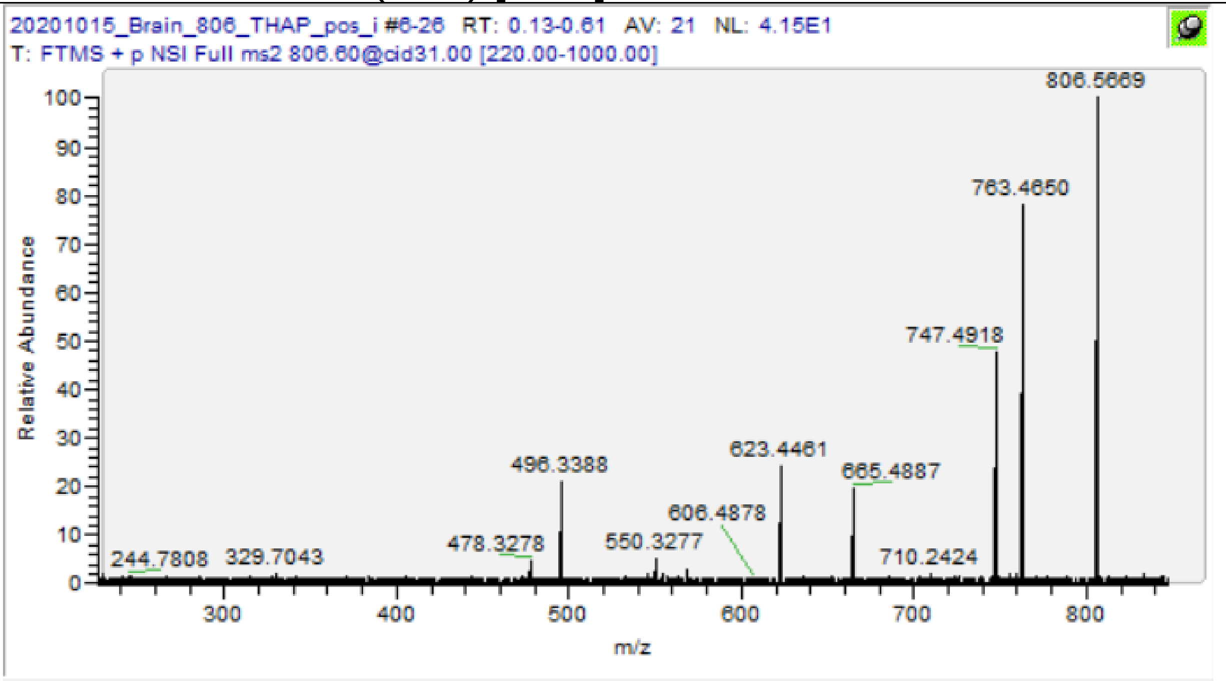

| m/z | [I] | Species | Formula | Da | ppm |
|---|---|---|---|---|---|
| 665.4884 | 21.08 | PL(38:4) -Hg +K | C41H70O4K | 665.4906 | 3.26 |
| 763.4652 | 72.64 | PA(38:4) +K | C41H73O8PK | 763.4675 | 2.96 |
| **806.5067** | **21.8** | **PE(38:4) +K** | **C43H78NO8PK** | **806.5097** | **3.67** |

PC(18:1/20:4) [M+H]+ m/z 808.5851

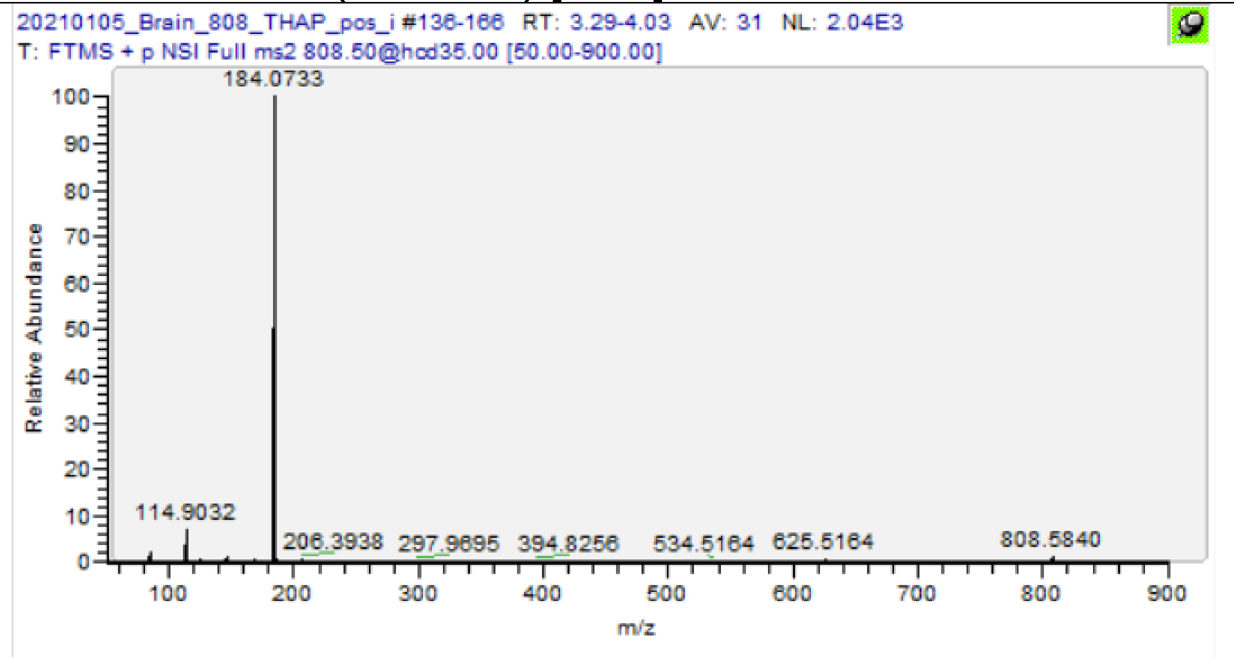

| m/z | [I] | Species | Formula | Da | ppm |
|---|---|---|---|---|---|
| 184.0733 | 100 | PC-Hg | C5H15O4NP | 184.0733 | 0.11 |
| 522.3546 | 0.05 | PC(18:1,–) | C26H53O7NP | 522.3554 | 1.56 |
| 625.5164 | 0.55 | PC(18:1/20:4) -Hg | C41H69O4 | 625.519 | 4.22 |
| 749.5094 | 0.12 | PC(18:1/20:4) -TMA | C43H74O8P | 749.5116 | 2.90 |
| **808.5839** | **0.93** | **PC(18:1/20:4)** | **C46H83O8NP** | **808.5851** | **1.45** |

HexCer(d18:1/24:2) [M+H]+ m/z 808.6661

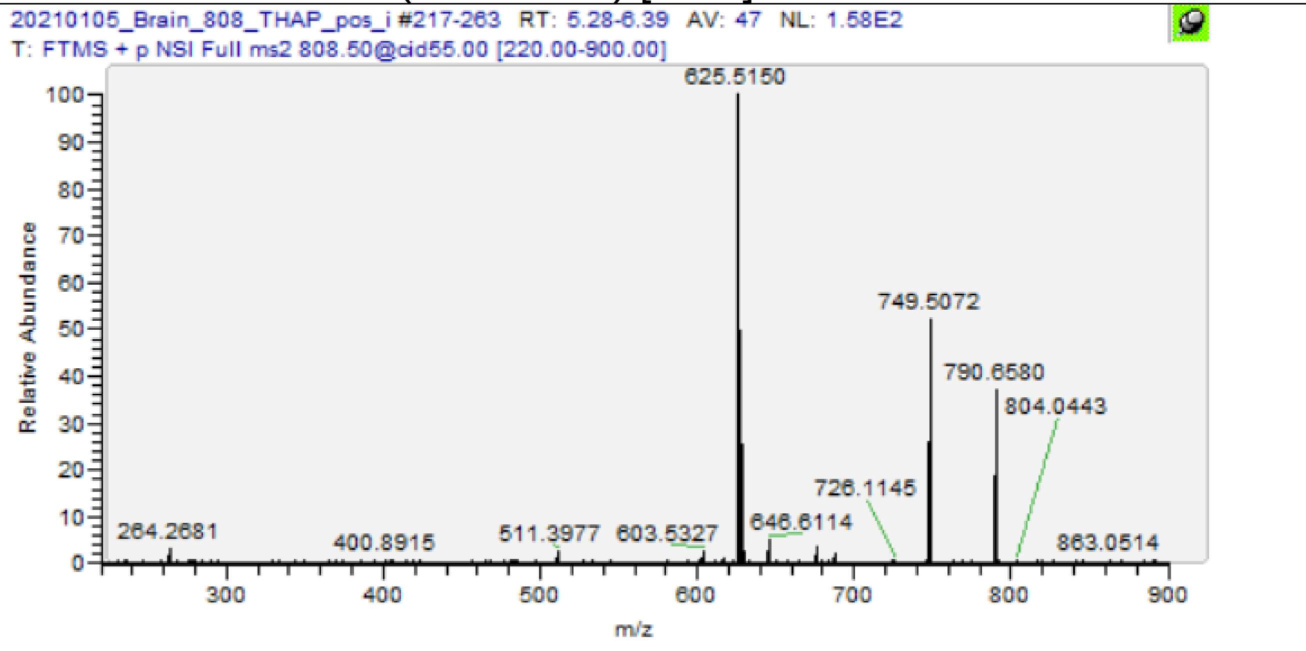

| m/z | [I] | Species | Formula | Da | ppm |
|---|---|---|---|---|---|
| 264.2685 | 0.7 | Cer d18:1 Frag. | C18H34N | 264.2686 | 0.30 |
| 628.601 | 6.07 | HexCer(d18:1/24:2) -Glc -H2O | C42H78O2N | 628.6027 | 2.72 |
| 646.6114 | 1.59 | HexCer(d18:1/24:2) -Glc | C42H80O3N | 646.6133 | 2.89 |
| 790.658 | 21.41 | HexCer(d18:1/24:2) -H2O | C48H88O7N | 790.6555 | -3.12 |
| **808.6642** | **1.22** | **HexCer(d18:1/24:2)** | **C48H90O8N** | **808.6661** | **2.34** |

SM(d42:2) [M+H]+ m/z 813.6844

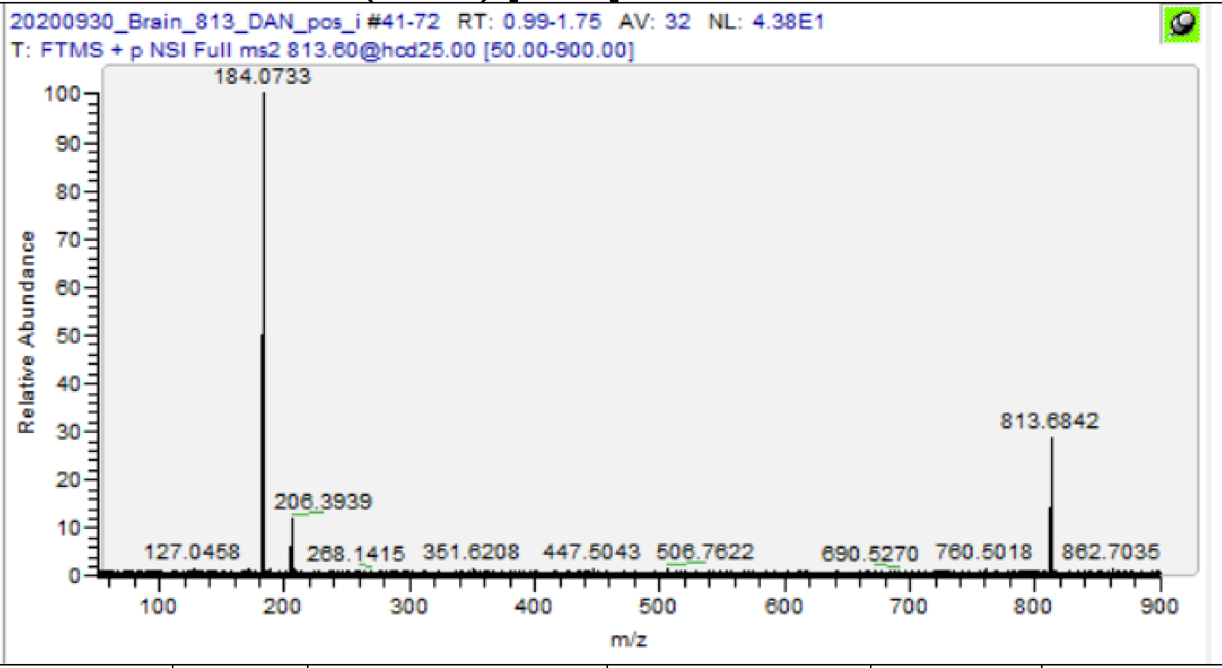

| m/z | [I] | Species | Formula | Da | ppm |
|---|---|---|---|---|---|
| 184.0733 | 100 | PC-Hg | C5H15O4NP | 184.0733 | 0.11 |
| **813.6846** | **27.74** | **SM (d42:2)** | **C47H94O6N2P** | **813.6844** | **-0.25** |

PC(18:1/18:0)* [M+K]+ m/z 826.5723

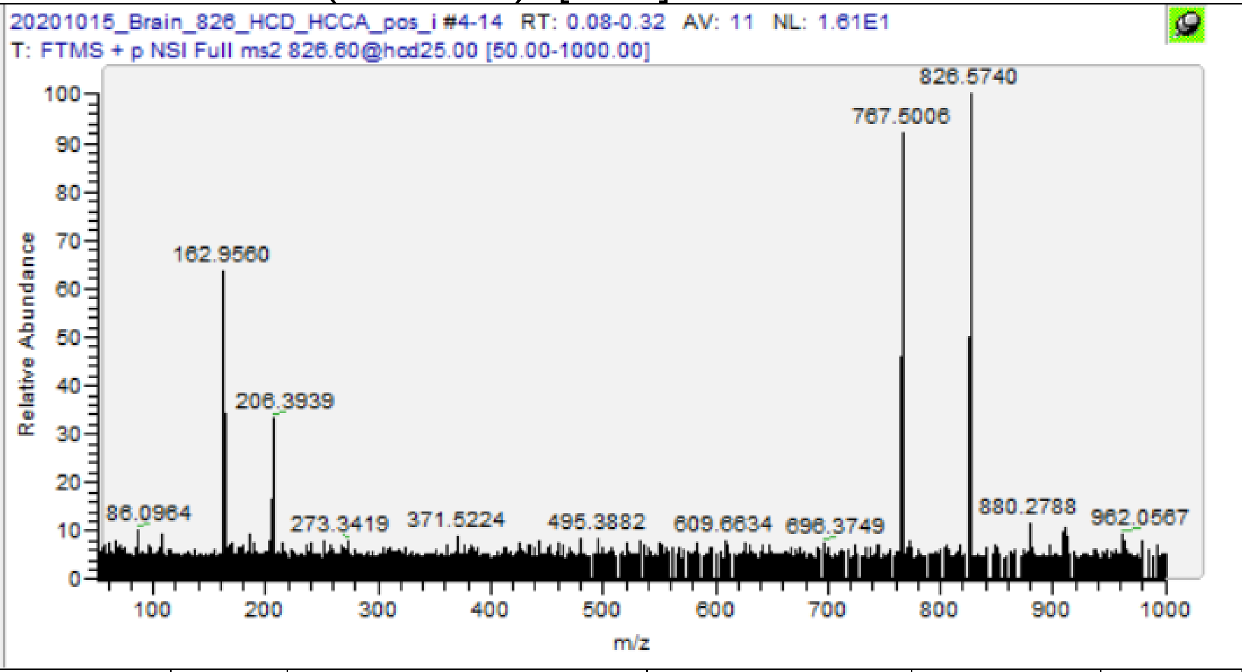

| m/z | [I] | Species | Formula | Da | ppm |
|---|---|---|---|---|---|
| 86.0964 | 8.75 | PC-Frag | C5H12N | 86.09643 | 0.35 |
| 162.9560 | 55.72 | PC-Frag +K | C2H5O4PK | 162.9557 | -1.84 |
| 643.5067 | 4.04 | PL(18:1/18:0) -Hg +K | C39H72O4K | 643.5062 | -0.75 |
| 767.4998 | 85.7 | PC(18:1/18:0) -TMA +K | C41H77O8PK | 767.4988 | -1.36 |
| **826.5739** | **100** | **PC(18:1/18:0) +K** | **C44H86O8NPK** | **826.5723** | **-1.98** |

PC(18:0/22:6) [M+H]+ m/z 834.6007

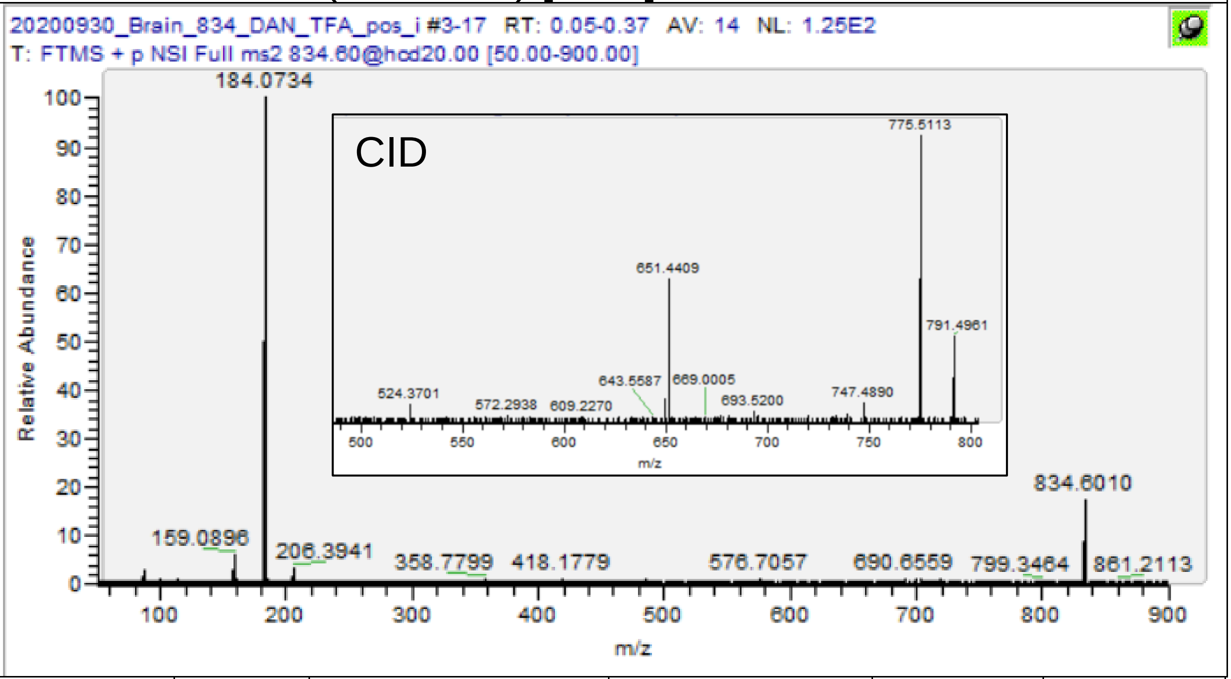

| m/z | [I] | Species | Formula | Da | ppm |
|---|---|---|---|---|---|
| 184.0734 | 100 | PC-Hg | C5H15O4NP | 184.0733 | -0.43 |
| 524.3701 | 1.25 | PC(18:0,–) | C26H55O7NP | 524.3711 | 1.84 |
| 651.5218 | 6.72 | PC(18:0/22:6) -Hg | C43H71O4 | 651.5347 | 19.78 |
| 775.5115 | 23.48 | PC(18:0/22:6) -TMA | C45H76O8P | 775.5272 | 20.282 |
| **834.6012** | **16.55** | **PC(18:0/22:6)** | **C48H85O8NP** | **834.6007** | **-0.56** |

PC(38:6) [M+K]+ m/z 844.5253

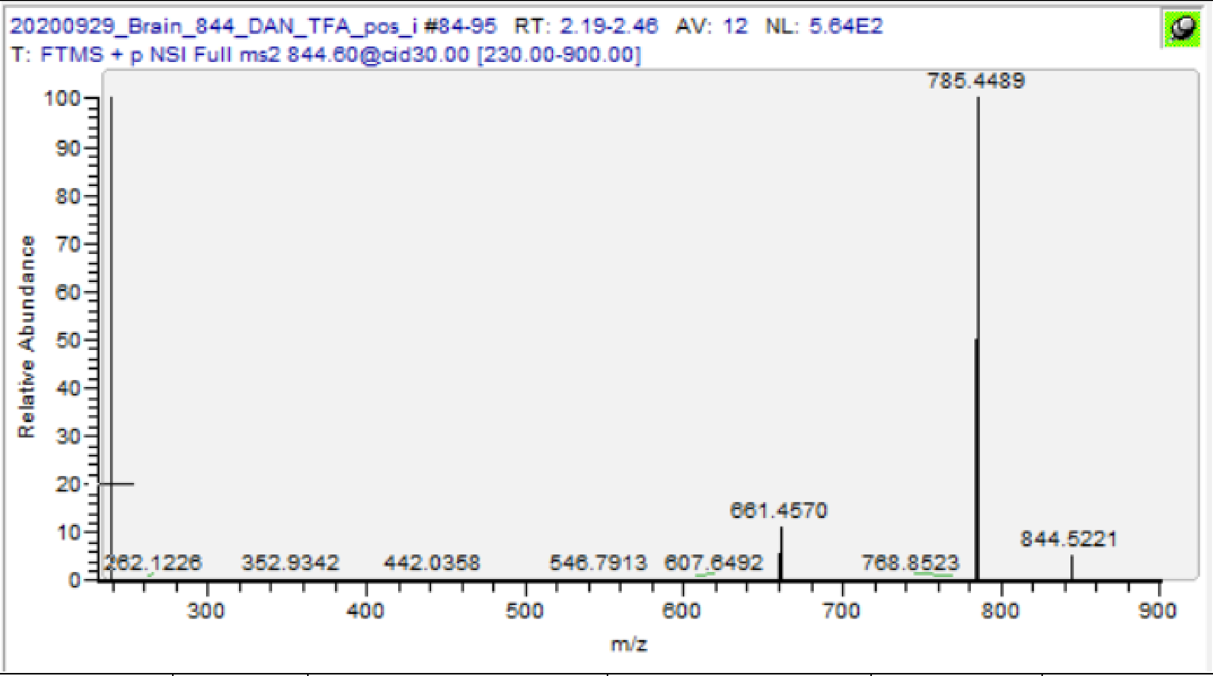

| m/z | [I] | Species | Formula | Da | ppm |
|---|---|---|---|---|---|
| 661.4575 | 11.35 | PL(38:6) -Hg +K | C41H66O4K | 661.4593 | 2.68 |
| 785.4495 | 100 | PC(38:6) -TMA +K | C43H71O8PK | 785.4518 | 2.94 |
| **844.5228** | **21.36** | **PC(38:6) +K** | **C46H80NO8PK** | **844.5253** | **2.97** |

PC(38:4) [M+K]+ m/z 848.5566

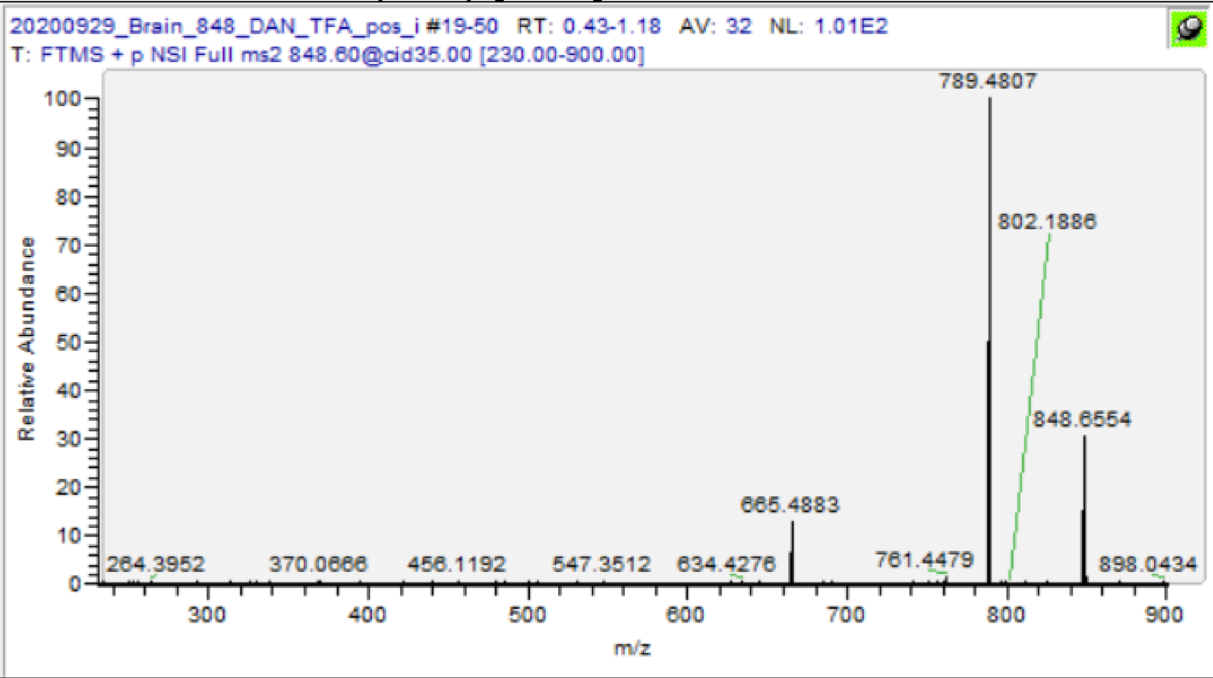

| m/z | [I] | Species | Formula | Da | ppm |
|---|---|---|---|---|---|
| 665.4885 | 12.82 | PL(38:4) -Hg +K | C41H70O4K | 665.4906 | 3.11 |
| 789.4807 | 100 | PC(38:4) -TMA +K | C43H75O8PK | 789.4831 | 3.05 |
| **848.5556ɣ** | **15.12** | **PC(38:4) +K** | **C46H84O8NPK** | **848.5566** | **1.19** |

ɣ Value from MS1 scan

HexCer(d18:1/24:1(2OH)) [M+Na]+ m/z 848.6586

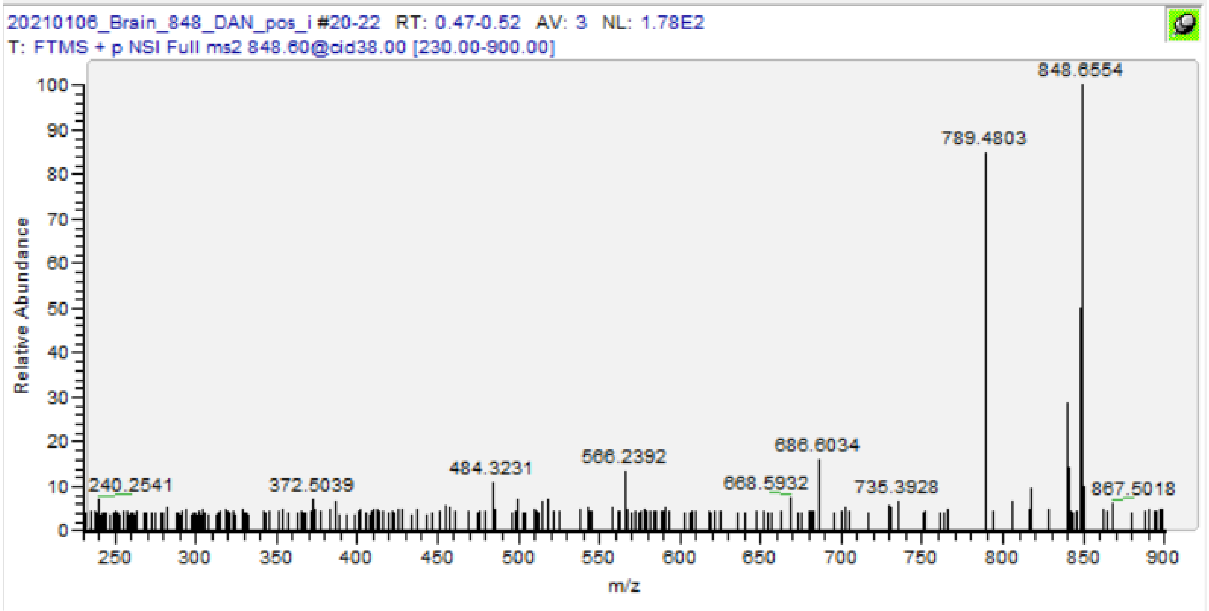

| m/z | [I] | Species | Formula | Da | ppm |
|---|---|---|---|---|---|
| 484.3231 | 3.82 | HexCer(d18:1/24:1(2OH)) -C24 + Na | C24H47O7NNa | 484.3245 | 2.83 |
| 668.5932 | 1.79 | HexCer(d18:1/24:1(2OH)) -Glc -H2O + Na | C42H79O3NNa | 668.5952 | 3.02 |
| 686.6034 | 4.79 | HexCer(d18:1/24:1(2OH)) -Glc + Na | C42H81O4NNa | 686.6058 | 3.47 |
| **848.6564** | **100** | **HexCer(d18:1/24:1(2OH)) + Na** | **C48H91O9NNa** | **848.6586** | **2.59** |

PC(40:6) [M+K]+ m/z 872.5566

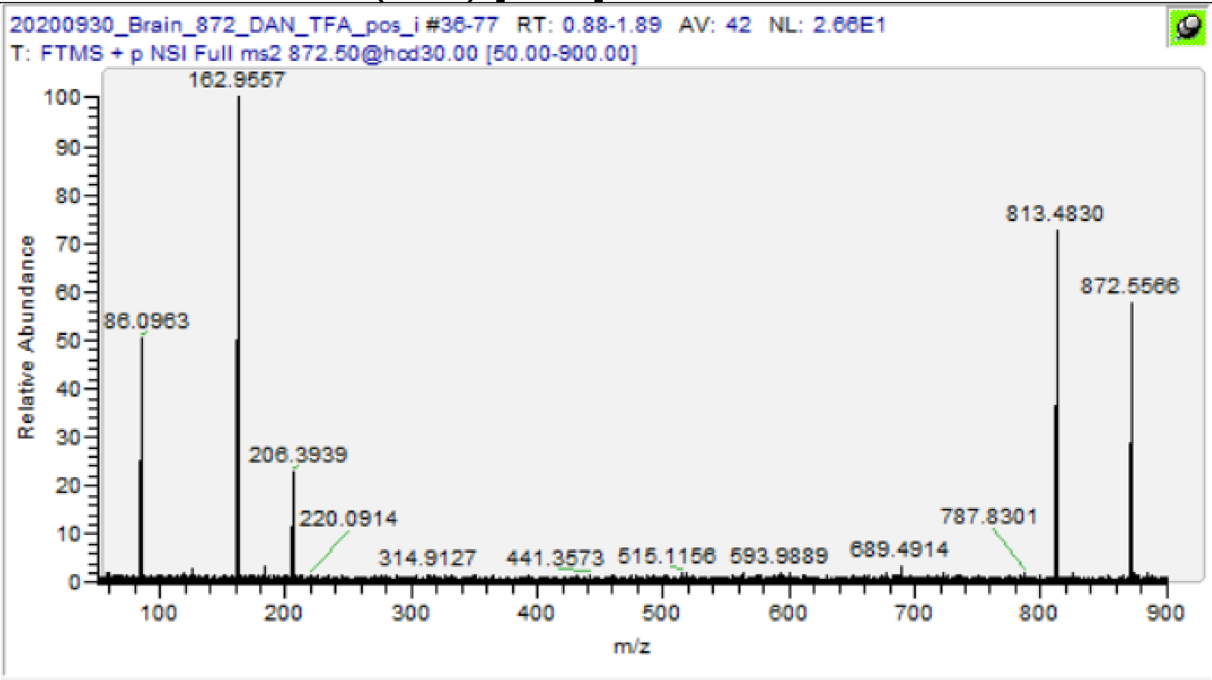

| m/z | [I] | Species | Formula | Da | ppm |
|---|---|---|---|---|---|
| 86.0963 | 49.49 | PC-Frag | C5H12N | 86.09643 | 1.51 |
| 162.9557 | 100 | PC-Frag +K | C2H5O4PK | 162.9557 | 0.00 |
| 184.0735 | 4 | PC-Hg | C5H15O4NP | 184.0733 | -0.98 |
| 689.4914 | 4.1 | PL(40:6) -Hg +K | C43H70O4K | 689.4906 | -1.20 |
| 813.4829 | 97.88 | PC(40:6) -TMA +K | C45H75O8PK | 813.4831 | 0.26 |
| **872.5565** | **77.68** | **PC(40:6) +K** | **C48H85O8NPK** | **872.5566** | **0.13** |

PC(16:0/18:1)+PC(16:0/16:0) [M+M+H]+ m/z 1494.147

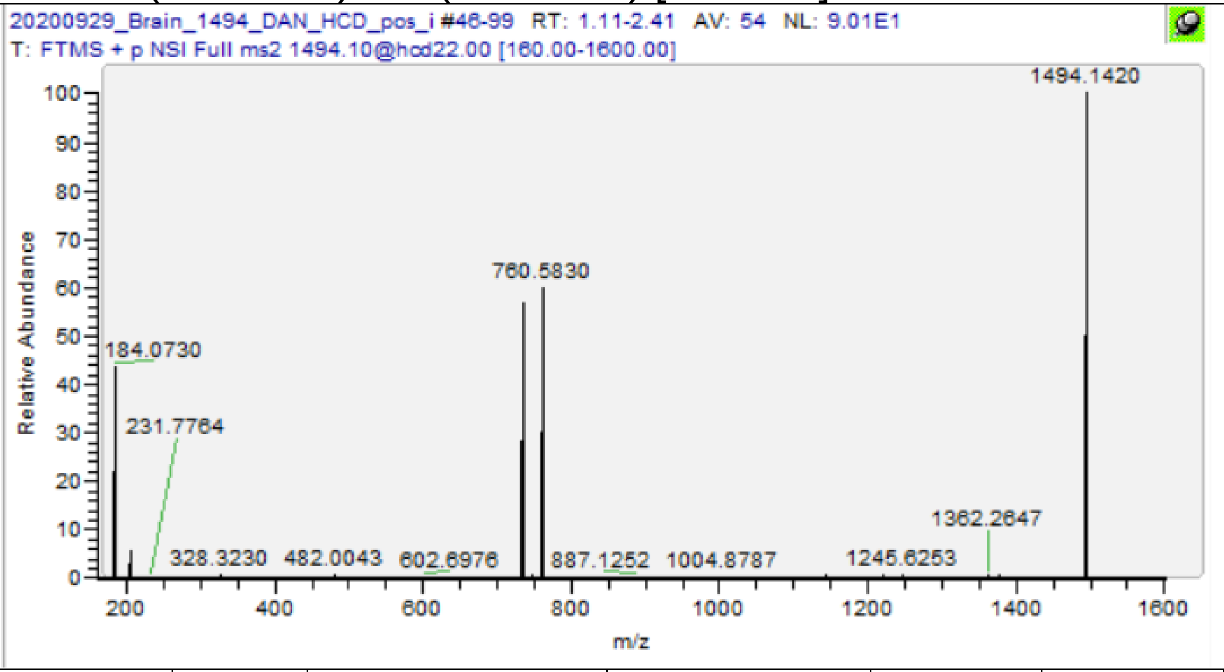

| m/z | [I] | Species | Formula | Da | ppm |
|---|---|---|---|---|---|
| 184.0736 | 27.75 | PC-Hg | C5H15O4NP | 184.0733 | -1.52 |
| 734.5698 | 96.83 | PC(16:0/16:0) | C40H81O8NP | 734.5694 | -0.50 |
| 760.5859 | 100 | PC(16:0/18:1) | C42H83O8NP | 760.5851 | -1.08 |
| **1494.1488** | **27.75** | **PC(16:0/18:1) +PC(16:0/16:0)** | **C82H163O16N2P2** | **1494.147** | **-1.04** |

PC(16:0/18:1) [2M+H]+ m/z 1520.163

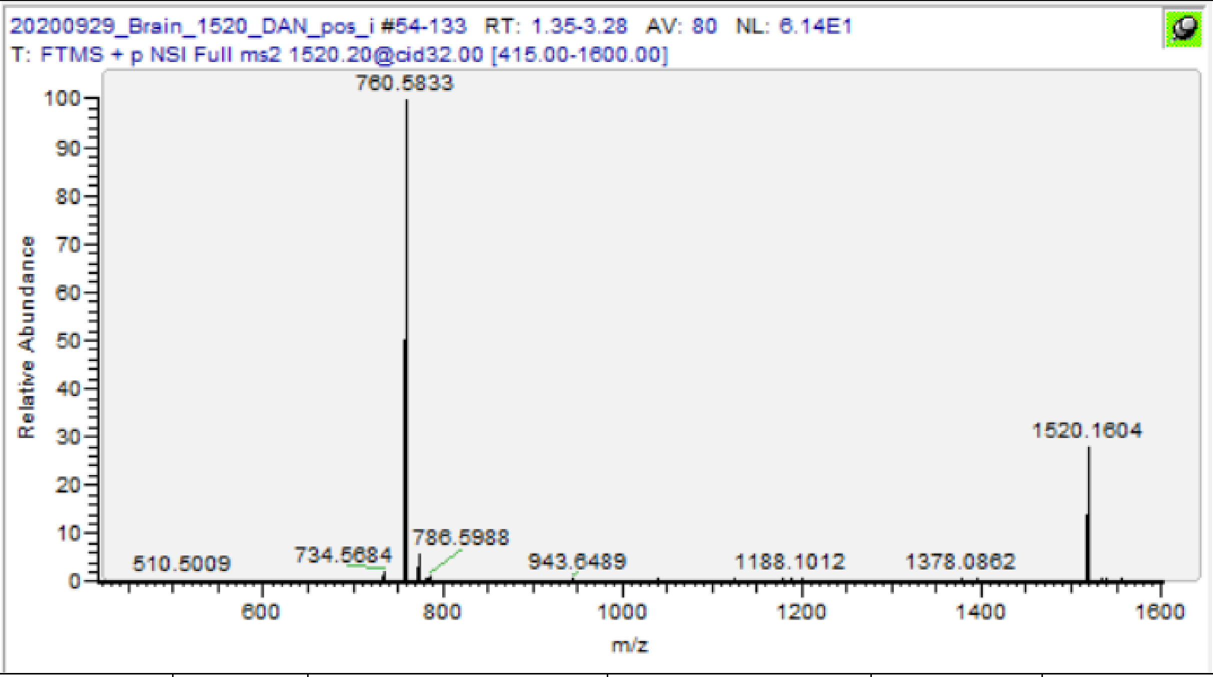

| m/z | [I] | Species | Formula | Da | ppm |
|---|---|---|---|---|---|
| 86.0964 | 1.86 | PC-Frag | C5H12N | 86.09643 | 0.35 |
| 124.9999 | 4.2 | PC-Frag | C2H6O4P | 124.9998 | -0.64 |
| 184.0735 | 100 | PC-Hg | C5H15O4NP | 184.0733 | -0.98 |
| 760.5833 | 100 | PC(16:0/18:1) | C42H83O8NP | 760.5851 | 2.34 |
| **1520.1604** | **26.74** | **2 xPC(16:0/18:1)** | **C84H165O16N2P2** | **1520.163** | **1.64** |
| 734.5683 | 2.3 | PC(16:0/16:0) | C40H81O8NP | 734.5694 | 1.54 |
| 786.5991 | 1.14 | PC(18:1/18:1) | C44H85O8NP | 786.6007 | 2.07 |

PC(18:1/18:0)+PC(16:0/18:1) [M+M+H]+ m/z 1548.194

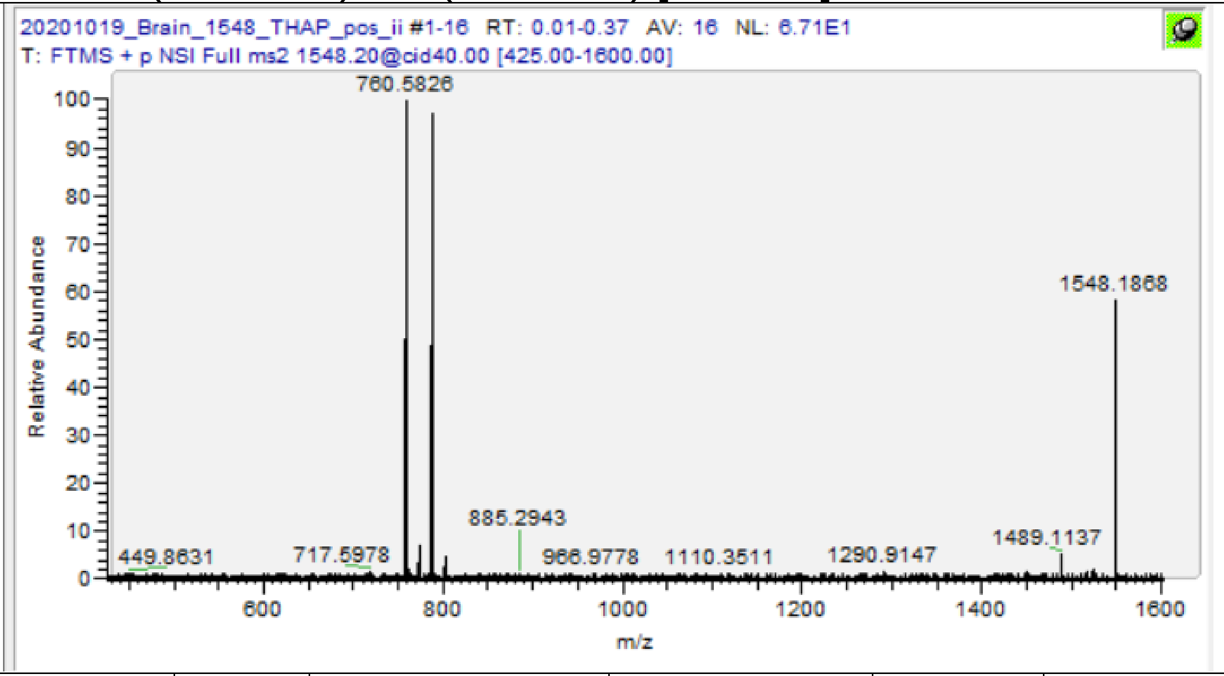

| m/z | [I] | Species | Formula | Da | ppm |
|---|---|---|---|---|---|
| 760.5826 | 100 | PC(16:0/18:1) | C42H83O8NP | 760.5851 | 3.26 |
| 788.6141 | 97.64 | PC(18:1/18:0) | C44H87O8NP | 788.6164 | 2.89 |
| **1548.1868** | **54.73** | **PC(18:1/18:0) +PC(16:0/18:1)** | **C86H169O16N2P2** | **1548.194** | **4.64** |

### Negative ion mode MS/MS, Identified species

Cer(d18:0/18:1) [M-H]- m/z 564.535

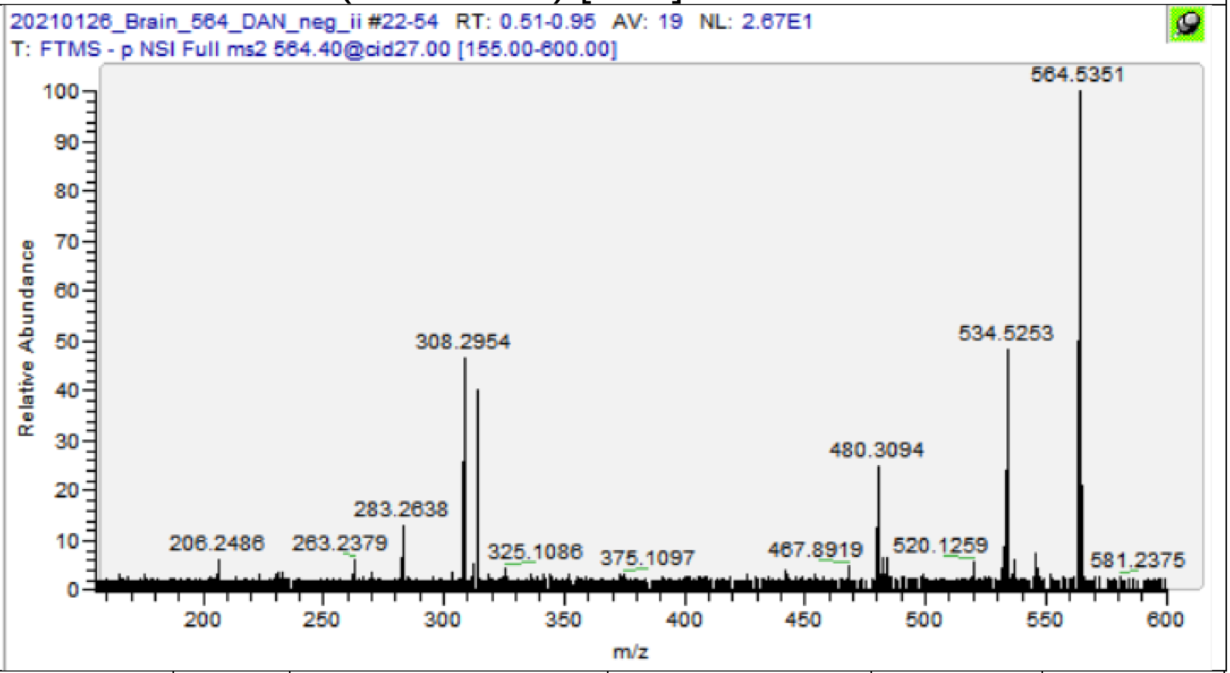

| m/z | [I] | Species | Formula | Da | ppm |
|---|---|---|---|---|---|
| 263.2378 | 2.17 | Cer(18:1)-Frag | C18H31O | 263.2369 | -3.27 |
| 283.2638 | 3.37 | FA 18:0 | C18H35O2 | 283.2643 | 1.59 |
| 308.2954 | 14.08 | Cer(18:1)-Frag | C20H38ON | 308.2948 | -1.98 |
| 534.5253 | 12.7 | Cer(d18:0/18:1) -COH2 | C35H68O2N | 534.5245 | -1.57 |
| **564.535** | **100** | **Cer(d18:0/18:1)** | **C36H70O3N** | **564.535** | **0.04** |

CerP(d18:1/18:0) [M-H]- m/z 644.5025

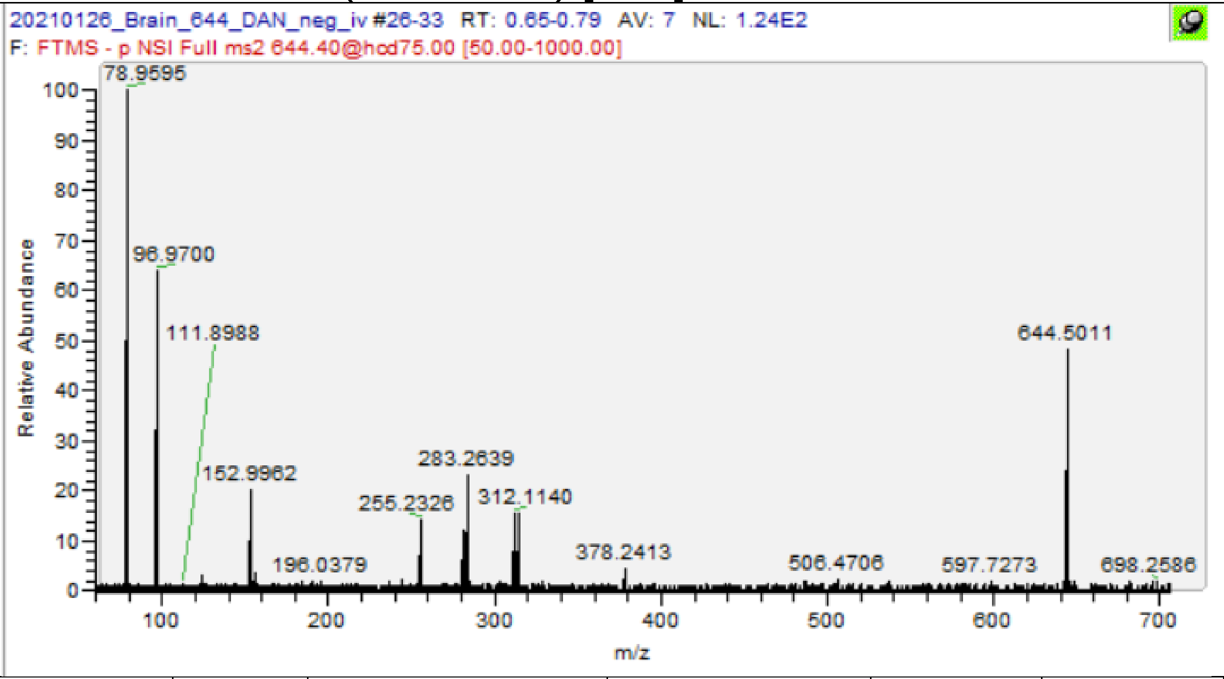

| m/z | [I] | Species | Formula | Da | ppm |
|---|---|---|---|---|---|
| 78.9595 | 100 | PL-Frag | PO3 | 78.95905 | -5.70 |
| 96.97 | 64.59 | PL-Frag | H2O4P | 96.96962 | -3.92 |
| 152.9962 | 20.18 | PL-Frag | C3H6O5P | 152.9958 | -2.42 |
| 378.2412 | 12.94 | CerP(d18:1) | C18H37O5NP | 378.2415 | 0.74 |
| **644.5019** | **100** | **CerP(d18:1/18:0)** | **C36H71NO6P** | **644.5025** | **0.85** |

## PA(16:0/16:1) [M-H]- m/z 645.4501

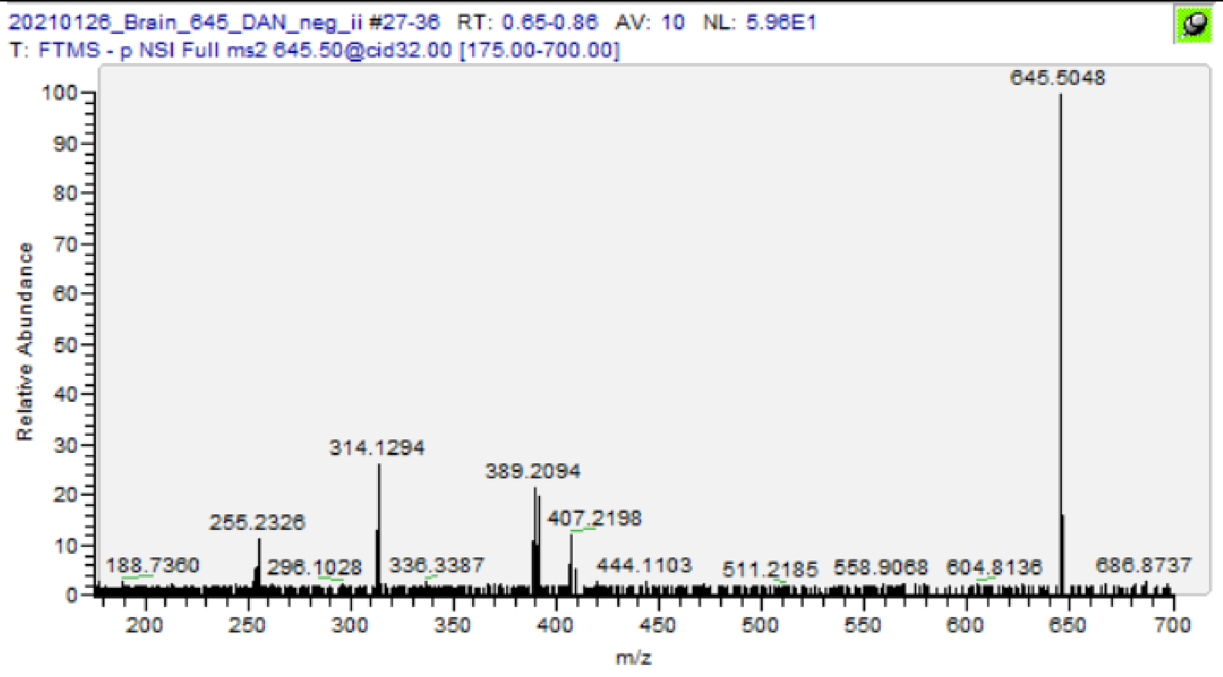

| m/z | [I] | Species | Formula | Da | ppm |
|---|---|---|---|---|---|
| 253.2169 | 4.2 | FA 16:1 | C16H29O2 | 253.2173 | 1.58 |
| 255.2326 | 13.04 | FA 16:0 | C16H31O2 | 255.233 | 1.37 |
| 389.2094 | 17.37 | PA(16:1) -H2O | C19H34O6P | 389.2099 | 1.16 |
| 391.225 | 16 | PA(16:0) -H2O | C19H36O6P | 391.2255 | 1.28 |
| 407.2198 | 12.5 | PA(16:1) | C19H36O7P | 407.2204 | 1.50 |
| 409.2355 | 7.71 | PA(16:0) | C19H38O7P | 409.2361 | 1.37 |
| **645.4503** | **3.83** | **PA(16:0/16:1)** | **C35H66O8P** | **645.4501** | **-0.34** |

PA(16:0/16:0) [M-H]- m/z 647.4657

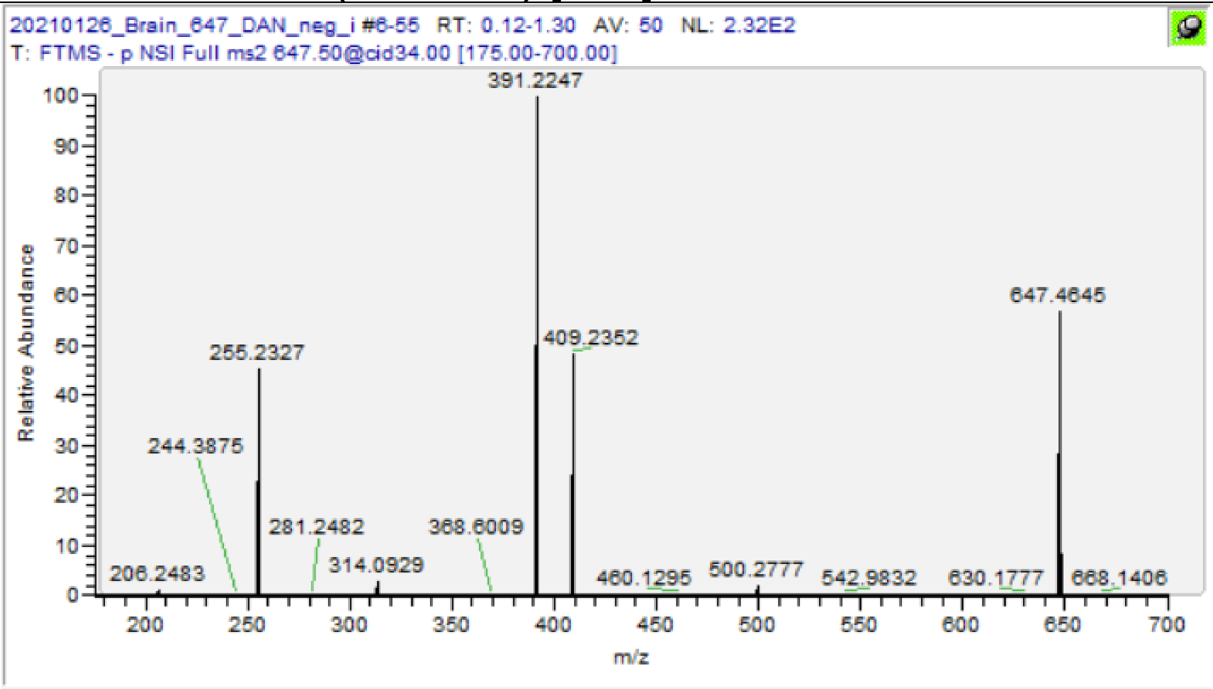

| m/z | [I] | Species | Formula | Da | ppm |
|---|---|---|---|---|---|
| 255.2327 | 53.81 | FA 16:0 | C16H31O2 | 255.233 | 0.98 |
| 391.2247 | 100 | PA(16:0) -H2O | C19H36O6P | 391.2255 | 2.04 |
| 409.2352 | 53.7 | PA(16:0) | C19H38O7P | 409.2361 | 2.10 |
| **647.4645** | **55.4** | **PA(16:0/16:0)** | **C35H68O8P** | **647.4657** | **1.90** |

### CerP(d38:2) [M-H]- m/z 670.5181

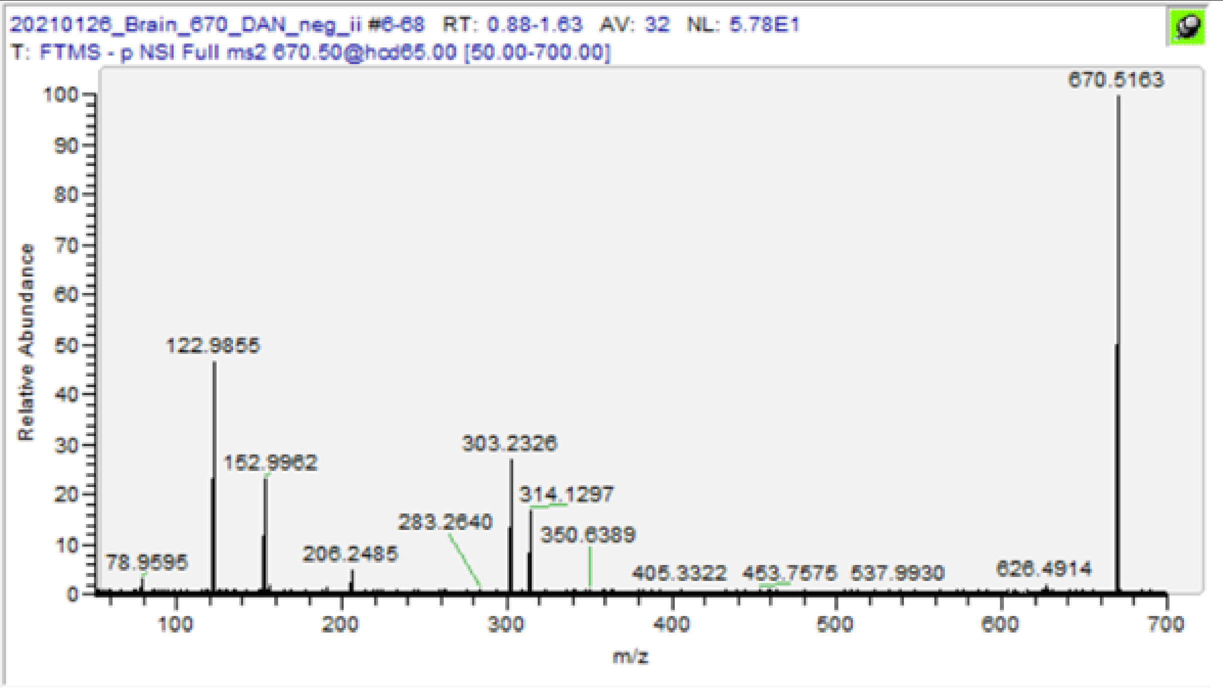

| m/z | [I] | Species | Formula | Da | ppm |
|---|---|---|---|---|---|
| 78.9595 | 5.49 | PL-Frag | PO3 | 78.95905 | -5.70 |
| 122.9855 | 63.42 | PL-Frag | C2H4O4P | 122.9853 | -1.87 |
| 152.9962 | 30.12 | PL-Frag | C3H6O5P | 152.9958 | -2.42 |
| 626.4922 | 2.18 | CerP 36:1;O2 -H2O | C36H69O5NP | 626.4919 | -0.51 |
| **670.5173** | **34.76** | **CerP(d38:2)** | **C38H73O6NP** | **670.5181** | **1.19** |

PA(16:0/18:1) [M-H]- m/z 673.4814

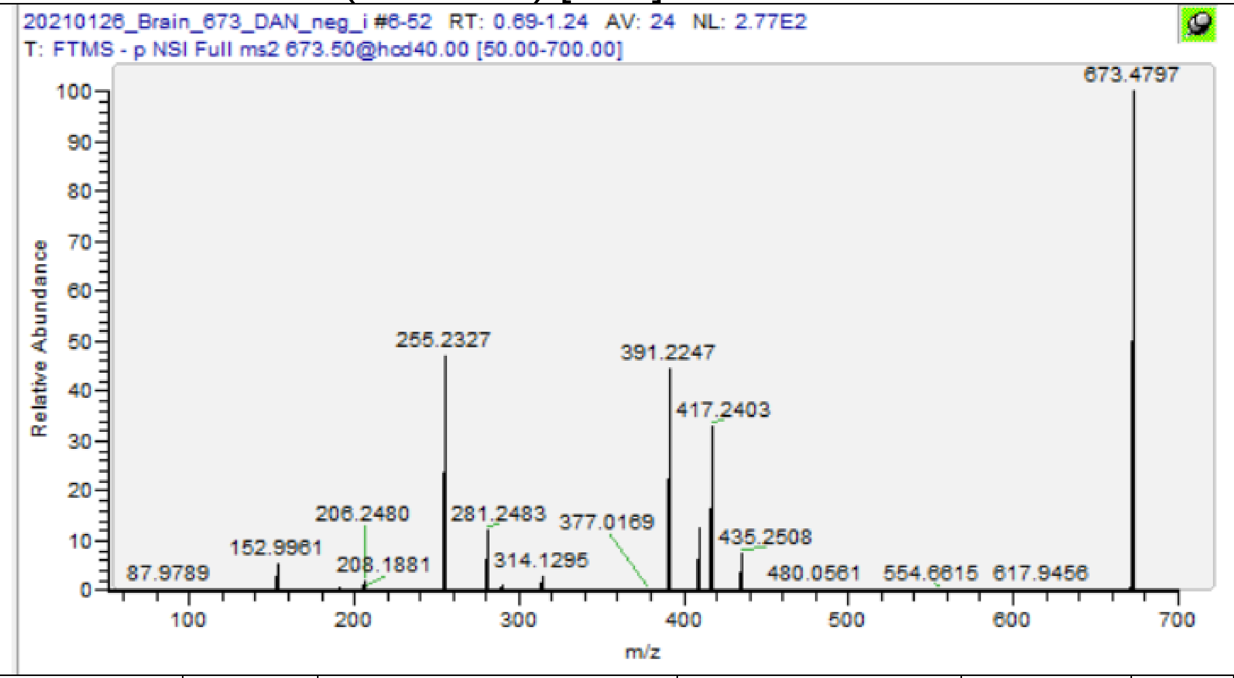

| m/z | [I] | Species | Formula | Da | ppm |
|---|---|---|---|---|---|
| 152.9962 | 9.48 | PL-Frag | C3H6O5P | 152.9958 | -2.42 |
| 255.2327 | 79.48 | FA 16:0 | C16H31O2 | 255.233 | 0.98 |
| 281.2483 | 23.66 | FA 18:1 | C18H33O2 | 281.2486 | 1.07 |
| 391.2246 | 62.74 | PA(16:0) -H2O | C19H36O6P | 391.2255 | 2.30 |
| 409.2352 | 19.64 | PA(16:0) | C19H38O7P | 409.2361 | 2.10 |
| 417.2402 | 48.62 | PA(18:1) -H2O | C21H38O6P | 417.2412 | 2.28 |
| 435.2506 | 12.22 | PA(18:1) | C21H40O7P | 435.2517 | 2.55 |
| **673.4795** | **100** | **PA(16:0/18:1)** | **C37H70O8P** | **673.4814** | **2.79** |

SM(d34:1) [M-CH3]- m/z 687.5447

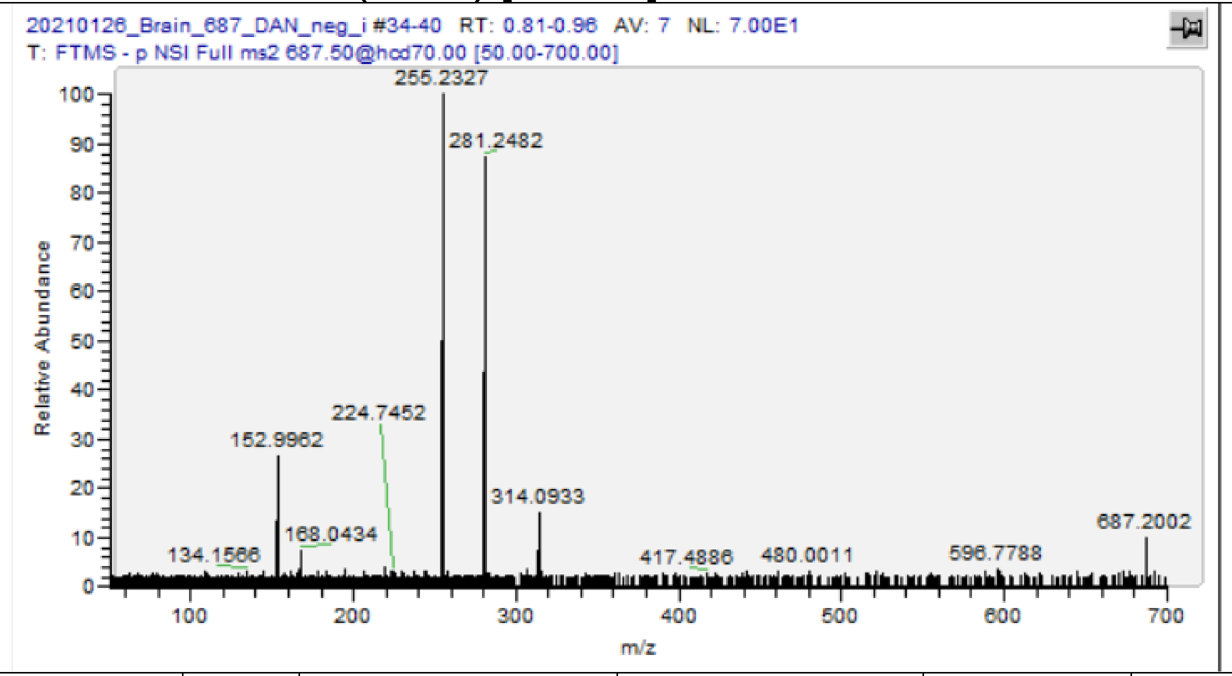

| m/z | [I] | Species | Formula | Da | ppm |
|---|---|---|---|---|---|
| 168.0434 | 7.87 | PC-Hg | C4H11O4NP | 168.0431 | -1.67 |
| 255.2327 | 86.39 | FA 16:0 | C16H31O2 | 255.233 | 0.98 |
| 281.2483 | 100 | FA 18:1 | C18H33O2 | 281.2486 | 1.07 |
| **687.5426** | **18.33** | **SM(d18:1/16:0) -CH3 / SM(d16:0/18:1) -CH3** | **C38H76O6N2P** | **687.5447** | **2.98** |

## SM(d18:1/18:0) [M-CH3]- m/z 715.576

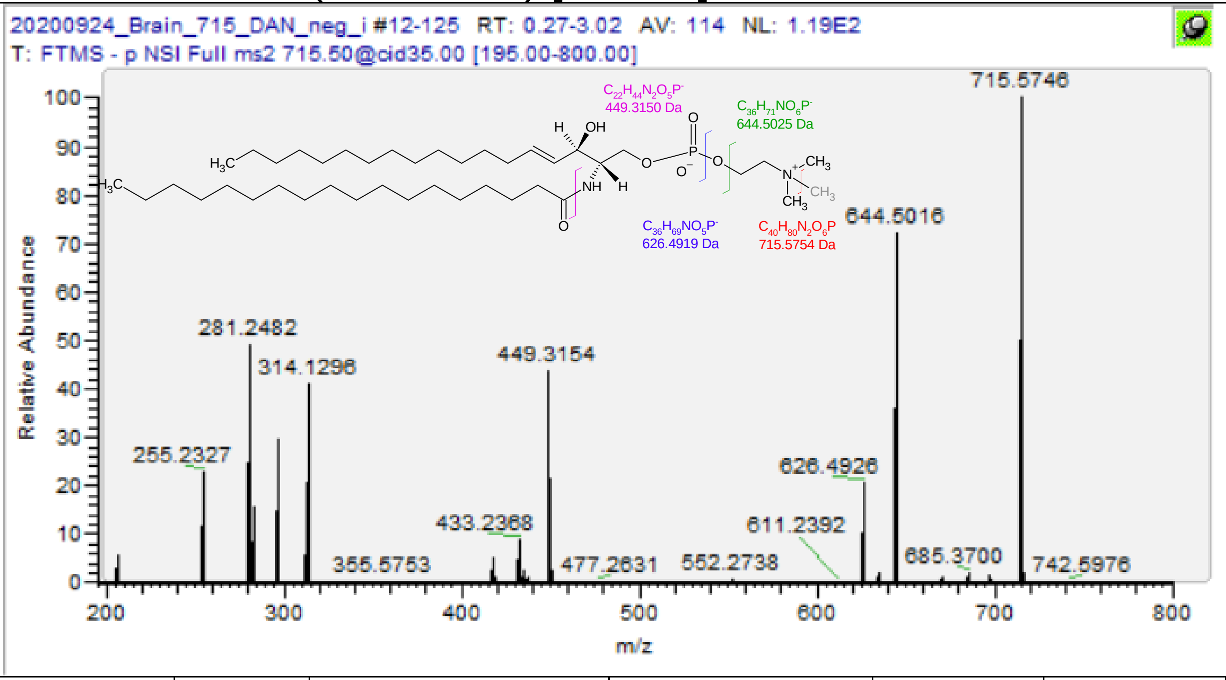

| m/z | [I] | Species | Formula | Da | ppm |
|---|---|---|---|---|---|
| 449.3155 | 60.64 | LSM(18:1);O2 -CH3 | C22H46N2O5P | 449.315 | -1.16 |
| 626.4927 | 28.36 | CerP 36:1;O2 -H2O | C36H69O5NP | 626.4919 | -1.31 |
| 644.5017 | 100 | CerP 36:1;O2 | C36H71NO6P | 644.5025 | 1.16 |
| **715.5750** | **55.63** | **SM(d18:1/18:0) -CH3** | **C40H80O6N2P** | **715.576** | **1.33** |
| 255.2327 | 33.11 | FA(16:0) | C16H31O2 | 255.233 | 0.98 |
| 281.2482 | 68.71 | FA(18:1) | C18H33O2 | 281.2486 | 1.42 |
| 283.2642 | 21.79 | FA(18:0) | C18H35O2 | 283.2643 | 0.18 |
| 297.2433 | 40.61 | FA(18:1);O | C18H33O3 | 297.2435 | 0.74 |
| 417.2417 | 6.56 | PA(18:1) -H2O | C21H38O6P | 417.2412 | -1.32 |
| 433.2368 | 13.01 | PA(18:2) | C21H38O7P | 433.2361 | -1.71 |
| 435.2523 | 3 | PA(18:1) | C21H40O7P | 435.2517 | -1.36 |
| 437.2674 | 1.75 | PA(18:0) | C21H42O7P | 437.2674 | -0.09 |

PE(16:0/18:0) [M-H]- m/z 718.5392

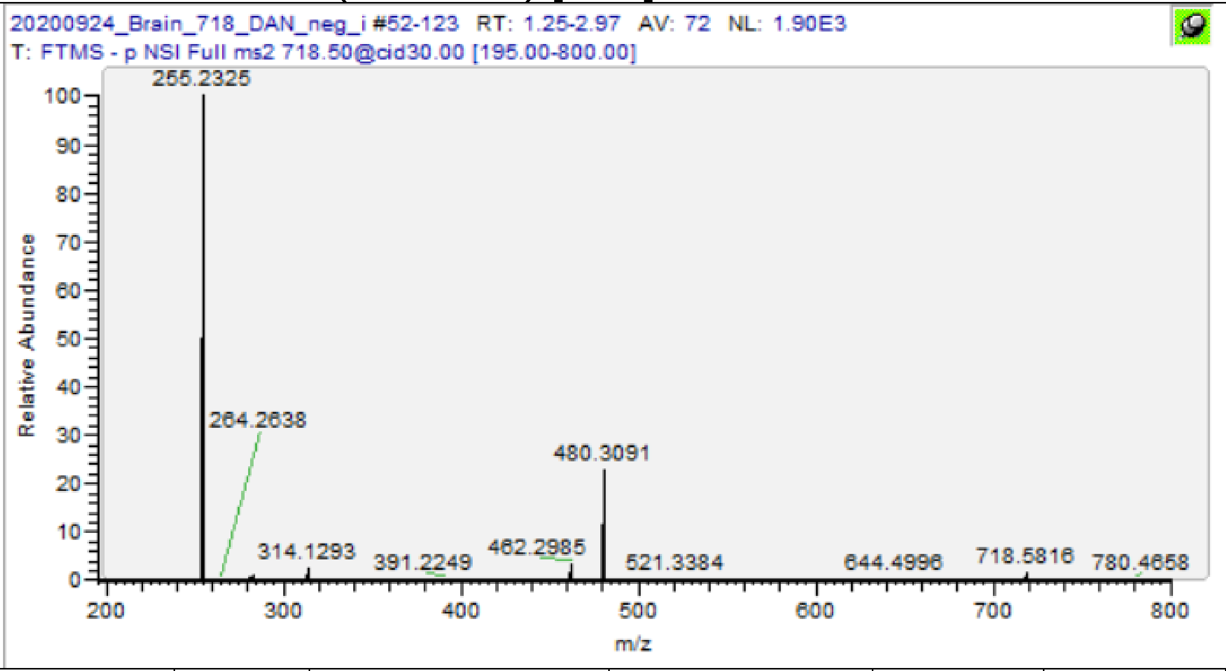

| m/z | [I] | Species | Formula | Da | ppm |
|---|---|---|---|---|---|
| 255.2325 | 100 | FA(16:0) | C16H31O2 | 255.233 | 1.76 |
| 283.2638 | 0.96 | FA(18:0) | C18H35O2 | 283.2643 | 1.59 |
| 419.2559 | 0.06 | PA(18:0) -H2O | C21H40O6P | 419.2568 | 2.15 |
| 437.2662 | 0.06 | PA(18:0/-) | C21H42O7P | 437.2674 | 2.65 |
| 462.2985 | 3.6 | PE(18:0) -H2O | C23H45O6NP | 462.299 | 1.08 |
| 480.3091 | 23.25 | PE(18:0) | C23H47O7NP | 480.3096 | 0.96 |
| **718.5377ɣ** | **1.84** | **PE(16:0/18:0)** | **C39H77O8NP** | **718.5392** | **2.13** |

ɣ Value from MS1 scan

PE(P-18:1/18:1) [M-H]- m/z 726.5443

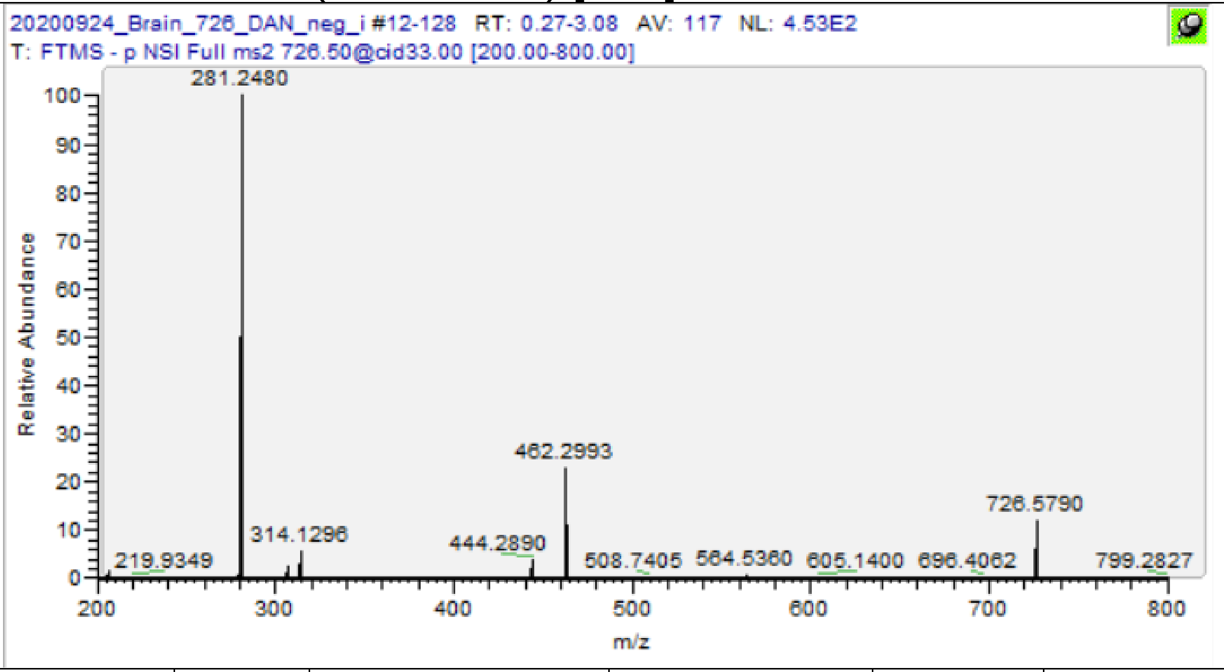

| m/z | [I] | Species | Formula | Da | ppm |
|---|---|---|---|---|---|
| 281.2480 | 100 | FA(18:1) | C18H33O2 | 281.2486 | 2.13 |
| 444.2890 | 3.88 | PE(P-18:1) -H2O | C23H43O5NP | 444.2884 | -1.28 |
| 462.2992 | 22.03 | PE(P-18:1) | C23H45O6NP | 462.299 | -0.43 |
| **726.5427** | **12.21** | **PE(P-18:1/18:1)** | **C41H77O7NP** | **726.5443** | **2.22** |
| 564.5361 | 0.82 | Cer 36:1;O2 | C36H70NO3 | 564.5361 | 0.00 |
| 726.5789 | 13.17 | HexCer 36:1;O2 | C42H80NO8 | 726.5889 | 13.76 |

PE(P-18:0/18:1) [M-H]- m/z 728.56
PE(P-16:0/20:1) [M-H]- m/z 728.56

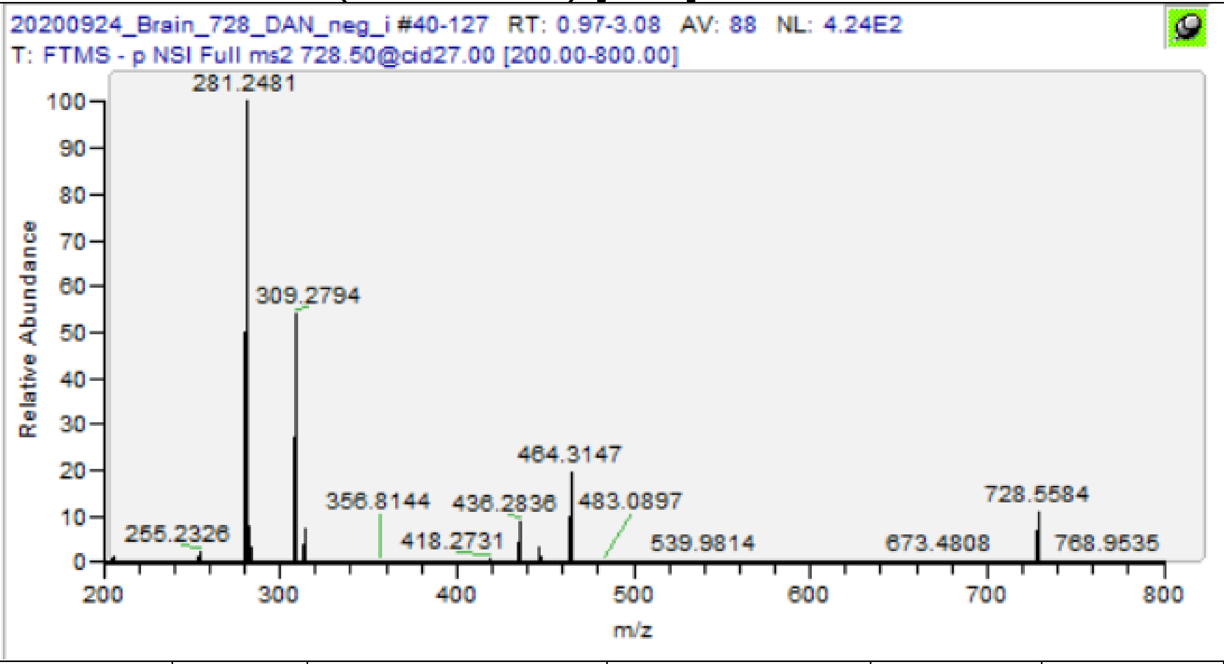

| m/z | [I] | Species | Formula | Da | ppm |
|---|---|---|---|---|---|
| 281.2481 | 100 | FA(18:1) | C18H33O2 | 281.2486 | 1.78 |
| 309.2794 | 55.94 | FA 20:1 | C20H37O2 | 309.2799 | 1.62 |
| 418.2731 | 0.76 | PE(P-16:0) -H2O | C21H41O5NP | 418.2728 | -0.77 |
| 436.2837 | 9.07 | PE(P-16:0) | C21H43O6NP | 436.2834 | -0.80 |
| 446.3046 | 3.18 | PE(P-18:0) -H2O | C23H45O5NP | 446.3041 | -1.17 |
| 464.3147 | 20.12 | PE(P-18:0) | C23H47O6NP | 464.3147 | -0.11 |
| **728.5584** | **9.35** | **PE(P-18:0/18:1)/ PE(P-16:0/20:1)** | **C41H79O7NP** | **728.56** | **2.14** |

PE(18:1/18:1) [M-H]- m/z 742.5392

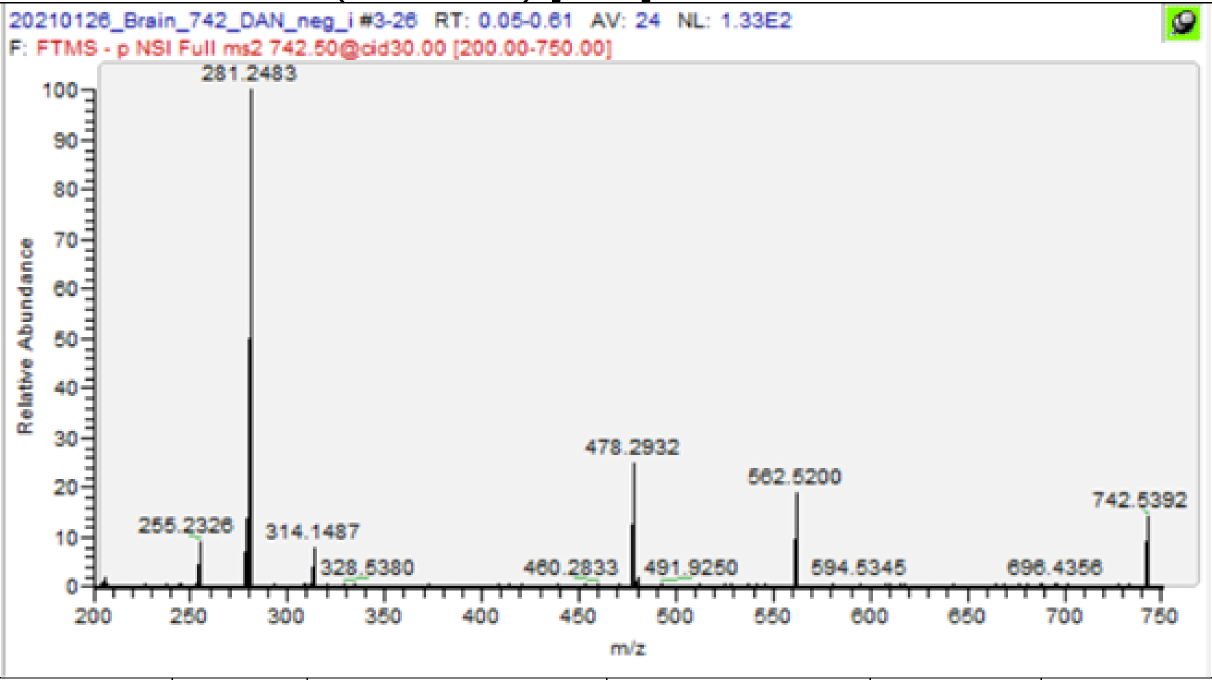

| m/z | [I] | Species | Formula | Da | ppm |
|---|---|---|---|---|---|
| 281.2483 | 100 | FA 18:1 | C18H33O2 | 281.2486 | 1.06 |
| 460.2833 | 0.89 | PE(18:1) -OH | C23H43O6NP | 460.2834 | 0.10 |
| 478.2933 | 25.8 | PC(18:1) | C23H45O7NP | 478.2939 | 1.27 |
| **742.5391** | **13.64** | **PE(18:1/18:1)** | **C41H77O8NP** | **742.5392** | **0.17** |

PC(16:0/18:1) [M-CH3]- m/z 744.5549

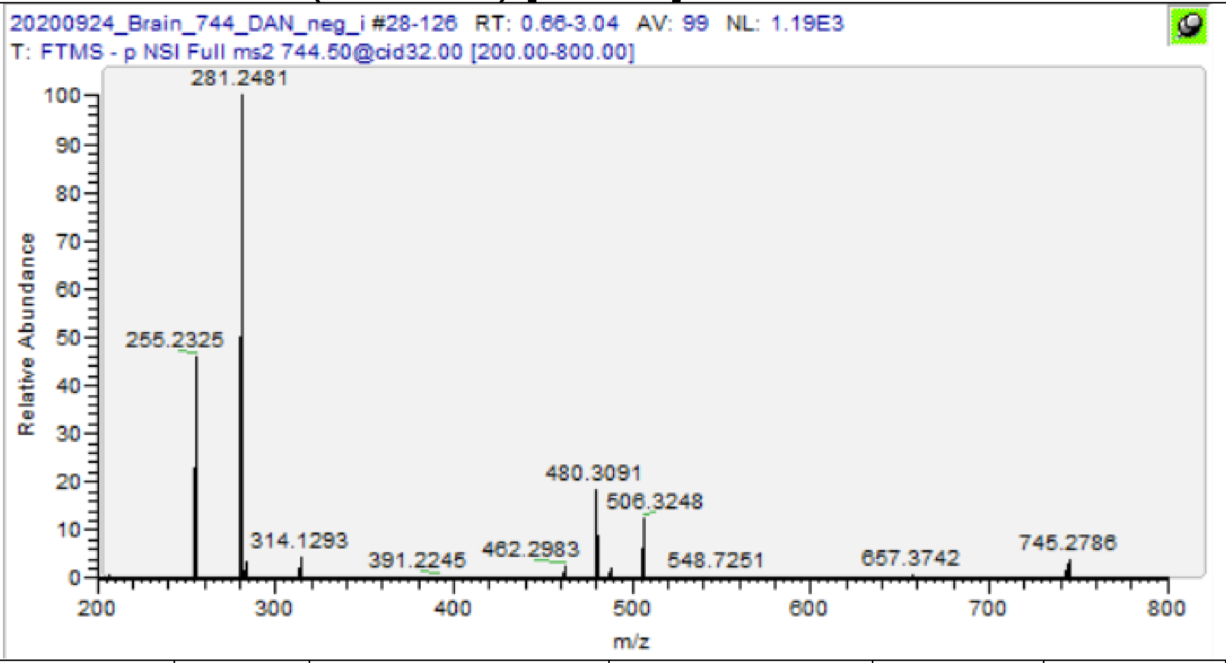

| m/z | [I] | Species | Formula | Da | ppm |
|---|---|---|---|---|---|
| 255.2325 | 46.82 | FA(16:0) | C16H31O2 | 255.233 | 1.76 |
| 281.2481 | 100 | FA(18:1) | C18H33O2 | 281.2486 | 1.78 |
| 462.2983 | 2.58 | PC(16:0) -CH3 -OH | C23H45O6NP | 462.299 | 1.51 |
| 480.3091 | 18.72 | PC(16:0) -CH3 | C23H47O7NP | 480.3096 | 0.96 |
| 488.3138 | 2.05 | PC(18:1) -CH3 -H2O | C25H47O6NP | 488.3147 | 1.74 |
| 506.3247 | 12.55 | PC(18:1) -CH3 | C25H49O7NP | 506.3252 | 1.01 |
| **744.5535** | **12.55** | **PC(16:0/18:1)-CH3** | **C41H79O8NP** | **744.5549** | **1.85** |

PA(18:0/22:6) [M-H]- m/z 747.497

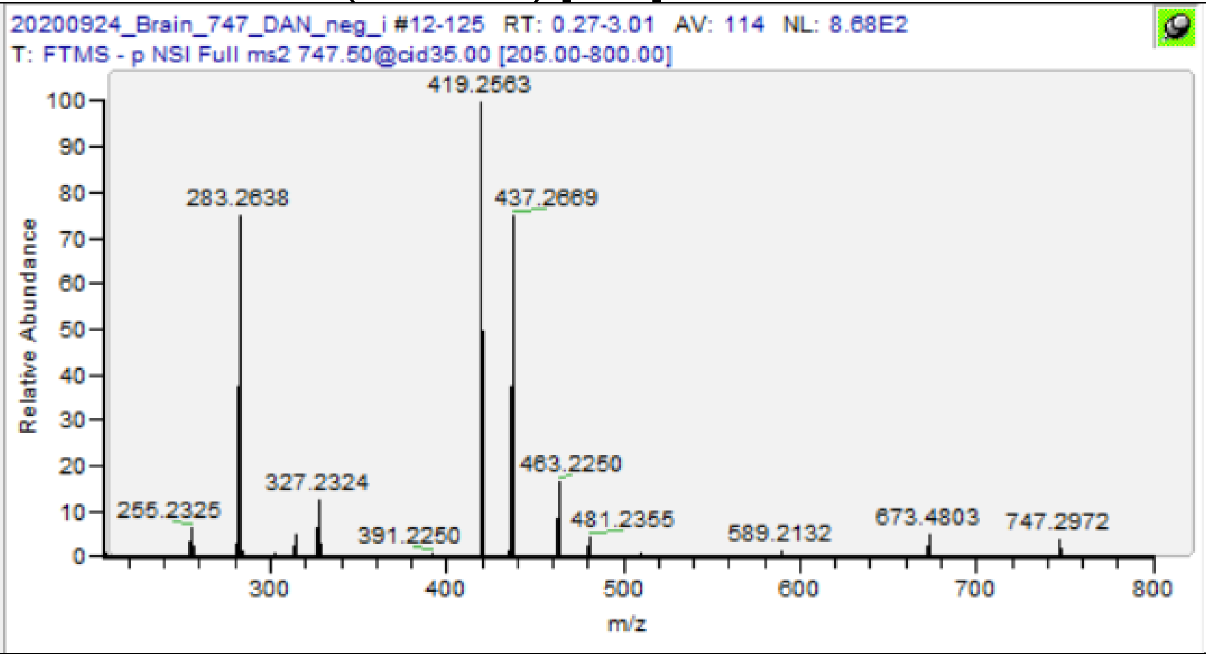

| m/z | [I] | Species | Formula | Da | ppm |
|---|---|---|---|---|---|
| 283.2427 | 4.9 | FA(22:6)-CO2 | C21H31 | 283.2431 | 1.48 |
| 283.2638 | 72.41 | FA(18:0) | C18H35O2 | 283.2643 | 1.59 |
| 327.2324 | 12.09 | FA 22:6 | C22H31O2 | 327.233 | 1.68 |
| 419.2563 | 100 | PA(18:0) -H2O | C21H40O6P | 419.2568 | 1.19 |
| 437.2668 | 77.11 | PA(18:0) | C21H42O7P | 437.2674 | 1.28 |
| 463.2250 | 16.25 | PA(22:6) -H2O | C25H36O6P | 463.2255 | 1.08 |
| 481.2355 | 4.1 | PA(22:6) | C25H38O7P | 481.2361 | 1.16 |
| **747.4959** | **30** | **PA(18:0/22:6)** | **C43H72O8P** | **747.497** | **1.51** |
| 255.2325 | 6.05 | FA(16:0) | C16H31O2 | 255.233 | 1.76 |
| 281.2481 | 6.03 | FA(18:1) | C18H33O2 | 281.2486 | 1.78 |
| 391.2250 | 0.88 | PA(16:0) -H2O | C19H36O6P | 391.2255 | 1.28 |
| 465.2620 | 0.15 | PG(16:0)-H2O | C22H42O8P | 465.2623 | 0.60 |
| 483.2720 | 0.26 | PG(16:0) | C22H44O9P | 483.2728 | 1.74 |
| 673.4803 | 4.61 | PA(18:1/16:0) | C37H70O8P | 673.4814 | 1.60 |
| 747.5150 | 4.5 | PG(18:1/16:0) | C40H76O10P | 747.5182 | 4.30 |

PE(P-18:1/20:1) [M-H]- m/z 754.5756

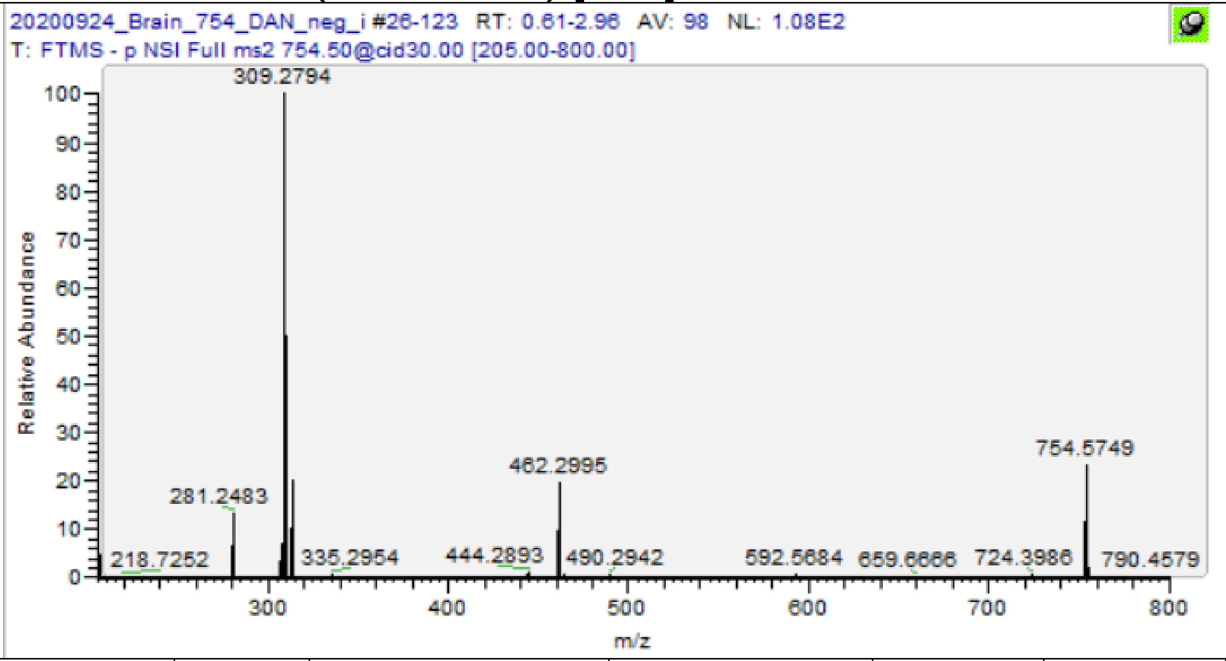

| m/z | [I] | Species | Formula | Da | ppm |
|---|---|---|---|---|---|
| 309.2794 | 100 | FA 20:1 | C20H37O2 | 309.2799 | 1.62 |
| 444.2892 | 0.6 | PE(P-18:1) -H2O | C23H43O5NP | 444.2884 | -1.73 |
| 462.2995 | 18.2 | PE(P-18:1) | C23H45O6NP | 462.299 | -1.08 |
| **754.5749** | **25.72** | **PE(P-18:1/20:1)** | **C43H81O7NP** | **754.5756** | **0.94** |

## PE(18:0/20:4) [M-H]- m/z 766.5392

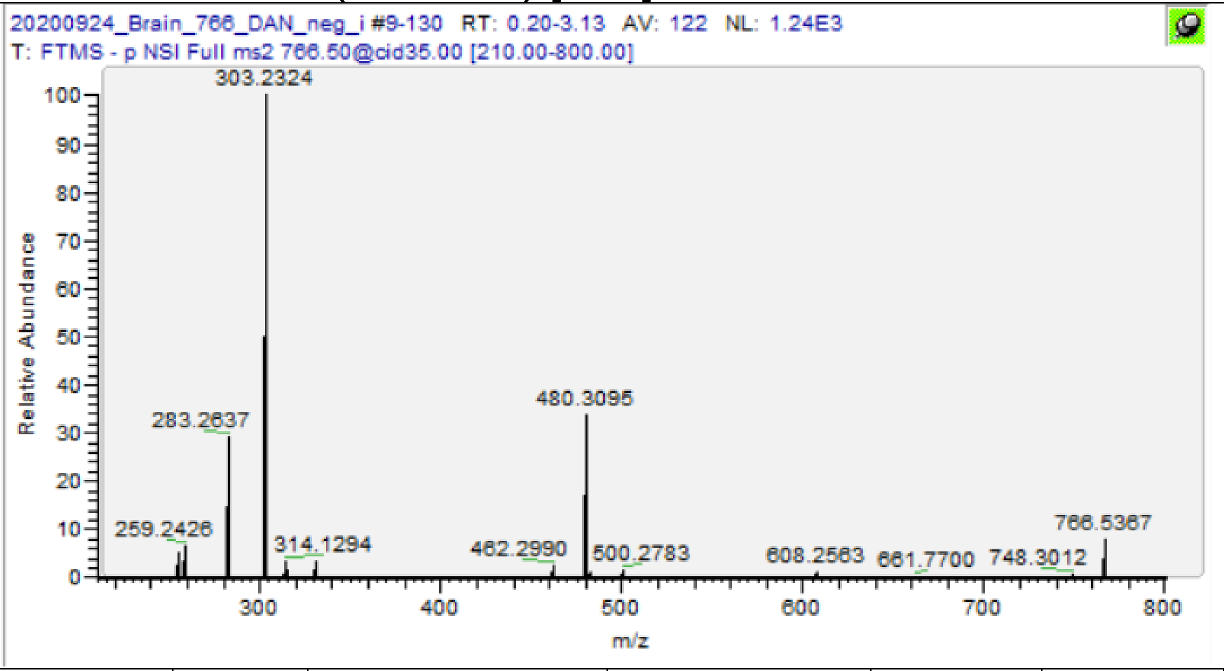

| m/z | [I] | Species | Formula | Da | ppm |
|---|---|---|---|---|---|
| 259.2427 | 6.18 | FA(20:4)-CO2 | C19H31 | 259.2431 | 1.62 |
| 283.2637 | 29.2 | FA(18:0) | C18H35O2 | 283.2643 | 1.94 |
| 303.2325 | 100 | FA(20:4) | C20H31O2 | 303.233 | 1.48 |
| 462.2990 | 2.58 | PE(18:0)-H2O | C23H45O6NP | 462.299 | 0.00 |
| 480.3095 | 33.25 | PE(18:0) | C23H47O7NP | 480.3096 | 0.12 |
| 482.2676 | 1.09 | PE(20:4)-H2O | C25H41O6NP | 482.2677 | 0.21 |
| 500.2784 | 1.71 | PE(20:4) | C25H43O7NP | 500.2783 | -0.28 |
| **766.5373** | **10.69** | **PE(18:0/20:4)** | **C43H77O8NP** | **766.5392** | **2.52** |
| 255.2325 | 4.9 | FA(16:0) | C16H31O2 | 255.233 | 1.76 |
| 331.2638 | 3.38 | FA 22:4 | C22H35O2 | 331.2643 | 1.36 |
| 452.2781 | 0.39 | PE(16:0) | C21H43O7NP | 452.2783 | 0.35 |
| 528.3091 | 0.22 | PE 22:4 | C27H47O7NP | 528.3096 | 0.87 |
| 766.5373 | 10.69 | PE(16:0/22:4) | C43H77O8NP | 766.5392 | 2.52 |

SM(d18:1/22:0) [M-CH3]- m/z 771.6386

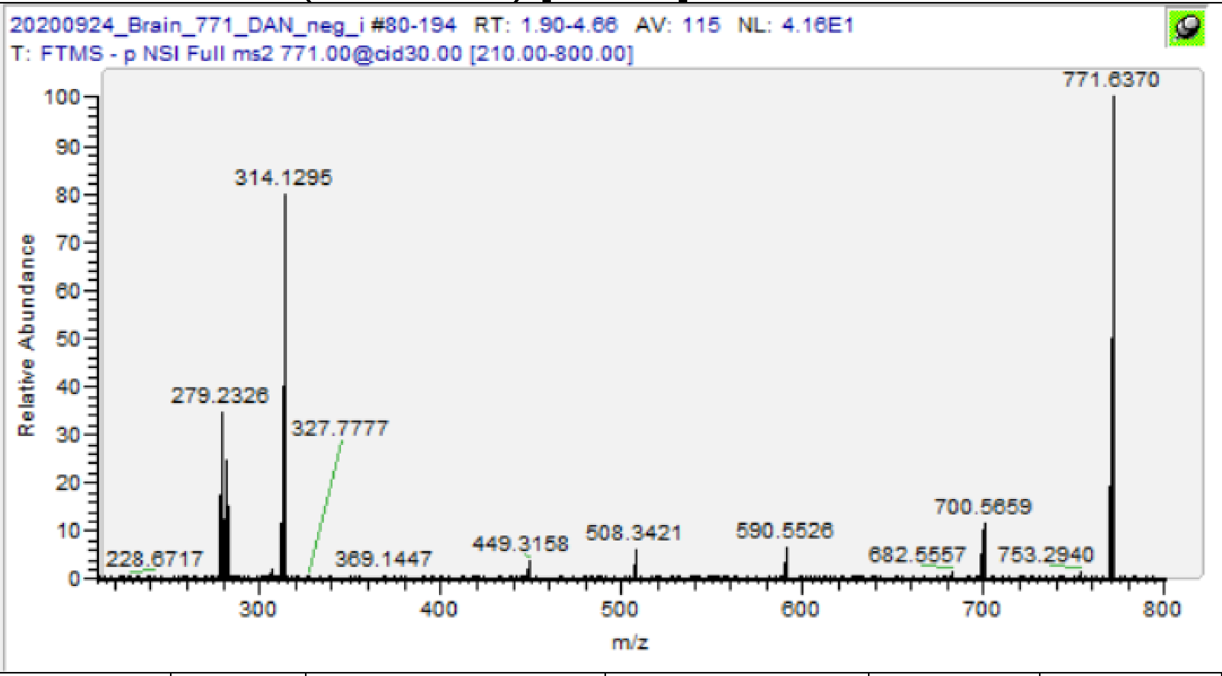

| m/z | [I] | Species | Formula | Da | ppm |
|---|---|---|---|---|---|
| 449.3158 | 3.53 | LSM 18:1;O2 -CH3 | C22H46N2O5P | 449.31498 | -1.83 |
| 682.5557 | 1.65 | CerP 40:1;O2 -H2O | C40H79O6NP | 682.55448 | -1.79 |
| 700.5659 | 11.84 | CerP 40:1;O2 | C40H79O6NP | 700.56505 | -1.21 |
| **771.637** | **100** | **SM(d18:1/22:0) -CH3** | **C44H88O6N2P** | **771.63855** | **2.01** |

PE(P-18:0/22:6) -H [M-H]- m/z 774.5443

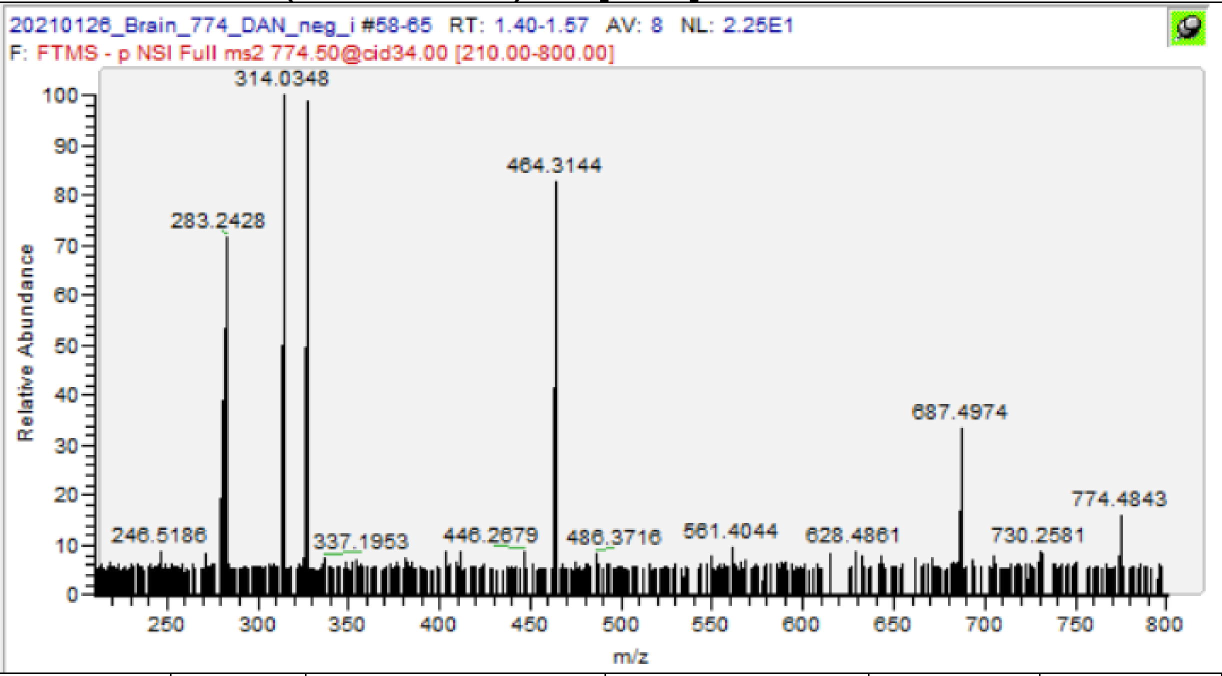

| m/z | [I] | Species | Formula | Da | ppm |
|---|---|---|---|---|---|
| 283.2426 | 48.86 | FA 22:6 -CO2 | C21H31 | 283.2431 | 1.84 |
| 327.2325 | 100 | FA 22:6 | C22H31O2 | 327.233 | 1.38 |
| 446.3045 | 11.79 | PE(P-18:0) -CH3 -H2O | C23H45O5NP | 446.3041 | -0.94 |
| 464.3148 | 85.87 | PE(P-18:0) | C23H47O6NP | 464.3147 | -0.32 |
| **774.5427** | **100** | **PE(P-18:0/22:6) -H** | **C45H77O7NP** | **774.5443** | **2.08** |

# PE(18:0/20:4(OH)) [M-H]- m/z 782.5341

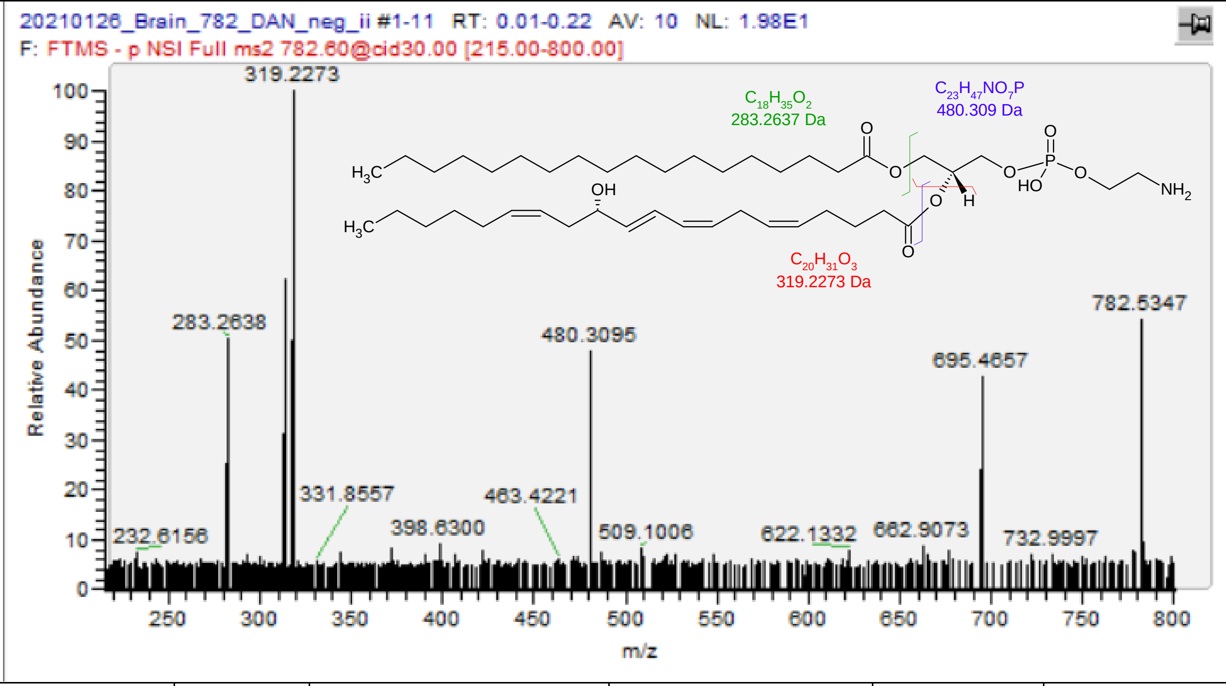

| m/z | [I] | Species | Formula | Da | ppm |
|---|---|---|---|---|---|
| 283.2638 | 49.38 | FA 18:0 | C18H35O2 | 283.2643 | 1.59 |
| 319.2273 | 100 | FA(20:4(OH) | C20H31O3 | 319.2279 | 1.79 |
| 480.3095 | 46.83 | PE(18:0) | C23H47O7NP | 480.3096 | 0.12 |
| **782.5347** | **53.53** | **PE(18:0/20:4(OH)** | **C43H77O9NP** | **782.5341** | **-0.72** |

PC(16:1/22:6) [M-CH3]- m/z 788.5236
PS(18:1/18:0) [M-H]- m/z 788.5447

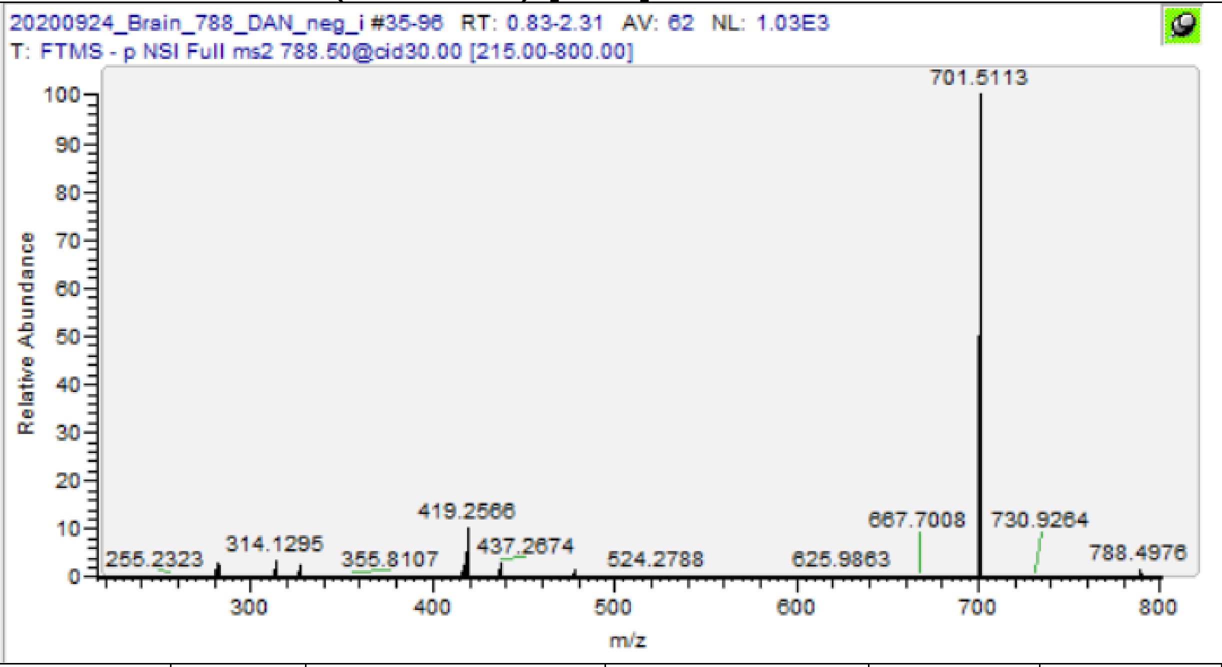

| m/z | [I] | Species | Formula | Da | ppm |
|---|---|---|---|---|---|
| 281.2482 | 2.96 | FA(18:1) | C18H33O2 | 281.2486 | 1.42 |
| 283.2638 | 2.78 | FA(18:0) | C18H35O2 | 283.2643 | 1.59 |
| 417.2409 | 2.74 | PA(18:1) -H2O | C21H38O6P | 417.2412 | 0.60 |
| 419.2565 | 10.72 | PA(18:0) -H2O | C21H40O6P | 419.2568 | 0.72 |
| 435.2516 | 0.15 | PA(18:1) | C21H40O7P | 435.2517 | 0.25 |
| 437.2672 | 3.28 | PA(18:0) | C21H42O7P | 437.2674 | 0.37 |
| 701.5113 | 100 | PA(18:1/18:0) | C39H74O8P | 701.5127 | 1.97 |
| **788.5447** | **89** | **PS(18:1/18:0)** | **C42H79O10NP** | **788.5447** | **0.01** |
| 283.2426 | 1.11 | FA(22:6)-CO2 | C21H31 | 283.2431 | 1.84 |
| 327.2325 | 2.63 | FA 22:6 | C22H31O2 | 327.233 | 1.38 |
| 478.2939 | 1.64 | PC(16:1)-CH3 | C23H45O7NP | 478.2939 | 0.02 |
| **788.5244** | **42.76** | **PC(16:1/22:6)-CH3** | **C45H75O8NP** | **788.5236** | **-1.04** |

PE(18:0/22:6) [M-H]- m/z 790.5392

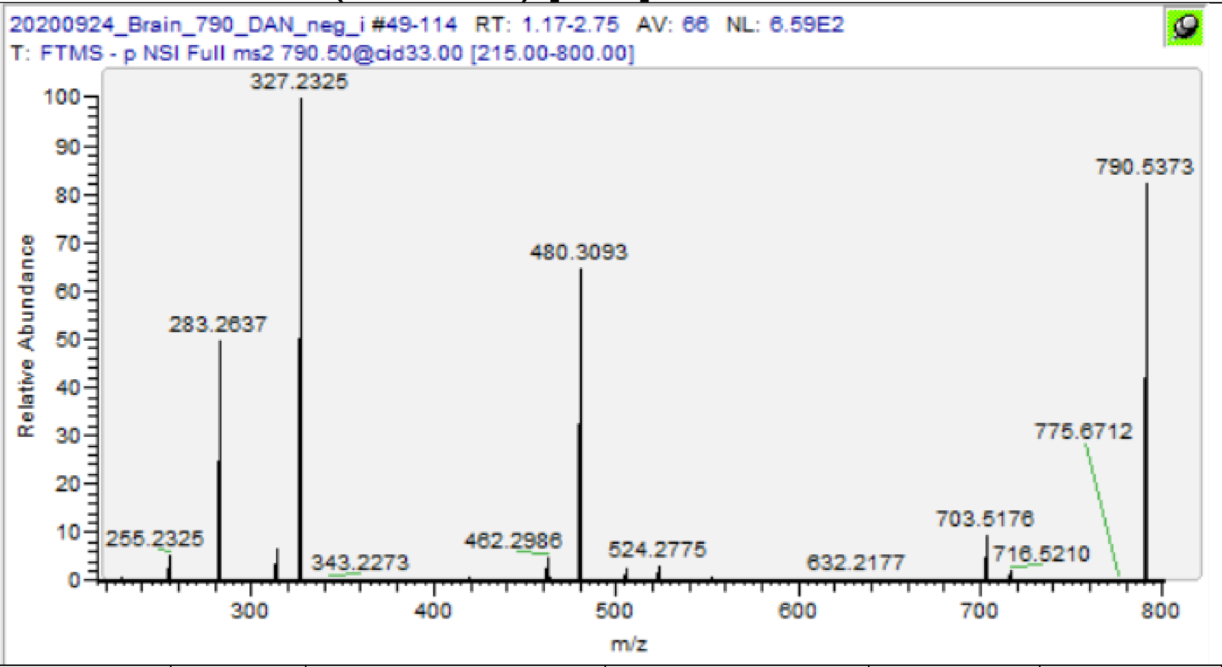

| m/z | [I] | Species | Formula | Da | ppm |
|---|---|---|---|---|---|
| 283.2426 | 29.82 | FA(22:6)-CO2 | C21H31 | 283.2431 | 1.84 |
| 283.2637 | 33.72 | FA(18:0) | C18H35O2 | 283.2643 | 1.94 |
| 327.2325 | 68.6 | FA 22:6 | C22H31O2 | 327.233 | 1.38 |
| 419.2562 | 0.36 | PA(18:0) -H2O | C21H40O6P | 419.2568 | 1.43 |
| 462.2986 | 3.17 | PE(18:0) -H2O | C23H45O6NP | 462.299 | 0.87 |
| 480.3093 | 43.06 | PE(18:0) | C23H47O7NP | 480.3096 | 0.54 |
| 506.2671 | 1.7 | PE(22:6) -H2O | C27H41O6NP | 506.2677 | 1.19 |
| 524.2776 | 2.17 | PE(22:6) | C27H43O7NP | 524.2783 | 1.26 |
| **790.5374** | **100** | **PE(18:0/22:6)** | **C45H77O8NP** | **790.5392** | **2.31** |

HexCer(d18:1/22:0(2OH)) [M-H]- m/z 798.6465

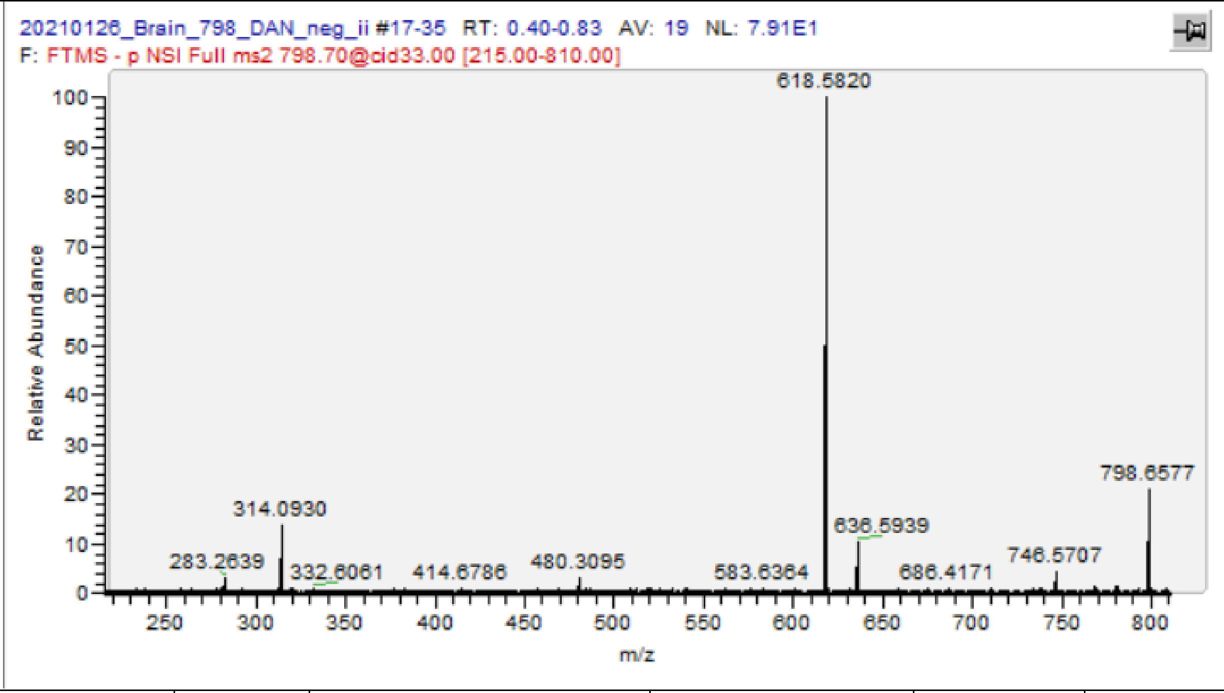

| m/z | [I] | Species | Formula | Da | ppm |
|---|---|---|---|---|---|
| 618.5819 | 100 | Cer(d40:1(2OH)) -H2O | C40H76O3N | 618.5831 | 1.89 |
| 636.594 | 10.49 | Cer(d40:1(2OH)) | C40H78O4N | 636.5936 | -0.58 |
| 780.6371 | 1.07 | HexCer(d40:1(2OH)) -H2O | C46H86O8N | 780.6359 | -1.55 |
| **798.6443** | **100** | **HexCer(d18:1/22:0(2OH))** | **C46H88O9N-** | **798.6465** | **2.70** |

C18:1-Sulf [M-H]- m/z 806.5458

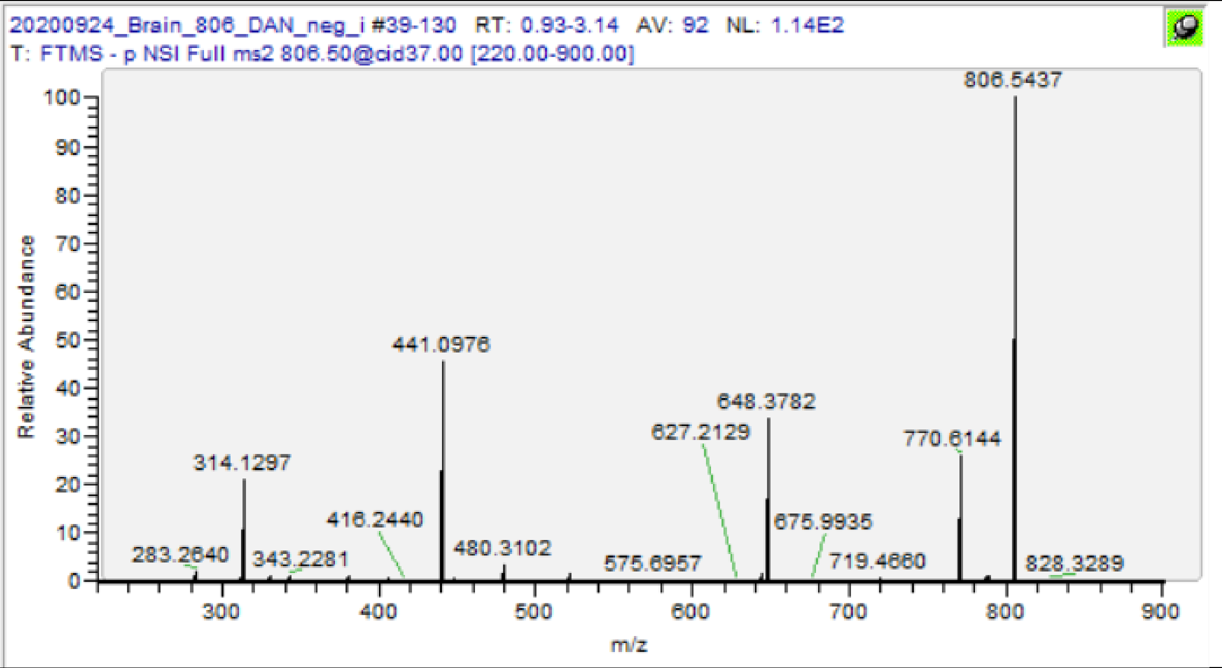

| m/z | [I] | Species | Formula | Da | ppm |
|---|---|---|---|---|---|
| 522.2748 | 1.82 | C18:1 Sulf. -C18 -H2O | C24H44O9NS | 522.2742 | -1.09 |
| 788.5358 | 0.92 | C18:1 Sulf. -H2O | C42H78O10NS | 788.5352 | -0.77 |
| **806.5436** | **100** | **C18:1 Sulf** | **C42H80O11NS** | **806.5458** | **2.68** |

C18(OH)-Sulf [M-H]- m/z 822.5407

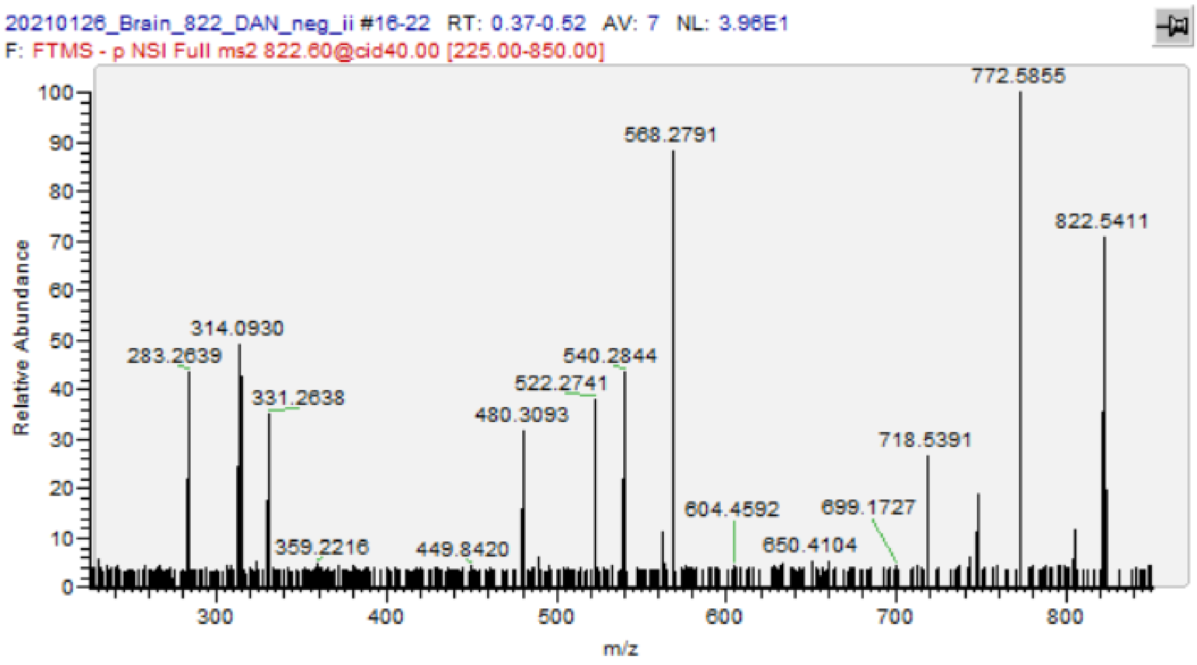

| m/z | [I] | Species | Formula | Da | ppm |
|---|---|---|---|---|---|
| 522.2741 | 38.2 | C18(OH) Sulf. -C18 -H2O | C24H44O9NS | 522.2742 | 0.25 |
| 540.2844 | 45.03 | C18(OH) Sulf. -C18 | C24H46O10NS | 540.2848 | 0.72 |
| 568.2791 | 89.36 | C18(OH) Sulf. -C17 | C25H46O11NS | 568.2797 | 1.07 |
| **822.5411** | **70.85** | **C18(OH) Sulf** | **C42H80O12NS** | **822.5407** | **-0.52** |

HexCer(d18:1/24:0(2OH)) [M-H]- m/z 826.6778

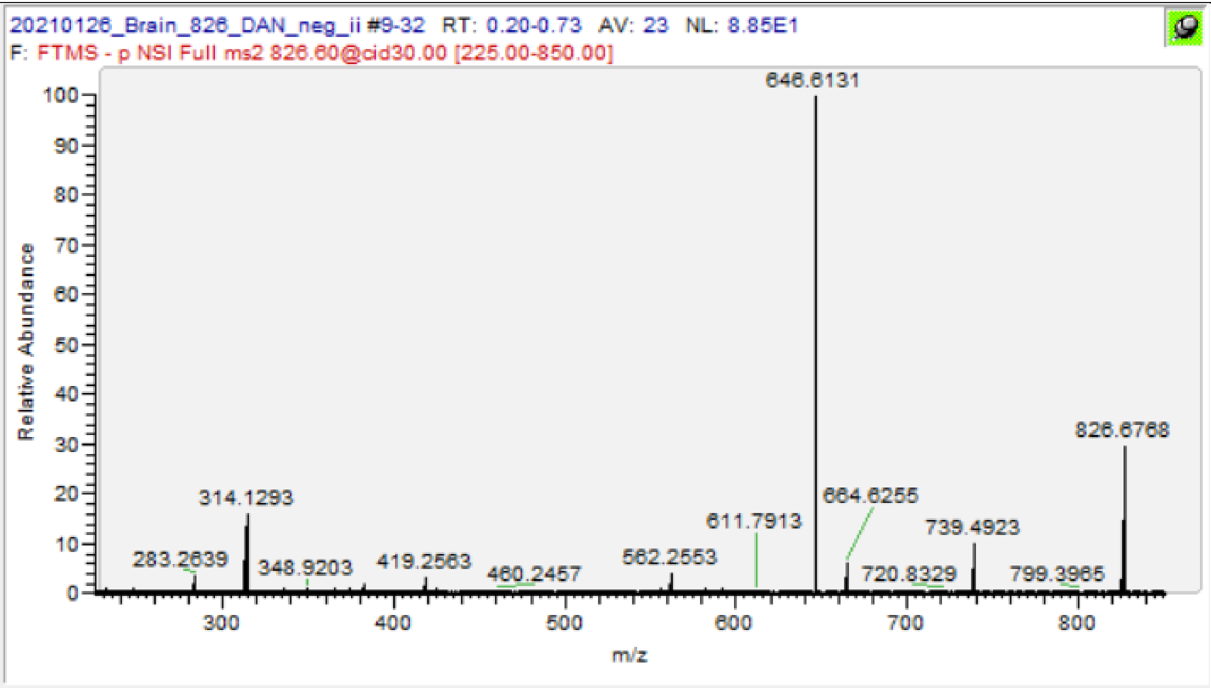

| m/z | [I] | Species | Formula | Da | ppm |
|---|---|---|---|---|---|
| 383.3528 | 2.32 | FA 24:0;O | C24H47O3 | 383.3531 | 0.70 |
| 646.613 | 100 | Cer(d18:1/24:0(2OH)) -H2O | C42H80O3N | 646.6144 | 2.12 |
| 664.6218 | 8.67 | Cer(d18:1/24:0(2OH)) | C42H82O4N | 664.6249 | 4.71 |
| **826.6757** | **59.34** | **HexCer(d18:1/24:0(2OH))** | **C48H92O9N** | **826.6778** | **2.49** |

PS(18:0/22:6) [M-H]- m/z 834.5291

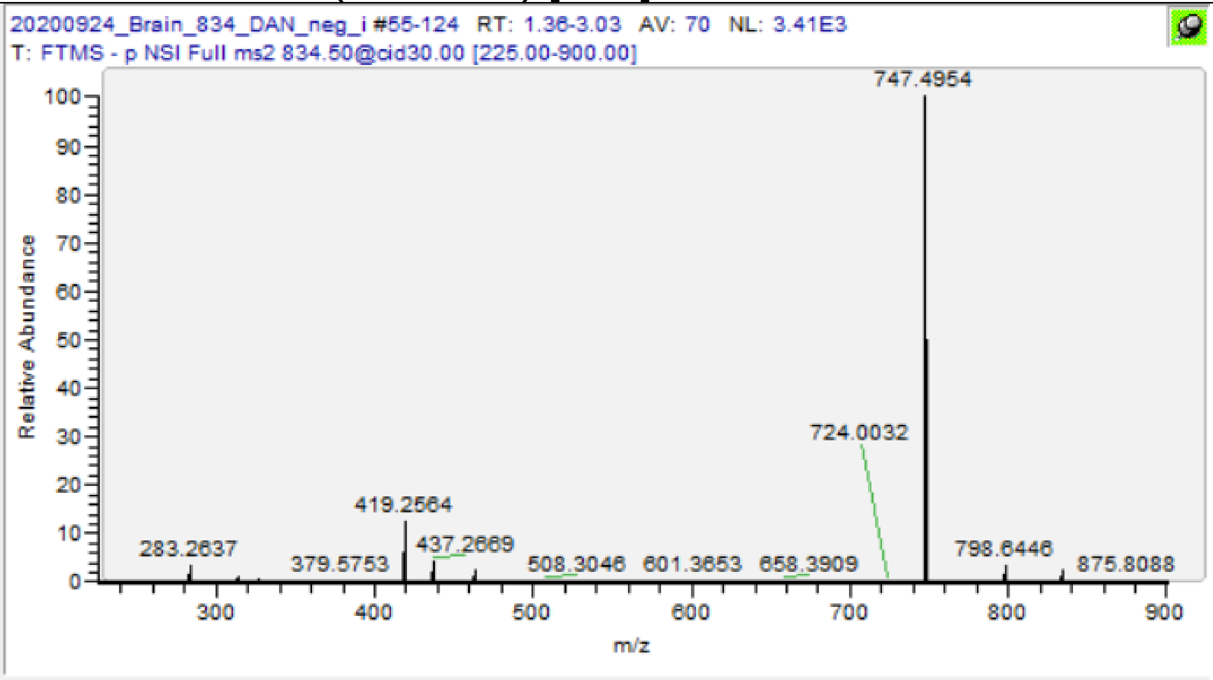

| m/z | [I] | Species | Formula | Da | ppm |
|---|---|---|---|---|---|
| 283.2427 | 0.1 | FA(22:6)-CO2 | C21H31 | 283.2431 | 1.48 |
| 283.2637 | 3.75 | FA (18:0) | C18H35O2 | 283.2643 | 1.94 |
| 327.2325 | 0.68 | FA (22:6) | C22H31O2 | 327.233 | 1.38 |
| 419.2563 | 13.12 | PA(18:0) -H2O | C21H40O6P | 419.2568 | 1.19 |
| 437.2669 | 4.51 | PA(18:0) | C21H42O7P | 437.2674 | 1.05 |
| 463.2251 | 2.75 | PA(22:6) -H2O | C25H36O6P | 463.2255 | 0.86 |
| 481.236 | 0.22 | PA(22:6) | C25H38O7P | 481.2361 | 0.12 |
| 747.4954 | 100 | PA(18:0/22:6) | C43H72O8P | 747.497 | 2.18 |
| **834.5274** | **78** | **PS(18:0/22:6)** | **C46H77O10NP** | **834.5291** | **1.99** |

PS(18:1/22:0) [M-H]- m/z 844.6073

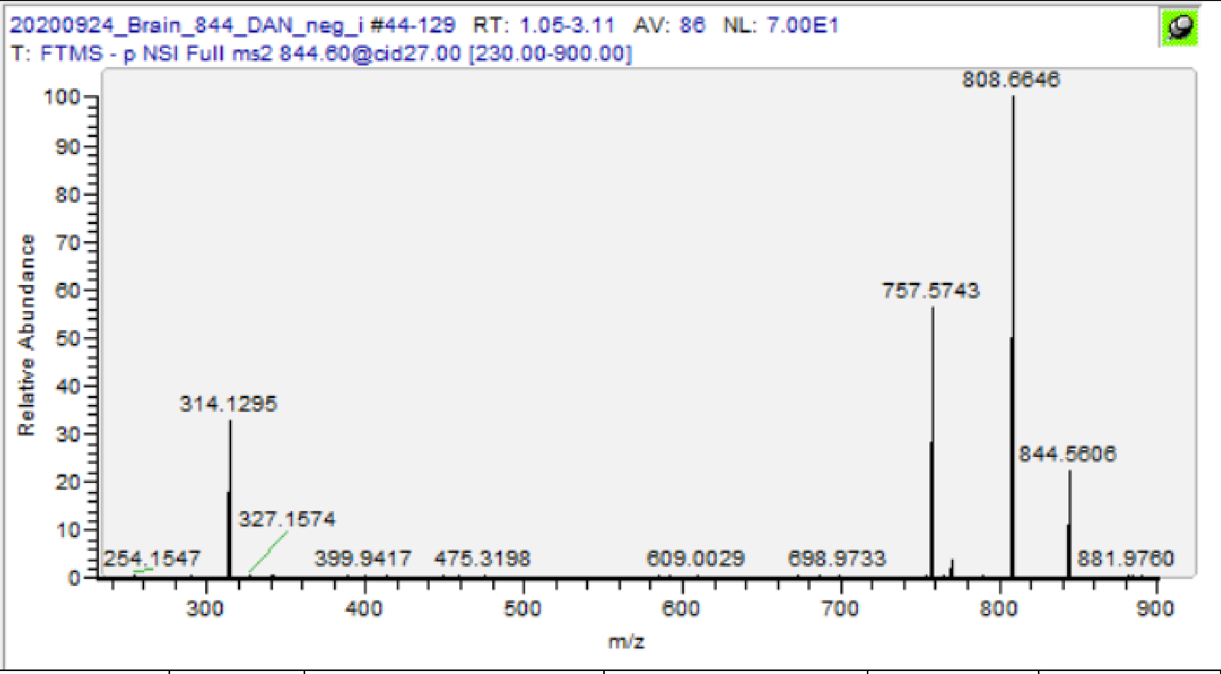

| m/z | [I] | Species | Formula | Da | ppm |
|---|---|---|---|---|---|
| 475.3198 | 0.57 | PA(22:0) -H2O | C25H48O6P | 475.3194 | -0.84 |
| 757.5743 | 56.73 | PA(18:1/22:0) | C43H82O8P | 757.5753 | 1.29 |
| **844.6081** | **10.6** | **PS(18:1/22:0)** | **C46H87O10NP** | **844.6073** | **-0.93** |

PI(16:0/20:4) [M-H]- m/z 857.5186

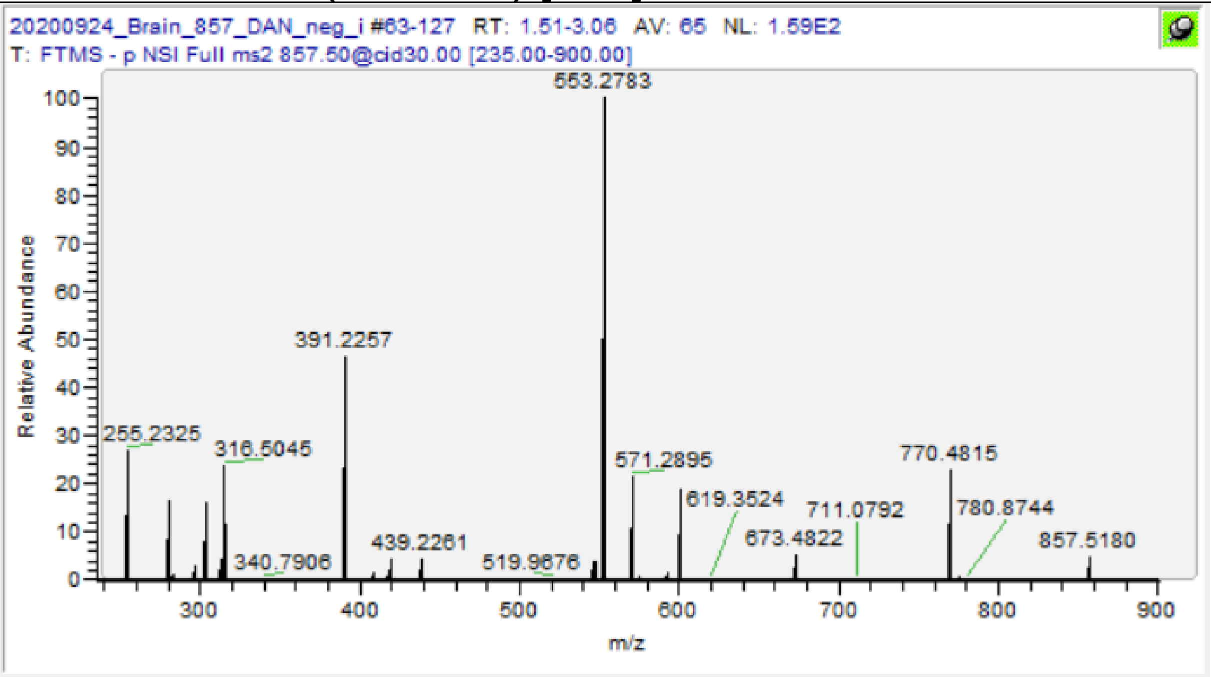

| m/z | [I] | Species | Formula | Da | ppm |
|---|---|---|---|---|---|
| 255.2325 | 26.3 | FA(16:0) | C16H31O2 | 255.233 | 1.76 |
| 303.2328 | 16.04 | FA(20:4) | C20H31O2 | 303.233 | 0.49 |
| 391.2256 | 47.15 | PA(16:0) -H2O | C19H36O6P | 391.2255 | -0.26 |
| 409.2363 | 1.67 | PA(16:0) | C19H38O7P | 409.2361 | -0.59 |
| 439.2260 | 4.16 | PA(20:4) -H2O | C23H36O6P | 439.2255 | -1.14 |
| 553.2784 | 100 | PI(16:0)-H2O | C25H46O11P | 553.2783 | -0.14 |
| 571.2896 | 21.69 | PI(16:0) | C25H48O12P | 571.2889 | -1.24 |
| 601.2789 | 18.69 | PI(20:4)-H2O | C29H46O11P | 601.2783 | -0.96 |
| **857.5180** | **4.63** | **PI(16:0/20:4)** | **C45H78O13P** | **857.5186** | **0.64** |

C22-Sulf. [M-H]- m/z 862.6084

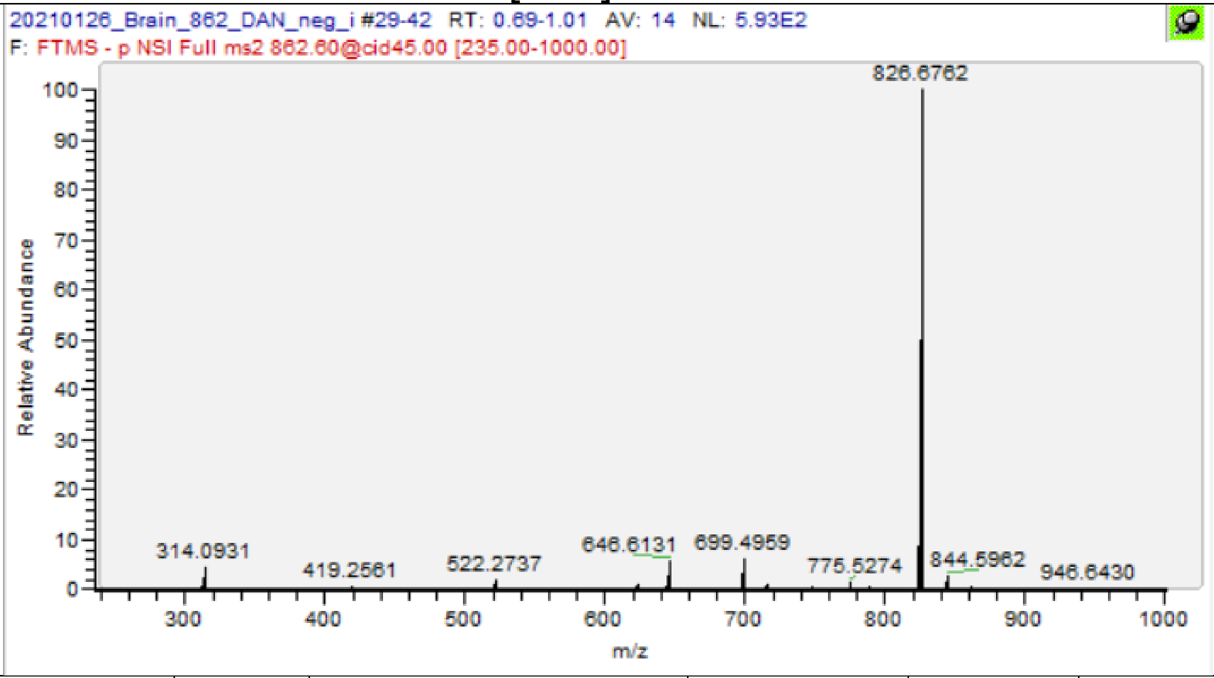

| m/z | [I] | Species | Formula | Da | ppm |
|---|---|---|---|---|---|
| 522.2736 | 1.8 | C22 Sulf. -C22 -H2O | C24H44O9NS | 522.2742 | 1.21 |
| 646.6132 | 5.69 | Cer(d18:1/24:0(2OH)) -H2O | C42H80O3N | 646.6144 | 1.81 |
| 699.496 | 5.81 | PA(18:0/18:2) | C39H72O8P | 699.497 | 1.47 |
| 826.6761 | 100 | HexCer(d18:1/24:0(2OH)) | C48H92O9N | 826.6778 | 2.01 |
| **862.6062** | **0.24** | **C22 Sulf.** | **C46H88O11NS** | **862.6084** | **2.50** |

PI(18:0/18:1) [M-H]- m/z 863.5655

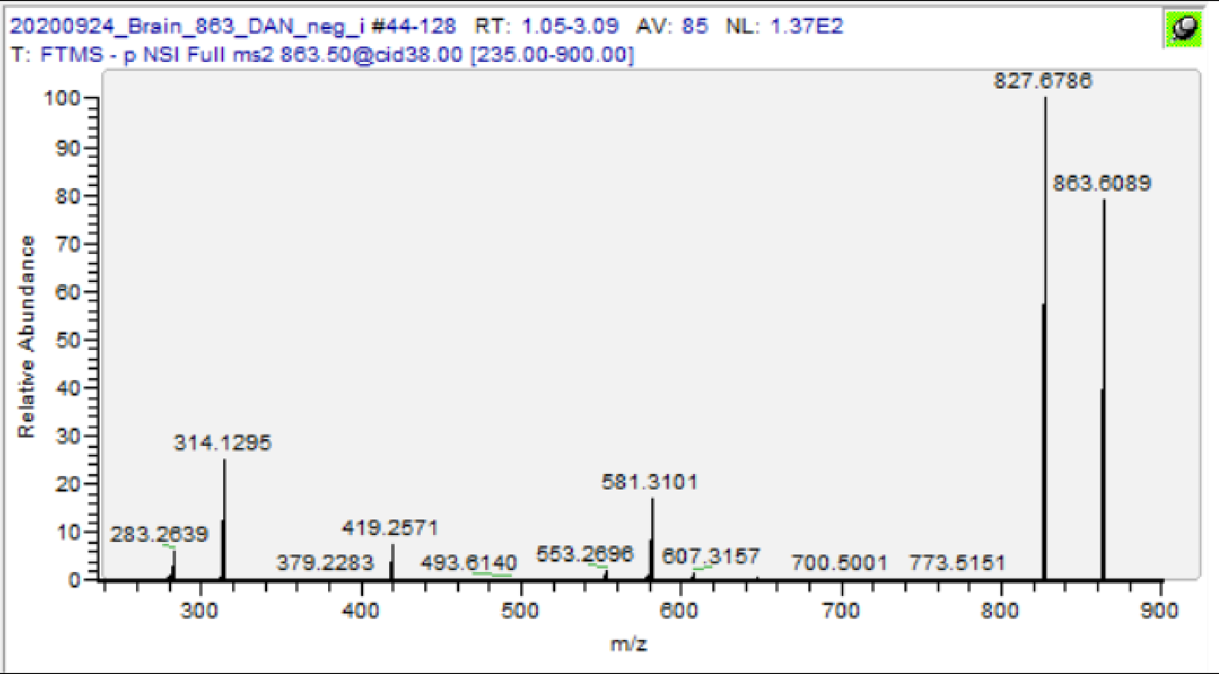

| m/z | [I] | Species | Formula | Da | ppm |
|---|---|---|---|---|---|
| 281.2484 | 1.07 | FA(18:1) | C18H33O2 | 281.2486 | 0.71 |
| 283.2639 | 6.62 | FA(18:0) | C18H35O2 | 283.2643 | 1.24 |
| 417.2421 | 0.37 | PA(18:1) -H2O | C21H38O6P | 417.2412 | -2.28 |
| 419.2572 | 7.27 | PA(18:0) -H2O | C21H40O6P | 419.2568 | -0.95 |
| 579.2947 | 1.33 | PI(18:0)-H2O | C27H48O11P | 579.294 | -1.26 |
| 581.3101 | 16.6 | PI(18:0)-H2O | C27H50O11P | 581.3096 | -0.83 |
| **863.5636** | **56** | **PI(18:0/18:1)** | **C45H84O13P** | **863.5655** | **2.20** |

# PS(18:1/24:0) [M-H]- m/z 872.6386

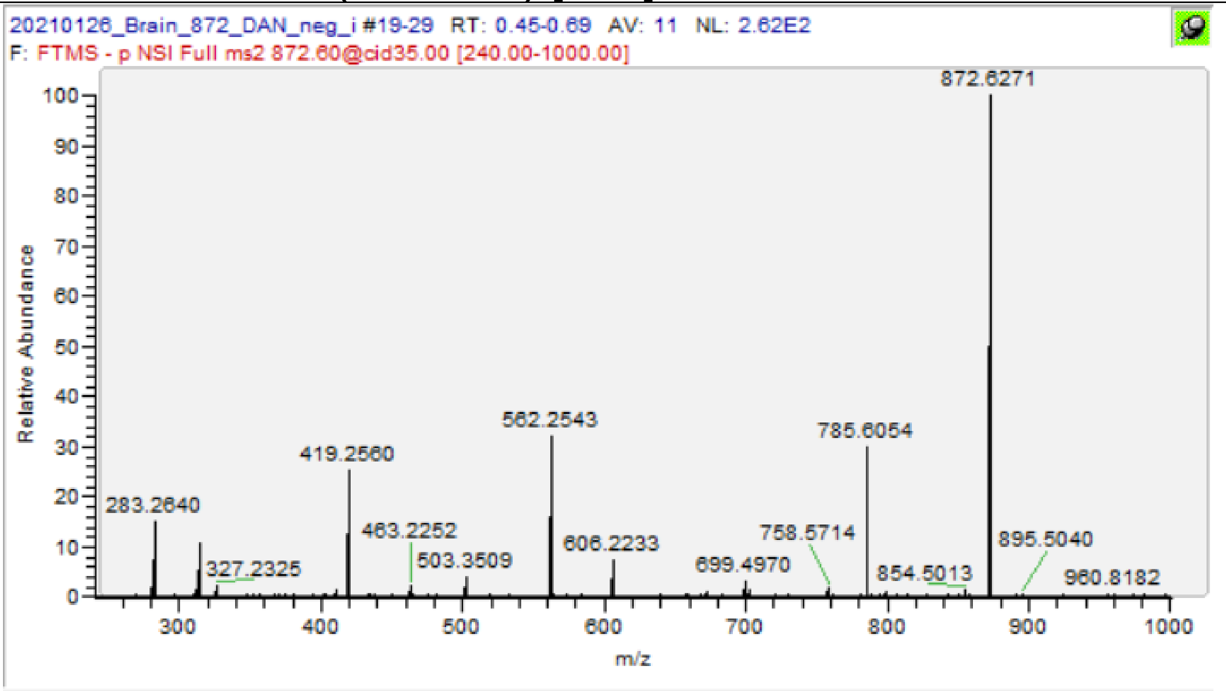

| m/z | [I] | Species | Formula | Da | ppm |
|---|---|---|---|---|---|
| 281.2482 | 7.12 | FA 18:1 | $C_{18}H_{33}O_2$ | 281.2486 | 1.42 |
| 283.264 | 32.11 | FA 18:0 | $C_{18}H_{35}O_2$ | 283.2643 | 0.88 |
| 367.3576 | 2.78 | FA 24:0 | $C_{24}H_{47}O_2$ | 367.3582 | 1.49 |
| 785.6053 | 83.61 | PA(18:1/24:0) | $C_{45}H_{86}O_8P$ | 785.6066 | 1.62 |
| **872.6341** | **1.93** | **PS(18:1/24:0)** | **$C_{48}H_{91}O_{10}NP$** | **872.6386** | **5.12** |

#### C22(OH)-Sulf [M-H]- m/z 878.6033

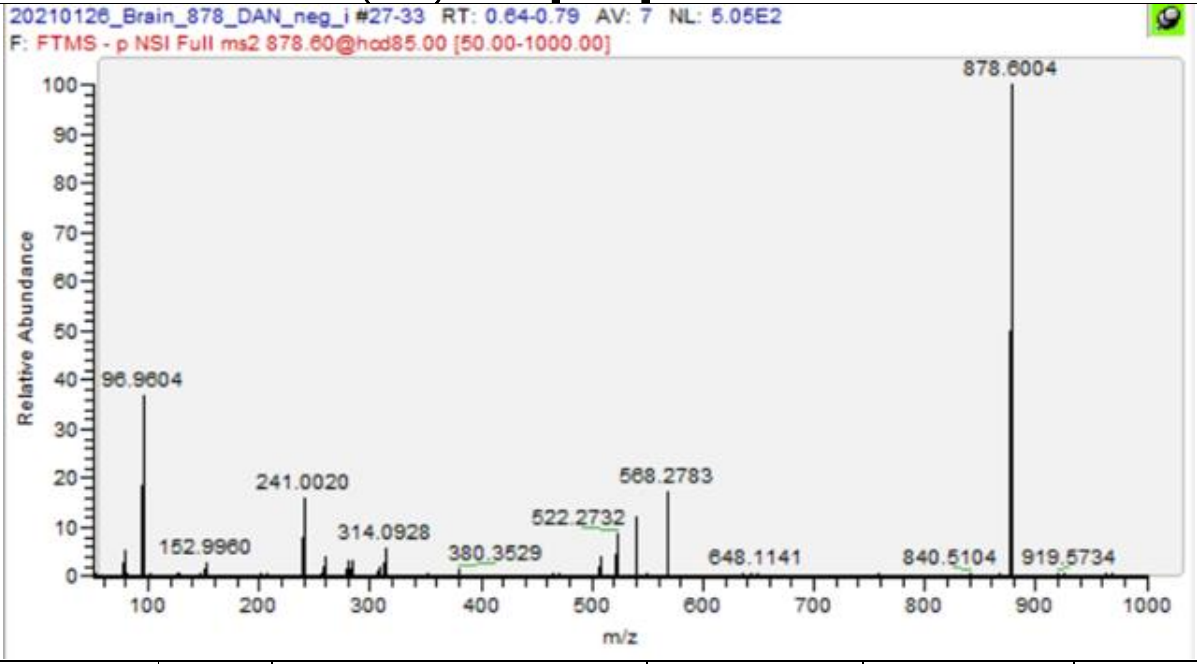

| m/z | [I] | Species | Formula | Da | ppm |
|---|---|---|---|---|---|
| 96.9604 | 32.7 | Sulfate | HO4S | 96.9601 | -3.09 |
| 241.002 | 16.54 | Sulf-Hg | C6H9O8S | 241.0024 | 1.49 |
| 507.2641 | 10.03 | C22(OH) Sulf. -C22 -N -H2O | C24H43O9S | 507.2633 | -1.52 |
| 522.2745 | 27.51 | C22(OH) Sulf. -C22 -H2O | C24H44O9NS | 522.2742 | -0.52 |
| 540.2849 | 28.72 | C22(OH) Sulf. -C22 | C24H46O10NS | 540.2848 | -0.20 |
| 550.2695 | 1.09 | C22(OH) Sulf. -C21 -H2O | C25H44O10NS | 550.2691 | -0.65 |
| 568.2793 | 81.45 | C22(OH) Sulf. -C21 | C25H46O11NS | 568.2797 | 0.72 |
| 618.5836 | 3.13 | C22(OH) Sulf. -Gala -H2O | C40H76O3N | 618.5831 | -0.86 |
| 636.5944 | 0.44 | C22(OH) Sulf. -Gala | C40H78O4N | 636.5936 | -1.21 |
| 860.5915 | 13.47 | C22(OH) Sulf. -H2O | C46H86O11NS | 860.5927 | 1.41 |
| **878.6008** | **100** | **C22(OH) Sulf** | **C46H88O12NS** | **878.6033** | **2.81** |

PI(18:1/20:4) [M-H]- m/z 883.5342

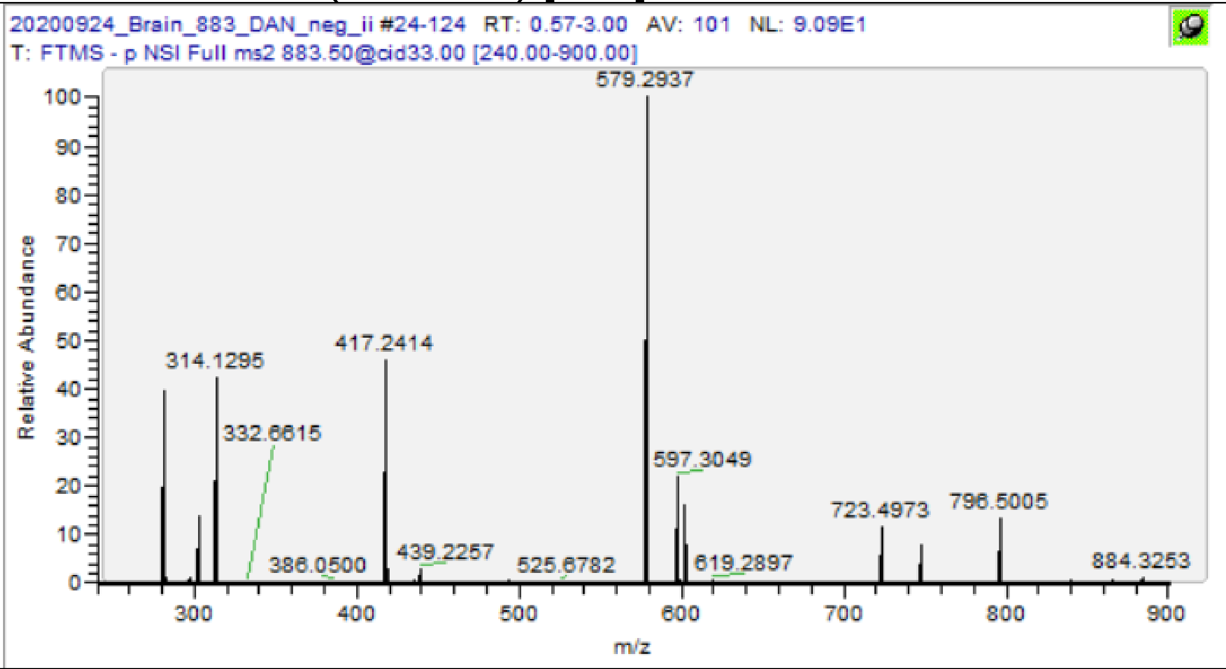

| m/z | [I] | Species | Formula | Da | ppm |
|---|---|---|---|---|---|
| 281.2482 | 39.3 | FA(18:1) | C18H33O2 | 281.2486 | 1.42 |
| 297.0378 | 1.15 | GPI -2H2O | C9H14O9P | 297.0381 | 0.98 |
| 303.2328 | 13.36 | FA(20:4) | C20H31O2 | 303.233 | 0.49 |
| 417.2414 | 45.61 | PA(18:1) -H2O | C21H38O6P | 417.2412 | -0.60 |
| 435.2524 | 0.8 | PA(18:1) | C21H40O7P | 435.2517 | -1.59 |
| 439.2257 | 2.98 | PA (20:4) -H2O | C23H36O6P | 439.2255 | -0.46 |
| 579.2937 | 100 | PI(18:0)-H2O | C27H48O11P | 579.294 | 0.47 |
| 597.3050 | 21.19 | PI(18:1) | C27H50O12P | 597.3045 | -0.77 |
| 601.2789 | 15.83 | PI(20:4)-H2O | C29H46O11P | 601.2783 | -0.96 |
| 619.2897 | 0.45 | PI(20:4) | C29H48O12P | 619.2889 | -1.31 |
| **883.5321ɣ** | **100** | **PI(18:1/20:4)** | **C47H80O13P** | **883.5342** | **2.38** |

ɣ Value from MS1 scan

PI(18:0/20:4) [M-H]- m/z 885.5499

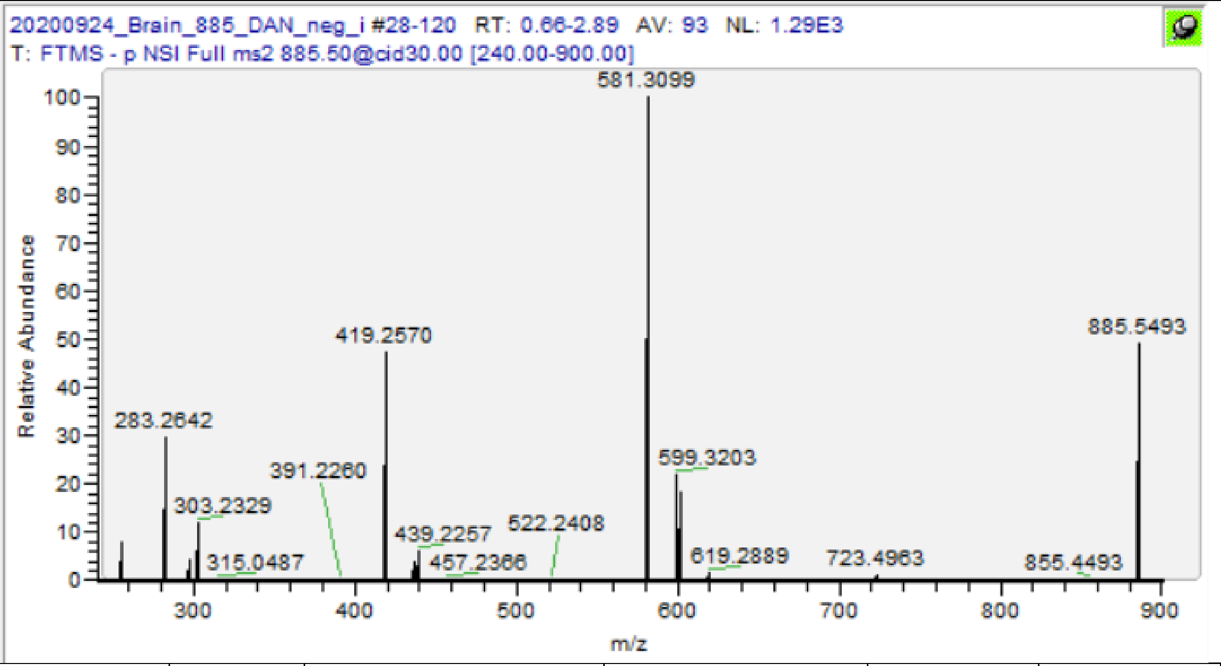

| m/z | [I] | Species | Formula | Da | ppm |
|---|---|---|---|---|---|
| 259.2431 | 0.36 | FA(20:4)-CO2 | C19H31 | 259.2431 | 0.08 |
| 283.2642 | 30.34 | FA(18:0) | C18H35O2 | 283.2643 | 0.18 |
| 297.0381 | 4.21 | GPI -2H2O | C9H14O9P | 297.0381 | -0.03 |
| 303.2329 | 12.24 | FA(20:4) | C20H31O2 | 303.233 | 0.16 |
| 315.0487 | 0.47 | GPI | C9H16O10P | 315.0487 | -0.13 |
| 419.2570 | 47.46 | PA(18:0) -H2O | C21H40O6P | 419.2568 | -0.48 |
| 437.2676 | 3.89 | PA(18:0) | C21H42O7P | 437.2674 | -0.55 |
| 439.2257 | 6.24 | PA (20:4) -H2O | C23H36O6P | 439.2255 | -0.46 |
| 581.3098 | 100 | PI(18:0)-H2O | C27H50O11P | 581.3096 | -0.31 |
| 599.3203 | 22.09 | PI(18:0) | C27H52O12P | 599.3202 | -0.18 |
| 601.2785 | 18.34 | PI(20:4)-H2O | C29H46O11P | 601.2783 | -0.30 |
| 619.2888 | 1.65 | PI (20:4) | C29H48O12P | 619.2889 | 0.15 |
| 723.4962 | 1.16 | PA(18:0/20:4) | C41H72O8P | 723.497 | 1.15 |
| **885.5493** | **48.69** | **PI(18:0/20:4)** | **C47H82O13P** | **885.5499** | **0.62** |

C24:1-Sulf. [M-H]- m/z 888.624

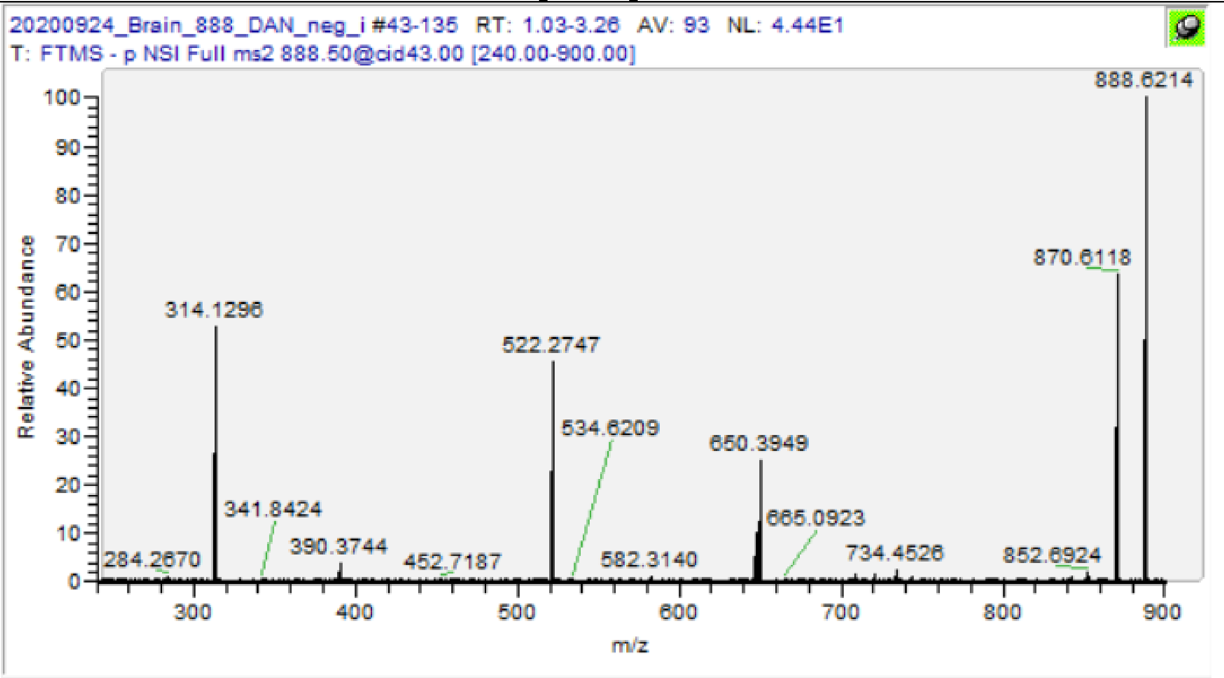

| m/z | [I] | Species | Formula | Da | ppm |
|---|---|---|---|---|---|
| 390.3745 | 8.84 | C24:1 Sulf. -C16 -Gala | C26H48ON | 390.3741 | -0.92 |
| 522.2748 | 72.85 | C24:1 Sulf. -C24 -H2O | C24H44O9NS | 522.2742 | -1.09 |
| 616.6045 | 1.17 | C24:1 Sulf. -Hg | C41H78O2N | 616.6038 | -1.14 |
| 648.3793 | 18.76 | C24:1 Sulf. -C16 -H2 | C32H58O10NS | 648.3787 | -0.94 |
| 650.3948 | 46.85 | C24:1 Sulf. -C16 | C32H60O10NS | 650.3943 | -0.71 |
| 870.6110 | 100 | C24:1 Sulf. -H2O | C48H88O10NS | 870.6134 | 2.80 |
| **888.6228** | **52.22** | **C24:1 Sulf.** | **C48H90O11NS** | **888.624** | **1.36** |

C24-Sulf. [M-H]- m/z 890.6397

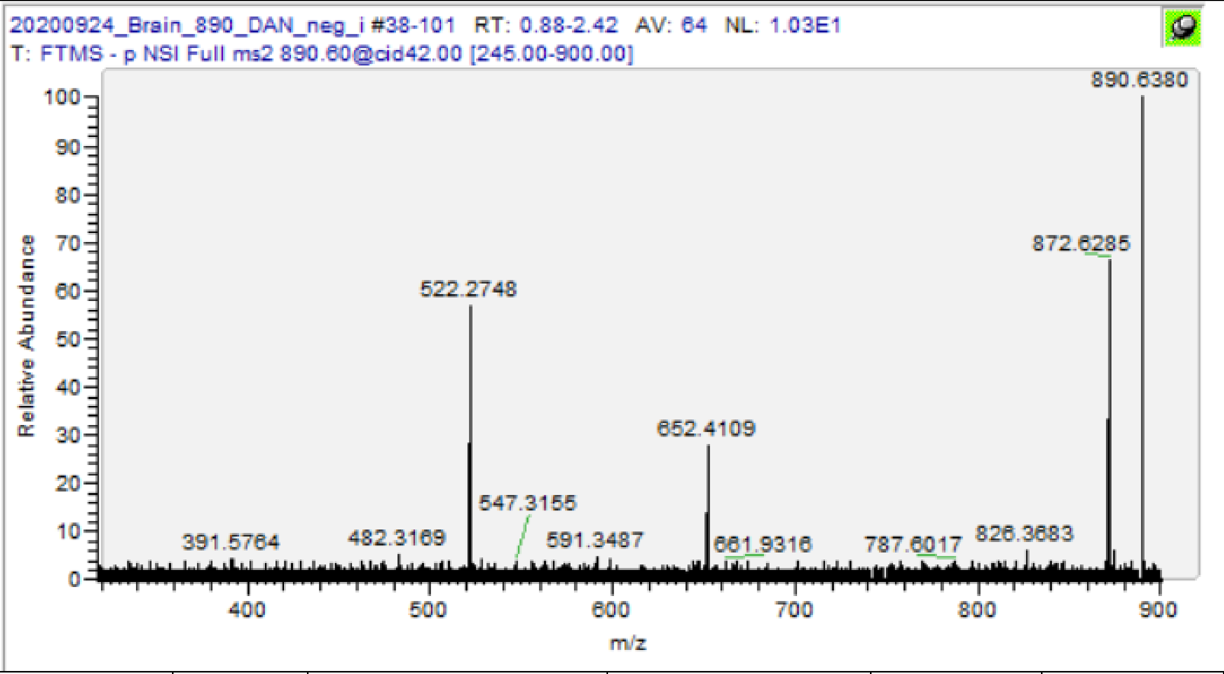

| m/z | [I] | Species | Formula | Da | ppm |
|---|---|---|---|---|---|
| 392.3891 | 0.3 | C24:0 Sulf. -C16 -Gala | C26H50ON | 392.3898 | 1.76 |
| 522.2749 | 3.16 | C24:0 Sulf. -C24 -H2O | C24H44O9NS | 522.2742 | -1.28 |
| 650.3949 | 0.47 | C24:0 Sulf. -C16 -H2 | C32H60O10NS | 650.3943 | -0.86 |
| 652.4108 | 0.85 | C24:0 Sulf. -C16 | C32H62O10NS | 652.41 | -1.24 |
| 872.6276 | 3.55 | C24 Sulf. -OH | C48H90O10NS | 872.6291 | 1.71 |
| **890.6376** | **100** | **C24 Sulf.** | **C48H92O11NS** | **890.6397** | **2.31** |

PI(18:0/20:4(OH)) [M-H]- m/z 901.5448

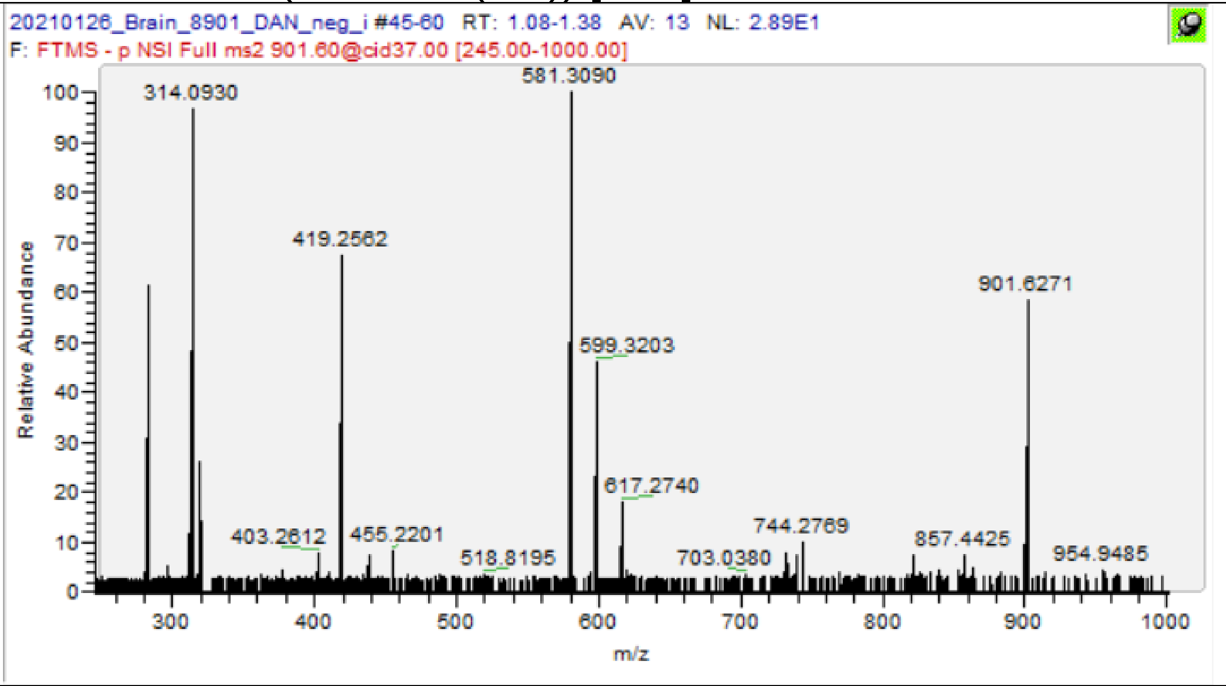

| m/z | [I] | Species | Formula | Da | ppm |
|---|---|---|---|---|---|
| 283.2638 | 5.06 | FA 18:0 | C18H35O2 | 283.2643 | 1.59 |
| 297.0378 | 5.11 | GPI -2H2O | C9H14O9P | 297.0381 | 0.98 |
| 319.2274 | 3.05 | FA 20:4(OH) -CO2 | C20H31O3 | 319.2279 | 1.47 |
| 419.2561 | 9.2 | PA(18:0) -H2O | C21H40O6P | 419.2568 | 1.67 |
| 437.2666 | 0.84 | PA(18:0) | C21H42O7P | 437.2674 | 1.74 |
| 455.2202 | 1.33 | PA (20:4(OH)) -H2O | C23H36O7P | 455.2204 | 0.46 |
| 581.3087 | 11.58 | PI 18:0 -H2O | C27H50O11P | 581.3096 | 1.58 |
| 599.3193 | 6.07 | PI 18:0 | C27H52O12P | 599.3202 | 1.49 |
| 617.2729 | 4.8 | PI 20:4(OH) -H2O | C29H46O12P | 617.2732 | 0.55 |
| **901.5425** | **100** | **PI(18:0/20:4(OH))** | **C47H82O14P** | **901.5448** | **2.52** |

C24(OH)-Sulf. [M-H]- m/z 906.6346

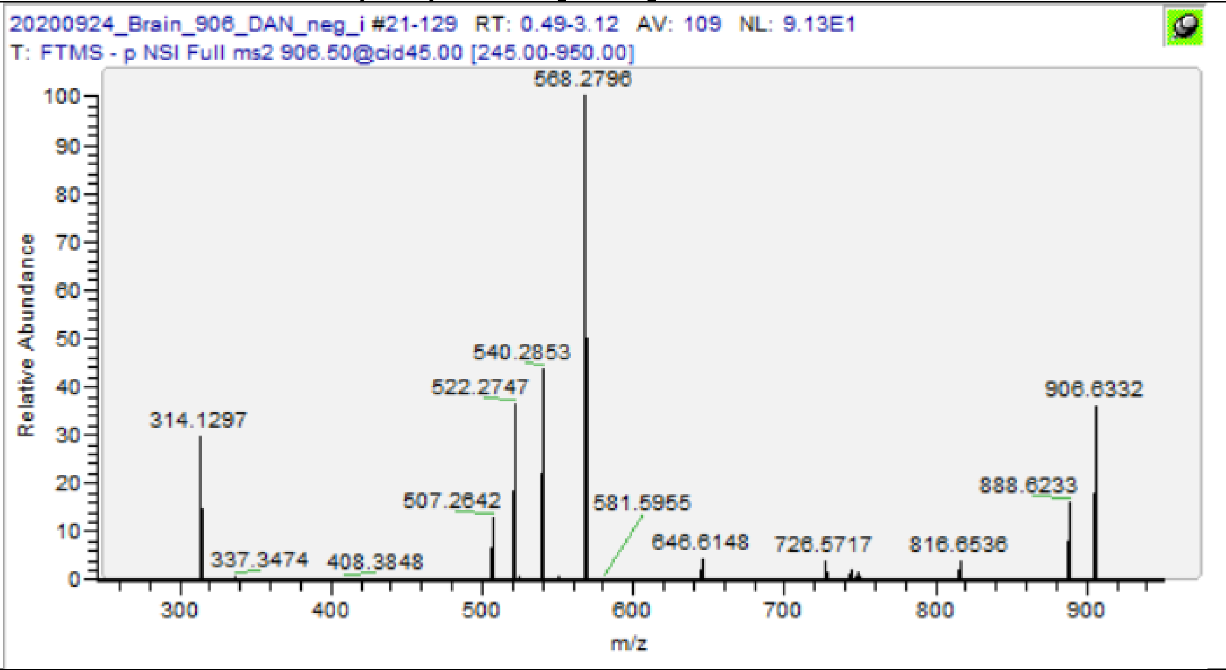

| m/z | [I] | Species | Formula | Da | ppm |
|---|---|---|---|---|---|
| 408.3848 | 0.42 | C24(OH) Sulf. -Gala | C26H50O2N | 408.3847 | -0.24 |
| 507.2642 | 13.31 | C24(OH) Sulf. -C24 -N -H2O | C24H43O9S | 507.2633 | -1.72 |
| 522.2747 | 39.17 | C24(OH) Sulf. -C24 -H2O | C24H44O9NS | 522.2742 | -0.90 |
| 540.2853 | 46.26 | C24(OH) Sulf. -C24 | C24H46O10NS | 540.2848 | -0.94 |
| 550.2698 | 0.8 | C24(OH) Sulf. -C23 -H2O | C25H44O10NS | 550.2691 | -1.20 |
| 568.2797 | 100 | C24(OH) Sulf. -C23 | C25H46O11NS | 568.2797 | 0.02 |
| 668.4034 | 0.4 | C24(OH) Sulf. -C16 | C32H62O11NS | 668.4049 | 2.26 |
| 888.6233 | 15.73 | C24(OH) Sulf. -H2O | C48H90O11NS | 888.624 | 0.80 |
| **906.6333** | **37.56** | **C24(OH) Sulf.** | **C48H92O12NS** | **906.6346** | **1.40** |

Gal-GalNAc-Gal-Glc-(d36:1) [M-H]- m/z 1254.777

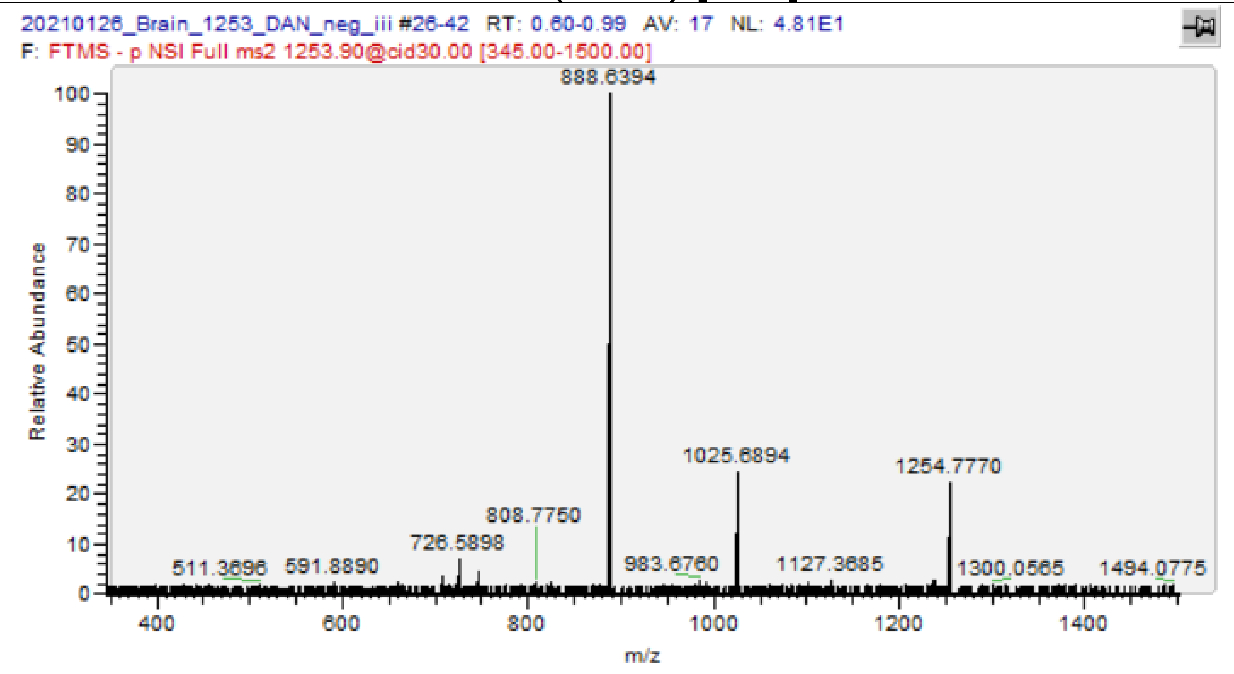

| m/z | [I] | Species | Formula | Da | ppm |
|---|---|---|---|---|---|
| 708.5791 | 3.34 | GlcCer(d36:1) -H2O | C42H78O7N | 708.5784 | -1.02 |
| 726.589 | 3.74 | GlcCer(d36:1) | C42H80O8N | 726.5889 | -0.08 |
| 888.6398 | 100 | Gal-Glc-Cer(d36:1) | C48H90O13N | 888.6418 | 2.22 |
| 983.6775 | 2.43 | GalNAc-Gal-Glc-Cer(d36:1) 0, 3 -X3 | C53H95O14N2 | 983.6789 | 1.40 |
| 1025.689 | 23.27 | GalNAc-Gal-Glc-Cer(d36:1) Z3a, Z3b - COH2 | C55H97O15N2 | 1025.689 | 0.04 |
| **1254.777** | **20.12** | **Gal-GalNAc-Gal-Glc-(d36:1)** | **C62H113O23N2 iso** | **1254.777** | **0.26** |

## GM1(d36:1) [M-H]- m/z 1544.869

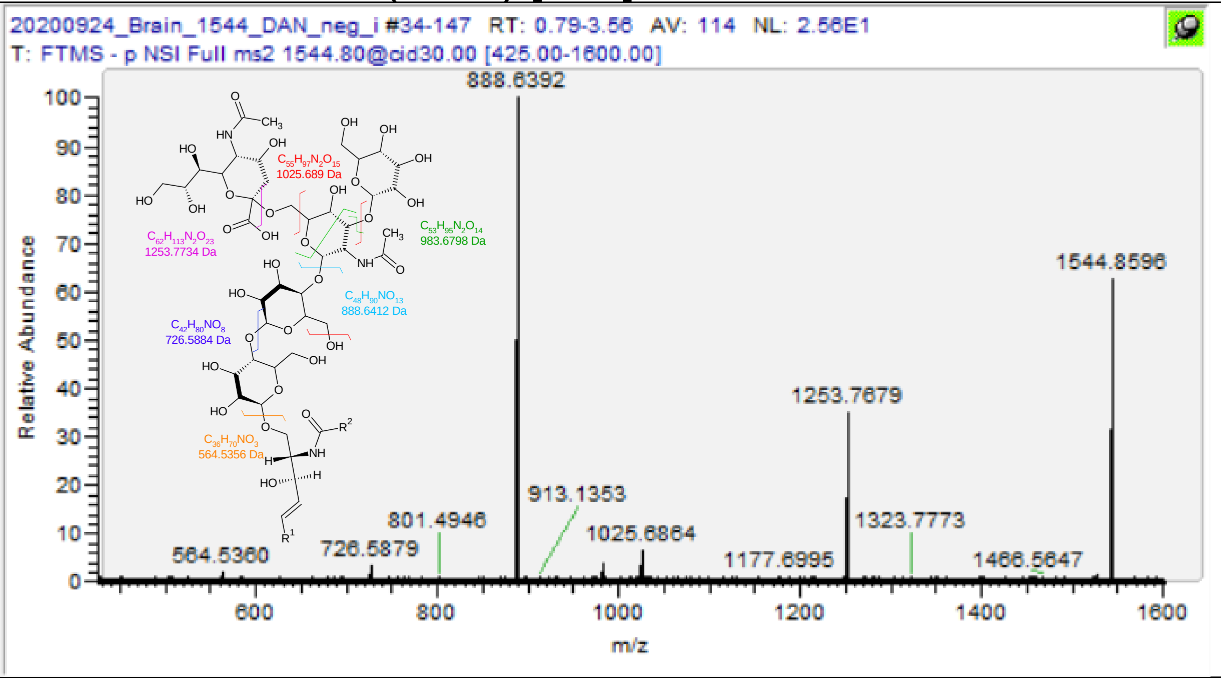

| m/z | [I] | Species | Formula | Da | ppm |
|---|---|---|---|---|---|
| 564.5360 | 1.63 | Cer(d36:1) | C36H70O3N | 564.5361 | 0.21 |
| 726.5876 | 3.12 | GlcCer(d36:1) | C42H80O8N | 726.5889 | 1.84 |
| 888.6396 | 100 | Gal-Glc-Cer(d36:1) | C48H90O13N | 888.6418 | 2.44 |
| 983.6758 | 4.56 | GalNAc-Gal-Glc-Cer(d36:1) 0, 3 -X3 | C53H95O14N2 | 983.6789 | 3.13 |
| 1025.6865 | 6.27 | GalNAc-Gal-Glc-Cer(d36:1) Z3a, Z3b - COH2 | C55H97O15N2 | 1025.689 | 2.87 |
| 1253.7679 | 36.67 | Gal-GalNAc-Gal-Glc-(d36:1) | C62H113O23N2 | 1253.774 | 4.83 |
| **1544.8632** | **66.08** | **GM1(d36:1)** | **C73H130O31N3** | **1544.869** | **4.00** |

GM1(d38:1) [M-H]- m/z 1573.904

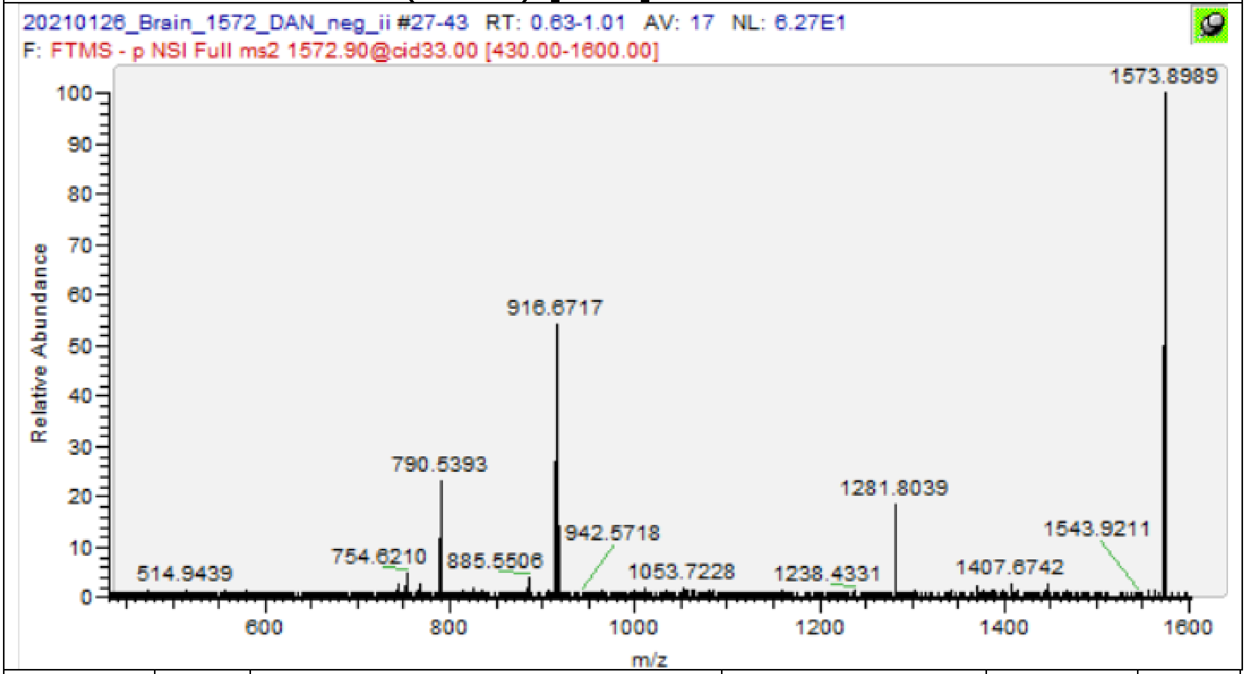

| m/z | [I] | Species | Formula | Da | ppm |
|---|---|---|---|---|---|
| 290.0882 | 34.19 | NeuAc | C11H16O8N | 290.0881 | -0.21 |
| 754.6212 | 3.02 | GlcCer(d38:1) | C44H84O8N | 754.6202 | -1.27 |
| 916.6718 | 53.53 | Gal-Glc-Cer(d38:1) | C50H94O13N | 916.6731 | 1.38 |
| 1011.71 | 1.84 | GalNAc-Gal-Glc-Cer(d38:1) 0, 3 -X3 | C55H99O14N2 | 1011.71 | 0.67 |
| 1053.723 | 2.14 | GalNAc-Gal-Glc-Cer(d38:1) Z3a, Z3b - COH2 | C57H101O15N2 | 1053.721 | -1.95 |
| 1281.804 | 19.3 | Gal-GalNAc-Gal-Glc-(d38:1) | C64H117O23N2 | 1281.805 | 1.06 |
| **1573.899** | **100** | **GM1(d38:1)** | **C75H134O31N3 iso** | **1573.904** | **3.19** |

GD1(d36:1) [M-H]- m/z 1835.9648

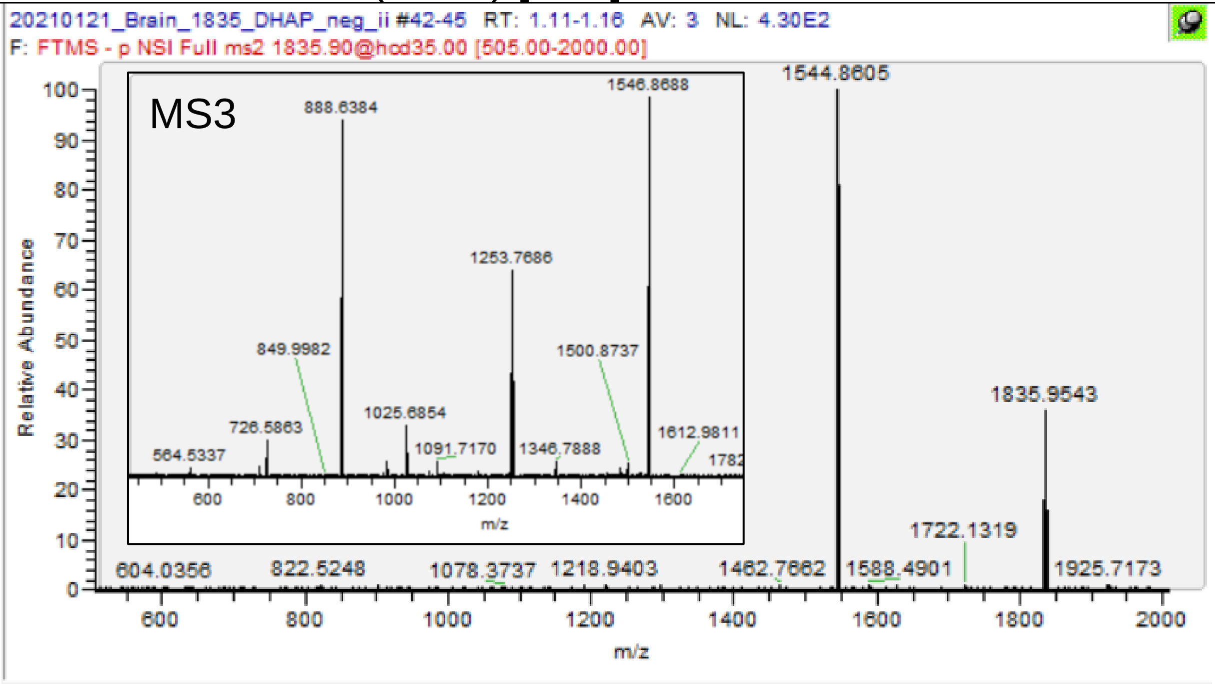

| m/ z | [I] | Species | Formula | Da | ppm |
|---|---|---|---|---|---|
| 564.5337** | 2.16 | Cer(d36:1) | C36H70O3N | 564.5361 | 4.29 |
| 708.5759** | 2.97 | GlcCer(d36:1), Z | C42H78O7N | 708.5784 | 3.50 |
| 726.5863** | 9.44 | GlcCer(d36:1), Y | C42H80O8N | 726.5889 | 3.63 |
| 888.6384** | 92.72 | Gal-Glc-Cer(d36:1) | C48H90O13N | 888.6418 | 3.79 |
| 983.6756** | 3.9 | GalNAc-Gal-Glc-Cer(d36:1) 0, 3 -X3 | C53H95O14N2 | 983.6789 | 3.33 |
| 1025.6854** | 13.74 | GalNAc-Gal-Glc-Cer(d36:1) Z3a, Z3b -COH2 | C55H97O15N2 | 1025.6894 | 3.94 |
| 1091.717** | 3.98 | GlcNAc-Gal-Glc-Cer(d36:1) | C56H103O18N2 | 1091.7211 | 3.79 |
| 1253.7686** | 54.36 | Gal-GalNAc-Gal-Glc-(d36:1) | C62H113O23N2 | 1253.7740 | 4.28 |
| 1544.8631* | 100 | GM1(d36:1) | C73H130O31N3 | 1544.8694 | 4.07 |
| **1835.957*** | **32.92** | **GD1(d36:1)** | **C84H147O39N4** | **1835.9648** | **4.24** |
| * MS2@1835.9 | | | | | |
| ** MS3@1544.8 | | | | | |

GD1(d36:1) [M-2H+K]- m/z 1873.921

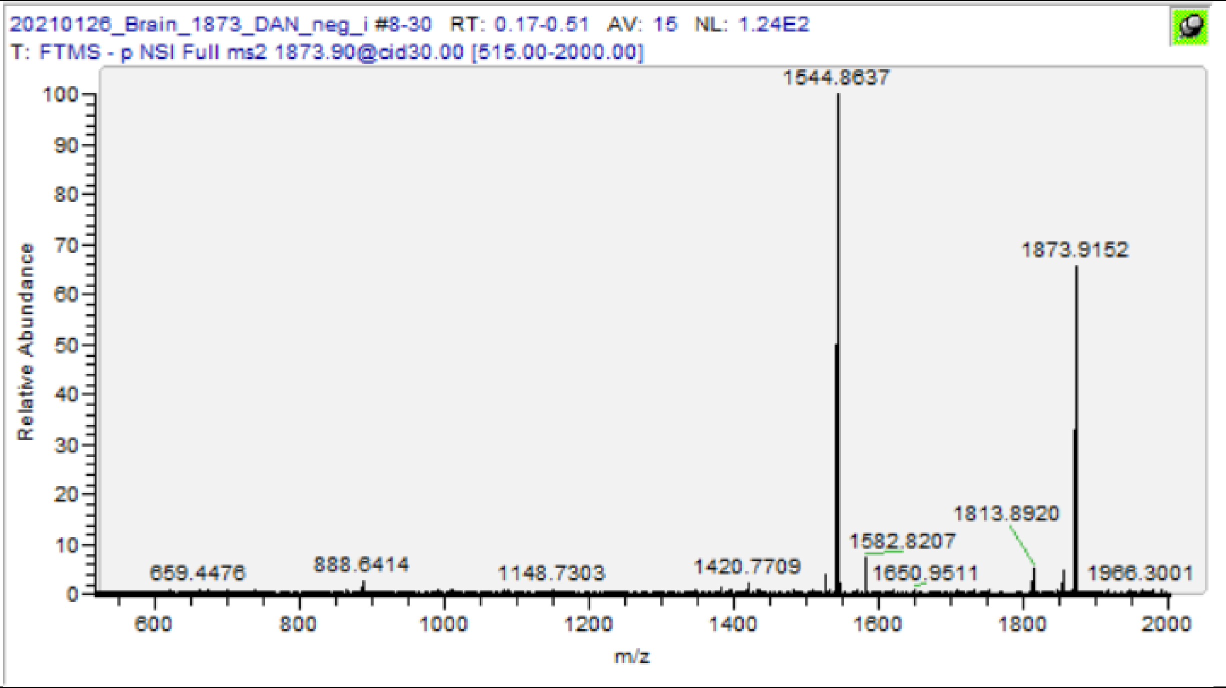

| m/z | [I] | Species | Formula | Da | ppm |
|---|---|---|---|---|---|
| 888.6414 | 2.64 | Gal-Glc-Cer(d36:1) | C48H90O13N | 888.6418 | 0.42 |
| 1253.773 | 3.99 | Gal-GalNAc-Gal-Glc-(d36:1) | C62H113O23N2 | 1253.774 | 0.85 |
| 1544.864 | 100 | GM1(d36:1) | C73H130O31N3 | 1544.869 | 3.61 |
| 1582.821 | 7.25 | GM1(d36:1)+K | C73H129O31N3K | 1582.825 | 2.88 |
| 1813.892 | 5.24 | GD1(d36:1)+K-C2H5O | C82H142O37N4K | 1813.9 | 4.16 |
| 1855.903 | 5.53 | GD1(d36:1)+K-H2O | C84H144O38N4K | 1855.91 | 3.62 |
| **1873.916** | **88.88** | **GD1(d36:1)+K** | **C84H146O39N4K** | **1873.921** | **2.71** |
